# Multidimensional telomere diversity and inheritance at individual and population scales

**DOI:** 10.64898/2026.08.19.745664

**Authors:** Huihui Li, Congying Chen, Ludong Yang, Zepu Miao, Yan Shuai, Wenjing Bao, Human Pangenome Reference Consortium, Jia-Xing Yue

## Abstract

Variation in telomere length, sequence composition and epigenetic state influences genome stability, aging and disease, yet its high-resolution characterization across species remains challenging. Here we present TeloXplorer, a computational framework for long-read data that jointly profiles telomere length, telomere variant repeats (TVRs) and DNA methylation at chromosome-end and haplotype resolution. Across simulated and empirical datasets from humans, *Arabidopsis* and yeast, TeloXplorer accurately resolved chromosome-end-specific telomere features and highlighted the importance of sample-matched, haplotype-resolved assemblies. Analysis of two human trios revealed concordant relative telomere-length profiles, predominantly Mendelian transmission of TVR haplotypes and family-conserved methylation patterns. Across 232 individuals from the Human Pangenome Reference Consortium, chromosome-end telomere-length rankings were conserved across five continental and 28 population groups. High-accuracy reads from 73 individuals further revealed elevated TVR haplotype diversity among individuals of African ancestry, together with extensive interchromosomal sharing and duplication of TVR architectures. Subtelomeric TAR1 elements were strongly associated with local DNA methylation and telomere motif diversity. Together, these analyses provide a multidimensional atlas of telomere diversity across species, chromosome ends, haplotypes and populations, revealing how telomere architecture varies and is inherited across biological scales.

## Introduction

Telomeres are functionally conserved across eukaryotes, capping and protecting chromosome ends from erosion and end-to-end fusion. In most eukaryotes, telomere DNA comprises short tandem repeats specified by the telomerase RNA template^1,2^. Human telomeres typically span 5–15 kb and consist predominantly of tandem copies of the 6-bp TTAGGG repeat^3,4^. Telomeres in the plant *Arabidopsis thaliana* contain the 7-bp TTTAGGG repeat and range from 1 to 12 kb in length^5–7^. Despite the predominance of these canonical repeats, a heterogeneous mix of telomere variant repeats (TVRs) frequently occurs in multiple organisms, including human and *Arabidopsis*^8–10^. In comparison, telomeres in the yeast *Saccharomyces cerevisiae* are considerably shorter, typically approximately 300 bp, and comprise intrinsically heterogeneous TG_1-3_ repeats^11,12^. Telomere shortening and dysfunction compromise genome integrity and contribute to cellular senescence, organismal aging and tumorigenesis. Yet the repetitive and heterogeneous architecture has hindered high-resolution characterization of their diversity and dynamics.

Understanding this variation requires resolution at individual chromosome ends. Telomere lengths are distributed non-uniformly across the genome, indicating that chromosome-end identity contributes to their regulation. In humans, telomeres at chromosome ends 17p, 20q, and 12p consistently rank among the shortest across studies, a pattern that appears to be established at birth and maintained during aging^13–16^. Similar non-uniform distributions of telomere lengths have also been observed in mice and yeast, potentially also predetermined in *cis* by chromosome-end-specific genomic architecture^17–20^, with those shortest ends implicated in cellular senescence and genomic instability^21–23^. Moreover, emerging evidence indicates that telomere shortening at certain chromosome-ends are associated with disease risks, including both solid and hematological cancers^24,25^. Finally, chromosome-end-specific subtelomeric repeats, including human TAR1 and yeast X/Y’ elements, can bind to telomere-associated proteins and remodel chromatin states at local chromosome ends, with additional associations with telomere length^26,19,18^. Therefore, accurate characterization of telomeres and their associated genomic contexts across different chromosome ends can be crucial for understanding how chromosome-end-specific telomere architecture drives genome instability and shape disease susceptibility.

While a few specialized experimental methods have been developed for chromosome-end-specific telomere characterization, they are often labor-intensive, offer limited chromosome-end coverage, and require substantial adaptation for non-human samples^27–29^. The recent advent of long-read sequencing technologies, combined with the availability of telomere-to-telomere (T2T) reference genomes, provides new opportunities for high-resolution telomere characterization across species. Here we present TeloXplorer, a computational framework for long-read-based, multidimensional telomere analysis that characterizes telomere length, TVR composition, and DNA methylation with chromosome-end specificity. Demonstrated with its application in human, *Arabidopsis*, and yeast, we identified widespread telomere length and TVR variation across chromosome ends, while underscoring the importance of using sample-matched, haplotype-phased genome assemblies for accurate telomere characterization. Leveraging on haplotype resolved human trios with diverse genetic backgrounds, we uncovered key principles shaping the inheritance in terms of length, TVR, and methylation patterns across generations. Finally, we analyzed multidimensional telomere diversity of 232 human individuals representing diverse continental and population ancestries from the second release of the Human Pangenome Reference Consortium (HPRC2)^30^. Our analysis revealed both genome-wide and chromosome-end-specific patterns governing telomere length, TVR architecture, and upstream subtelomeric methylation, while highlighted a major *cis*-acting role for subtelomeric repeats like TAR1 in shaping chromosome-end diversity.

## Results

### Overview of the TeloXplorer algorithm

TeloXplorer is a modular computational framework for integrated analysis of telomere length, telomere variant repeat (TVR) architecture and DNA methylation from long-read sequencing data and genome assemblies (Fig. 1). Its primary workflow identifies telomere-containing reads, assigns them to chromosome ends, delineates telomere–subtelomere boundaries while accommodating TVRs, and estimates chromosome-end-specific telomere length, including TVR-containing regions (Fig. 1a,b). Beyond telomere length, TeloXplorer further resolves TVR haplotypes and profiles DNA methylation relative to each boundary at chromosome-end and haplotype resolution (Fig. 1c,d). An assembly-only mode delineates boundaries and estimates telomere length directly from assembled chromosome ends when sequencing reads are unavailable. Conversely, a read-only mode estimates genome-wide telomere length in bulk when there is no suitable reference genome assembly. With these complementary modules and modes, TeloXplorer provides a versatile and powerful framework for high-resolution multidimensional telomere characterization, enabling advanced discoveries in telomere biology through long-read sequencing data.

**Figure 1.**
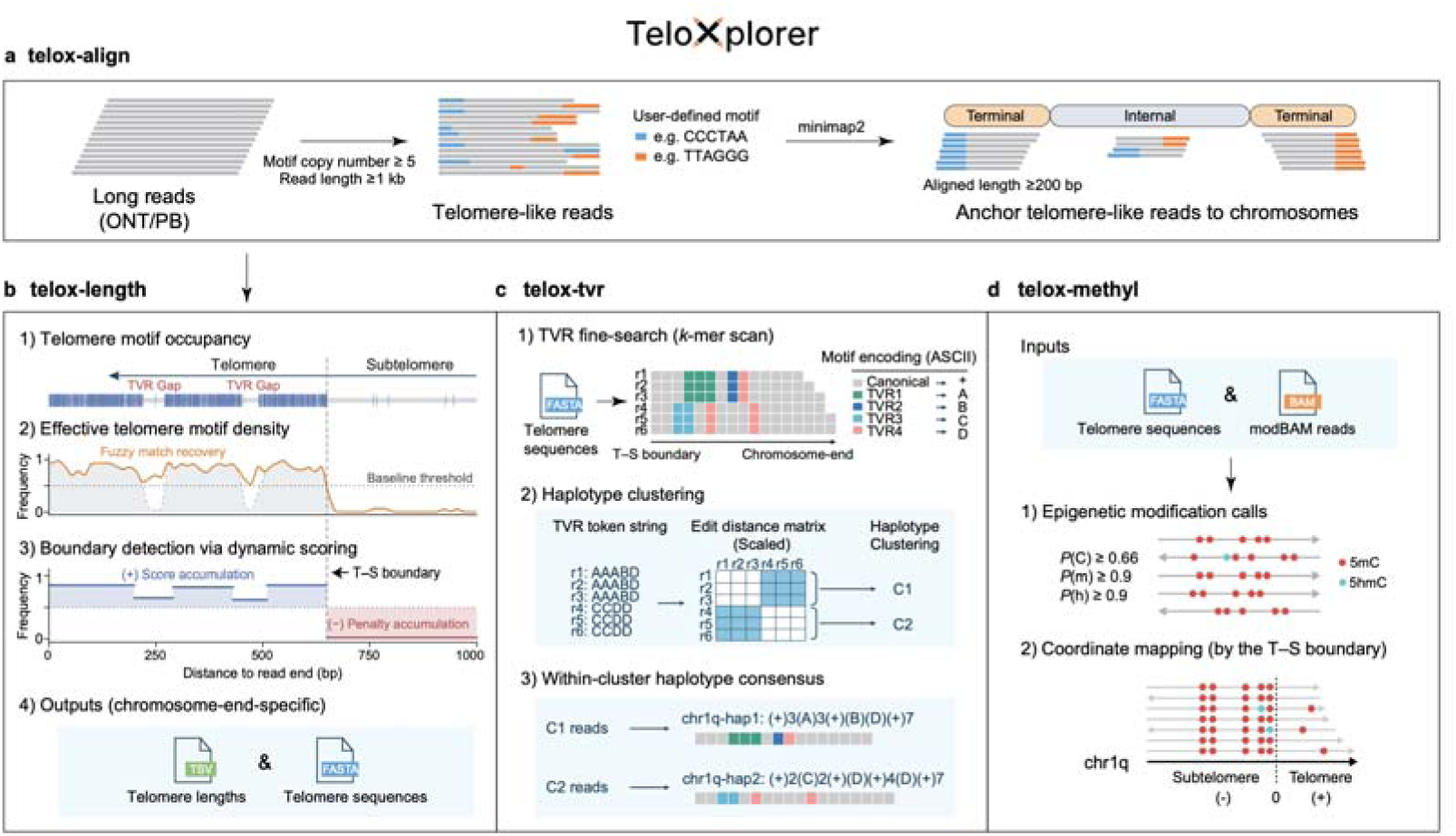
Algorithmic and functional design of TeloXplorer. **(a)** The telox-align module screens for telomere-like reads and aligns them to a native or generic reference genome assembly to identify reads of unambiguous telomeric origin. **(b)** The telox-length module pinpoints read-specific telomere–subtelomere (T–S) boundaries and summarizes chromosome-end-specific telomere lengths accordingly. **(c)** The telox-tvr module performs alignment-based TVR discovery and haplotype-aware clustering. **(d)** The telox-methyl module profiles read-specific DNA methylation signals and aligns them based on telomere–subtelomere boundaries.

### Robust telomeric region identification and characterization

Accurate delineation of telomere–subtelomere boundaries is a prerequisite for reliable telomere characterization. Using T2T or near-T2T genome assemblies from human, *Arabidopsis*, and yeast, TeloXplorer precisely identified these boundaries at all assembled chromosome-ends. This performance was maintained across species with either stable canonical repeats (humans and *Arabidopsis*) or variable repeat sequences (yeast) (Supplementary Fig. 1). Using assembly-derived telomere lengths as ground truth, we benchmarked TeloXplorer against four existing methods: TeloBP, Topsicle, Telometer, and Telogator2 (Supplementary Table 1), using simulated ONT and PacBio reads across a range of sequencing depths (5×–100×) and error rates (1%–10%). TeloXplorer, TeloBP, and Topsicle demonstrated high precision and recall for telomeric-reads identification across all tested conditions (Supplementary Fig. 2–4; Supplementary Table 2). By contrast, Telometer and Telogator2, which were applicable to primarily human datasets, showed reduced recall at low sequencing depths and high error rates, particularly Telogator2. For human datasets, all five tools produced reasonably accurate chromosome-end-specific length estimates, although Topsicle showed larger deviations (Supplementary Fig. 5). For the *Arabidopsis* and yeast datasets where only three methods are applicable, TeloXplorer consistently outperformed TeloBP and Topsicle, yielding the highest precision and narrowest error margins.

Beyond telomere length, only TeloXplorer and Telogator2 support TVR analysis, with TeloXplorer uniquely capable of leveraging the DNA methylation information embedded in long-read sequencing data for methylation analysis. Benchmarking TVR characterization on the same empirical human dataset revealed that TeloXplorer outperforms Telogator2 in revealing finer-scaled and sequence-resolved TVR landscapes (Supplementary Fig. 6).

### Integrated multi-dimensional telomere characterization

To demonstrate TeloXplorer’s capability for integrated multi-dimensional telomere characterization, we performed ONT whole genome sequencing (WGS) of the human HG002 lymphoblastoid cell line (LCL) and examined its telomeres (Fig. 2a). This cell line was selected because its fully phased T2T assembly enables haplotype-resolved analysis^31^. We found its maternal telomere lengths ranged from 0.80 kb (chr19q) to 5.67 kb (chr1q), whereas its paternal telomere lengths ranged from 0.66 kb (chr19p) to 9.61 kb (chr1p) (Fig.2b). Substantial differences between homologous chromosome ends were observed, particularly at chr1p, chr21p, chr15p, chr19p (Supplementary Fig. 7). Public HG002 datasets generated using either WGS or telomere capture (e.g., Telo-seq^32^), yielded highly concordant estimates across multiple ONT and PacBio chemistries (Supplementary Table 4). This concordance was observed at both haplotype and chromosome-end resolutions (Fig. 2b,c and Supplementary Fig. 7), demonstrating TeloXplorer’s robust telomere-length profiling across datasets and sequencing platforms.

**Figure 2.**
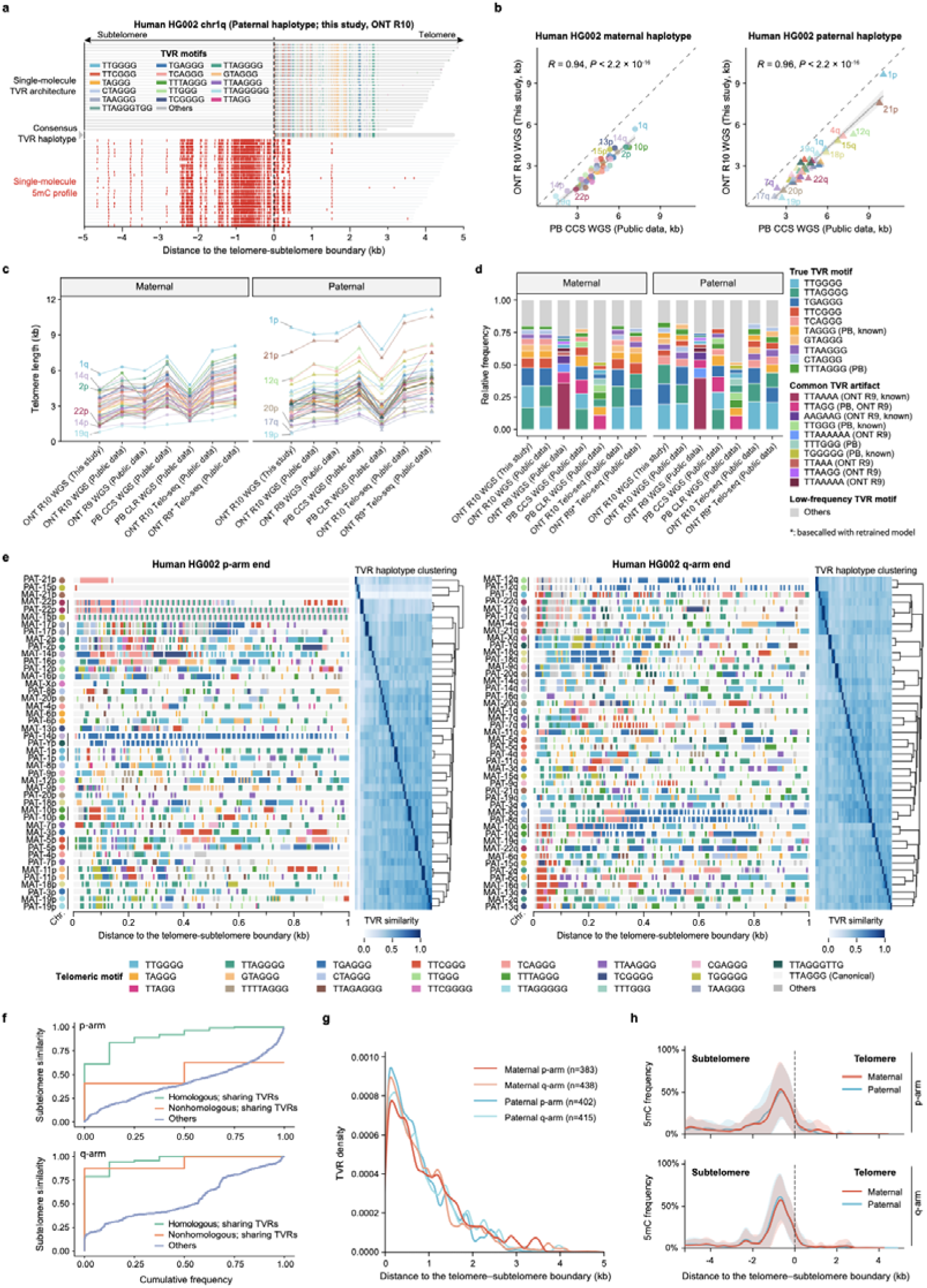
Multidimensional telomere characterization of the human HG002 cell line. **(a)** Integrated single-molecule profiling of telomere length, TVR architecture, and methylation patterns for the paternal haplotype of chromosome 1q. **(b)** Comparison of telomere lengths profiled by different sequencing platforms. **(c**,**d)** Comparison of telomere lengths (c) and TVR composition (d) profiled across multiple studies. **(e)** Landscape of haplotype-resolved, chromosome-end-specific TVR architectures. Canonical telomere repeats are shown in light gray as the background, with colored stripes representing variant repeats (TVRs). Hierarchical clustering of TVR haplotypes is shown alongside. Chromosome identities are indicated by colored circles. Chromosome ends sharing TVR signatures (TVR similarity score ≥ 0.7) within the initial 250 bp distal to the telomere–subtelomere boundary are indicated by vertical bars. **(f)** Subtelomeric sequence similarity (1 kb proximal to the telomere–subtelomere boundary) between pairing and non-pairing chromosome ends, stratified by the presence or absence of shared TVR signatures. **(g)** Spatial distribution of TVRs relative to telomere–subtelomere boundaries. **(h)** Chromosomal distribution of DNA methylation (5mC) sites relative to telomere–subtelomere boundaries.

TVR spectra were similarly concordant between WGS and Telo-seq datasets generated using ONT R10 chemistry. By contrast, datasets produced using legacy sequencing chemistries and basecalling models contained several platform-specific TVR motifs (Fig. 2d). Closer inspection indicated that these motifs were likely sequencing artifacts, several of which have been reported previously^33,34^ (Supplementary Fig. 8). Overall, these findings highlight that ONT’s R10 chemistry yields the most consistent and reliable TVR profiles.

Among the resolved TVRs, TTGGGG, TTAGGGG, TGAGGG, TTCGGG, and TCAGGG were the five most abundant motifs. Many additional TVR motifs nevertheless showed highly variable distributions both within and among chromosome ends. Notably, shared TVR architecture between maternal and paternal haplotypes were often observed on the same chromosome end, revealing chromosome-end-specific TVR signatures (Fig. 2e). Additional shared TVR patterns were found between a few non-pairing chromosome-ends (e.g., maternal chr9q and paternal chr20q), which are likely shaped by interchromosomal sequence exchanges as their immediate upstream subtelomeric regions also exhibited higher sequence similarity (Fig. 2f). Overall, TVRs were enriched near telomere–subtelomere boundaries, with the majority concentrated within the proximal 2 kb of the telomere (Fig. 2g).

TeloXplorer further used methylation calls in ONT modBAM reads to profile subtelomeric and telomeric methylation at chromosome-end and single-molecule resolution. In contrast to the heavily methylated subtelomeric regions, the 5-methylcytosine (5mC) signals are largely absent within telomeric regions, consistent with the fact that canonical (TTAGGG)_n_ repeats lack CpG sites (Fig. 2h). However, as illustrated in Fig. 2a, some TVRs introduced CpG sites and thereby allowed for DNA methylation within telomeric arrays.

### Cross-species applicability and the importance of reference choice

To assess TeloXplorer’s cross-species applicability, we extended the analysis to *Arabidopsis* and yeast, the two model organisms featuring markedly different telomere lengths and repeat architectures. Mirroring our findings in humans, TeloXplorer reported highly correlated chromosome-end-specific telomere-length estimates across different sequencing datasets (Fig. 3a,b; Supplementary Table 4). Estimates for *Arabidopsis* chr4p were unavailable because their large ribosomal DNA (rDNA) arrays remain incompletely assembled. Expanding to population-wide surveys, genome-wide median telomere lengths measured by TeloXplorer correlated strongly with estimates obtained using the conventional TRF method in both species (Supplementary Fig. 9). For yeast, this comparison used published TRF estimates adjusted for strain-specific differences in subtelomeric structure, defined by the distance between the telomere–subtelomere boundary and the proximal XhoI restriction site (Supplementary Fig. 10 and Supplementary Table 5).

**Figure 3.**
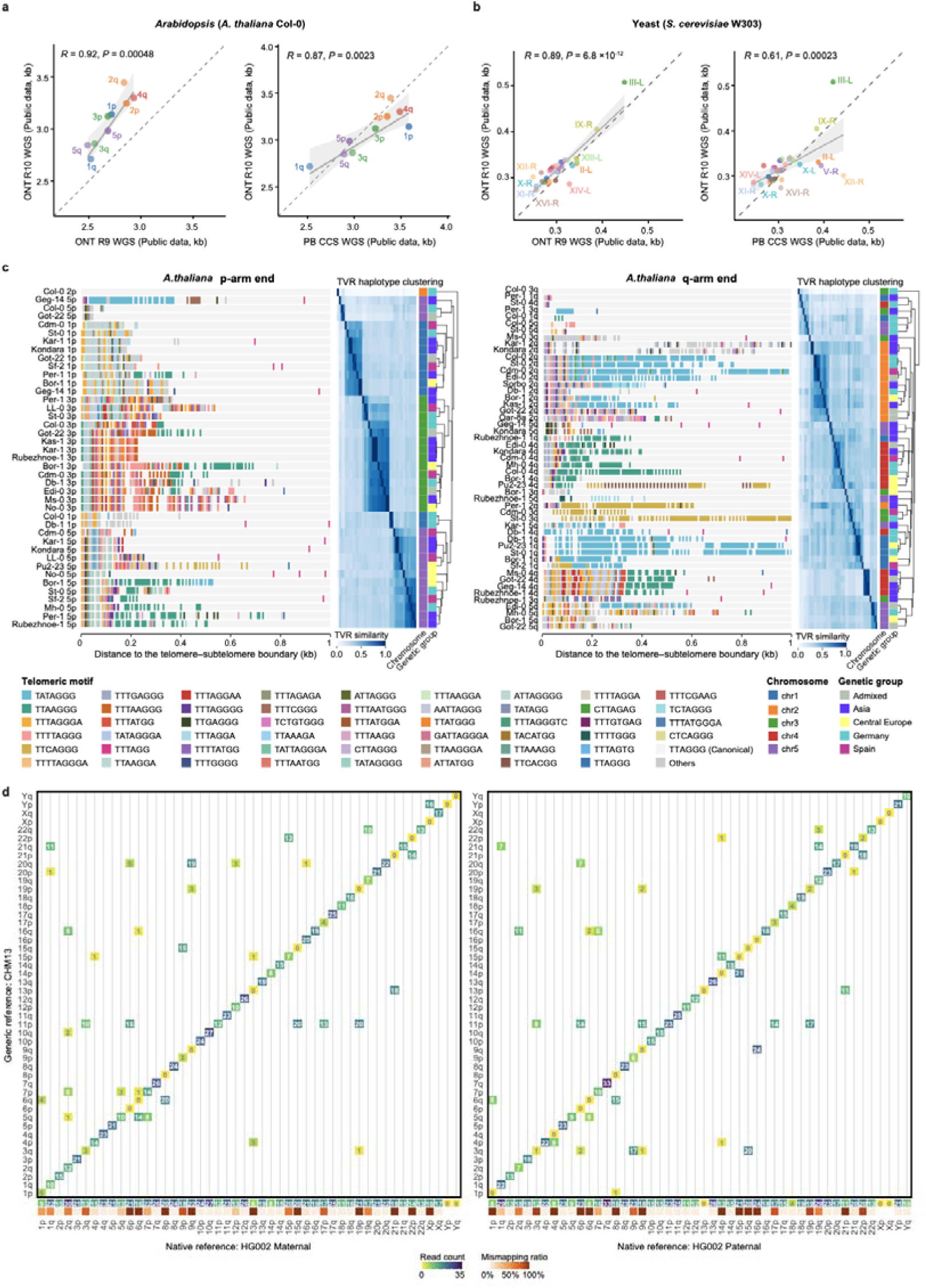
Cross-species performance and the impact of reference choice on telomere characterization. **(a)** Comparison of *Arabidopsis* (*A. thaliana* ecotype Col-0) telomere lengths profiled across different sequencing platforms and chemistries. **(b)** Comparison of yeast (*S. cerevisiae* strain W303) telomere lengths profiled across distinct sequencing platforms and chemistries. **(c)** Population-level chromosome-end-specific TVR architectures of *A. thaliana* ecotypes representing distinct genetic groups. Canonical telomere repeats are shown in light gray as the background, with colored stripes representing variant repeats (TVRs). Hierarchical clustering of TVR haplotypes is shown alongside. **(d)** Comparison of telomeric read-mapping profiles between native and generic reference genome assemblies. Taking the native HG002 alignment (maternal and paternal) as the ground truth, the plot quantifies the number of telomeric reads that map to the identical chromosome end versus alternative ends when switched to the generic CHM13 reference. The sum of read counts along each column represents the total number of reads originally mapped to that specific HG002 chromosome end, which is shown along the x-axis. The corresponding mismapping ratio for each HG002 chromosome end is further indicated underneath.

We next examined TVR variation within and among *Arabidopsis* populations. As observed for human datasets, *Arabidopsis* ONT R9 reads contained substantially more noise than ONT R10 and PB CCS reads, and were therefore excluded from subsequent TVR analyses (Supplementary Fig. 11). Comparisons across ecotypes sequenced using ONT R10 and PB CCS revealed conserved chromosome-end-specific TVR patterns (Fig. 3c and Supplementary Fig. 11). Yeast strains, by contrast, displayed highly variable TVR architectures even at the same chromosome end (Supplementary Fig. 12). No ONT-R9-specific sequencing artifacts were detected in yeast telomeres, which is consistent with a previous report^34^.

Using data from all three species, we next assessed how reference genome choice affects telomere characterization. For human HG002, switching from its native haplotype-phased assemblies to the generic CHM13 reference caused 34.94% of telomere-supported reads to be assigned to different chromosome ends, resulting in substantially altered telomere length estimates across different ends (Fig. 3d and Supplementary Fig. 13). Consistent with a previous observation^15^, acrocentric chromosome ends appear to be especially susceptible, likely due to their excessive repetitive architecture and frequent interchromosomal exchanges^35^. Similar reference-choice effects were also observed in *Arabidopsis* and yeast, although they were less pronounced (Supplementary Fig. 13). This is likely due to that these samples were less genetically diverse and lacked within-sample heterozygosity (being either haploid or homozygous diploid). Together, these findings underscore the critical importance of using sample-matched and, where applicable, haplotype-resolved assemblies for accurate telomere profiling.

### Telomere inheritance in human family trios

Understanding how parental telomeric variation shapes the telomeric landscape of children is essential for uncovering the genetic determinants of telomere maintenance. We therefore examined chromosome-end-specific inheritance of telomere length and TVR architecture, together with family-conserved DNA methylation patterns, in two human parent–offspring trios represented by LCLs (Supplementary Table 3, 4). The HG002/HG003/HG004 trio comprised the son, father and mother, respectively, and represented a family of Ashkenazi Jewish ancestry. The HG005/HG006/HG007 trio had the corresponding family structure and represented Han Chinese ancestry. The child in the latter trio had significantly longer telomeres than either parent, whereas this pattern was not observed in the former trio (Fig. 4a). This difference may partly reflect culture-associated telomere dynamics in the extensively passaged HG002 cell line, which represents the child of the former trio. Nevertheless, telomere lengths at individual chromosome ends in both children remained significantly and positively correlated with those of their respective parents (Fig. 4b). Thus, relative telomere-length profiles were preserved despite potential passage-associated shortening, consistent with a heritable component of chromosome-end-specific telomere regulation.

**Figure 4.**
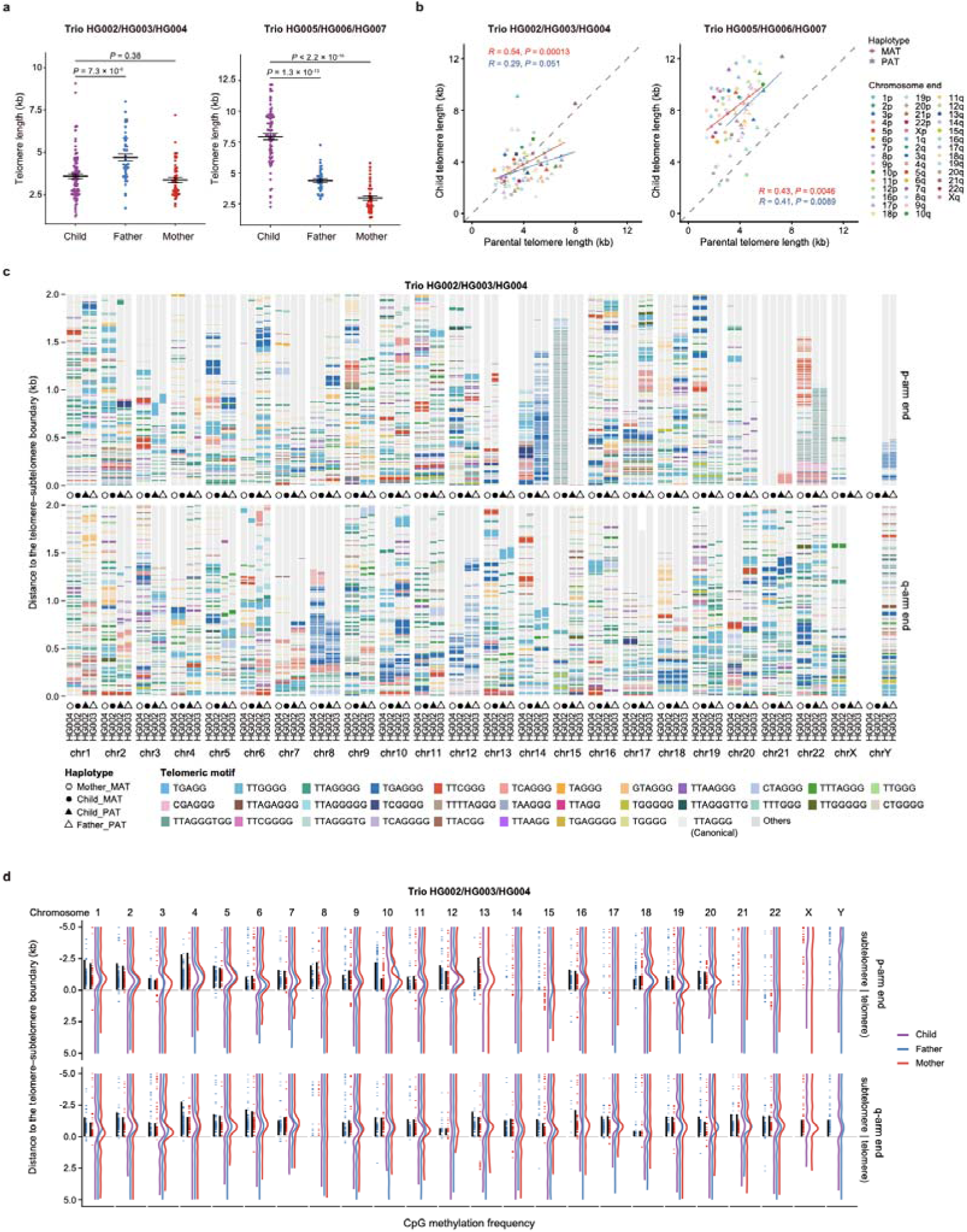
Chromosome-end-specific telomere inheritance patterns revealed across human trios. **(a)** Distribution of telomere lengths across two human trios. **(b)** Parent–child correlation of telomere lengths. **(c)** Parental origin of TVR haplotypes in the child (HG002; maternal alleles marked by filled circles, paternal alleles by filled triangles) aligned with matching haplotypes from the mother (HG004; open circles) and father (HG003; open triangles). Canonical telomere repeats are shown in light gray as the background, with colored stripes representing TVRs. **(d)** Chromosomal distribution of DNA methylation patterns across the HG002/HG003/HG004 trio (child: purple; mother: red; father: blue), shown alongside annotations for CpG clusters (blue and red stripes) and TAR1 elements (black bars) of the corresponding maternal (red) and paternal (blue) homologous chromosomes.

We next traced the transmission of TVR architectures within each family. In the HG002/HG003/HG004 trio, the child’s maternally phased architecture recapitulated that of the mother, whereas the paternally phased architecture recapitulated that of the father at every accessible chromosome end (Fig. 4c). This concordance extended beyond motif identity to their linear ordering and positional distribution across the proximal telomere. Notably, occasional child-specific TVR gains or losses were observed in more distal regions (Supplementary Fig. 14), consistent with de novo mutations in the telomeric repeat array. The HG005/HG006/HG007 trio showed the same overall parent-of-origin correspondence, with similarly rare distal deviations detected (Supplementary Fig. 15). Together, these observations support predominantly Mendelian transmission of chromosome-end- and haplotype-specific TVR architectures, extending previous findings on Mendelian inheritance of Xp/Yp telomere repeat maps^10^.

At the epigenetic level, chromosome-end-specific DNA methylation profiles were highly similar among members of each family. These profiles contained prominent local peaks immediately subtelomeric to the telomere–subtelomere boundary (Fig. 4d and Supplementary Fig. 16). These peaks are tightly linked to local CpG clusters derived from TAR1 elements. Exceptions to this pattern correspond to chromosome ends lacking TAR1 elements completely: chr14p, chr15p, chr17p, chr21p, chr22p, chrXp, chrYp, and chr8q. Other exceptions involved truncated TAR1 elements at chr12q and chr18q. Therefore, these observations identify subtelomeric TAR1 status as a major correlate of chromosome-end-specific DNA methylation profiles. Haplotype-specific TAR1 loss was also confirmed at paternal chr13p in HG002. However, no telomeric reads mapped to this chromosome end, precluding assessment of whether this haplotype-specific TAR1 loss directly affected the local methylation profile. Beyond subtelomeres, focal CpG clusters were identified within several telomeres (e.g., chr22p and the maternal chr15p haplotype), arising from CpG-containing TVRs such as TTCGGG and CGAGGG. These intratelomeric CpG clusters were also methylated, although their methylation levels were substantially lower than those of subtelomeric TAR1-bearing regions. Notably, both the strong Mendelian inheritance patterns of TVRs and TAR1-driven positional patterns of DNA methylation at chromosome ends are consistently recapitulated in the Han Chinese HG005/HG006/HG007 trio, pointing to a deeply conserved chromosome-end-specific genetic and epigenetic constraints across diverse human ancestries.

### Population-scale atlas of human telomere variation

To comprehensively map the landscape of human telomere diversity at a global scale, we extended our analysis to encompass the HPRC2 cohort of 232 individuals representing diverse genetic ancestries and geographical origins (Supplementary Table 6). Within this cohort, 159 individuals were sequenced via ONT R9 and 73 via ONT R10. We utilized all 232 individuals (464 haplotype-phased assemblies) for telomere length analysis, but restricted the TVR and methylation analyses to the 73 R10 individuals (146 haplotype-phased assemblies) to leverage their superior base-level sequence accuracy.

Telomere lengths (genome-wide median) varied substantially in the HPRC2 cohort (median: 6.79 kb, range: 1.97–21.39 kb), showing significant differences between males and females as well as among some continental groups (Supplementary Fig. 17). These findings warrant cautious interpretation due to missing donor age metadata and potential telomeric drift during LCL transformation. Nonetheless, we found no strong ALT-positive signatures (e.g., extreme inter-telomeric length heterogeneity^32^ or extensive telomere fusions^36^) in most samples (Supplementary Fig. 18), which is in line with the HPRC’s choice of using low-passage lines to mitigate cell-culture-induced genomic instability.

To assess chromosome-end-specific telomere length variation, we normalized the median telomere lengths at each chromosome end to the genome-wide median of its respective phased haplotype assembly. Despite a few outliers, the resulting chromosome-end rankings were highly consistent. Telomeres were generally shortest at chr8q, chr20q, chr17q, and chr7p, and longest at chr4p, chr3p, chr19q, and chr21p (Fig. 5a). Both the global ranking and the rankings within continental groups correlated positively with the relative mean telomere-length rankings from an independent cohort of 147 individuals (Fig. 5b and Supplementary Fig. 19). This end-specific telomere-length ordering persisted across all 28 HPRC2 populations, spanning diverse genetic backgrounds (Fig. 5c). This conservation across cohorts and populations suggests a shared genetic architecture that constrains relative telomere length at individual chromosome ends.

**Figure 5.**
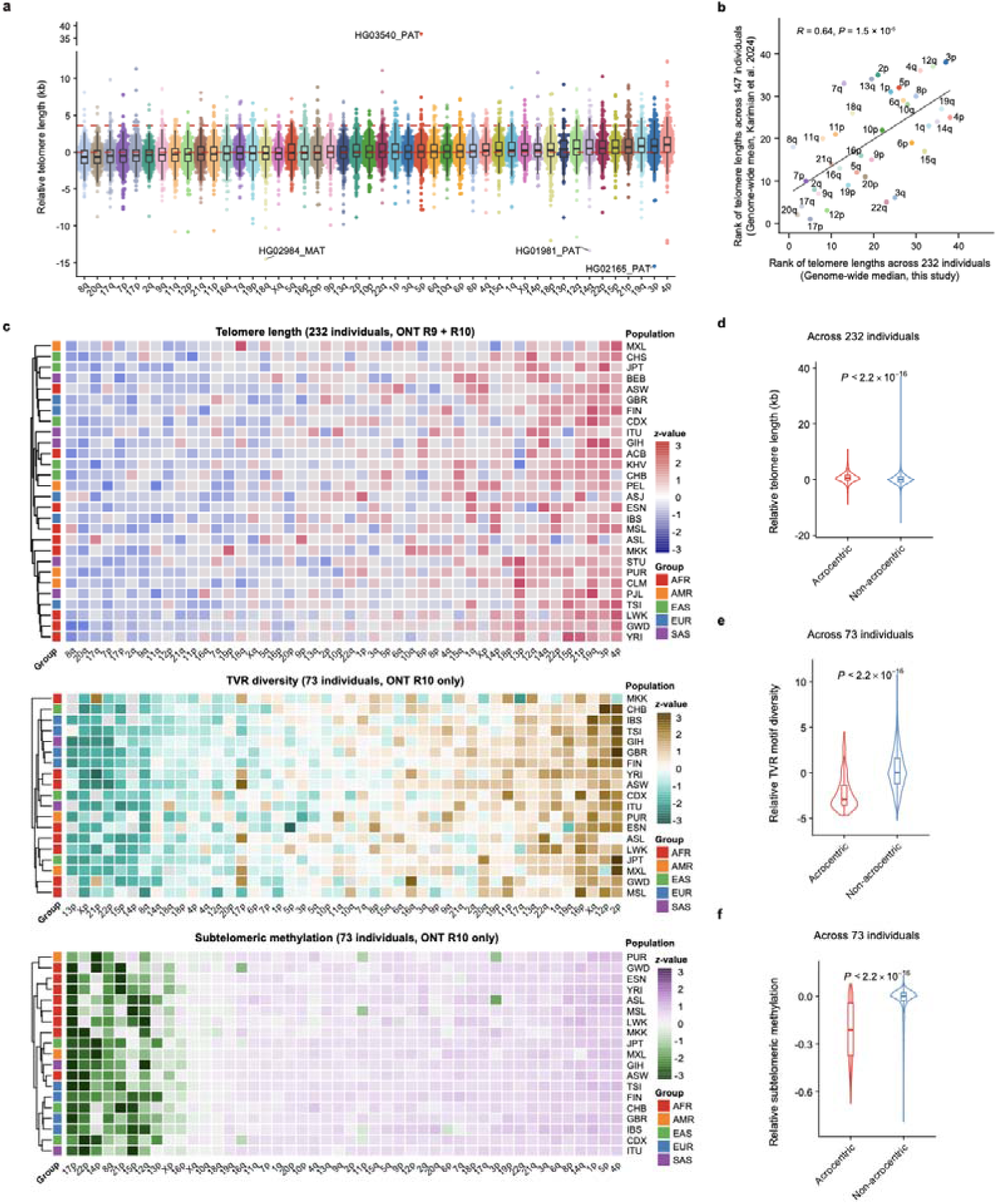
Chromosome-end-specific profiles of telomere length, TVR motif diversity, and subtelomeric methylation in the HPRC2 cohort. **(a)** Relative median telomere lengths across chromosome ends, ranked by population-wide median. Sample- and haplotype-specific outliers are indicated (PAT: paternal haplotype; MAT: maternal haplotype). **(b)** Spearman correlation of telomere-length-based chromosome-end rankings between the HPRC2 cohort and an independent published cohort. **(c)** Heatmaps of relative telomere length, TVR motif diversity (within proximal 1-kb telomeric regions), and subtelomeric methylation (within the telomere-proximal 5-kb subtelomeric regions), with chromosome ends ordered by their respective population-wide medians. **(d–f)** Comparisons between acrocentric and non-acrocentric chromosome ends for relative median telomere length **(d)**, TVR motif diversity **(e)**, and subtelomeric methylation **(f)**.

We next asked whether comparable population-wide stability extended to telomere sequence composition and subtelomeric methylation. TVR motif diversity within the first 1 kb of each telomere was quantified using a read-frequency-weighted score integrating non-canonical motif richness and the proportion of the window occupied by TVRs. Subtelomeric methylation within 5 kb of the telomere junction was measured for each chromosome end and haplotype as the proportion of methylated sites among valid CpG sites. Both relative metrics showed similarly stable chromosome-end-specific profiles across populations, although their rankings differed from each other and from that of telomere length (Fig. 5c). Specifically, TVR motif diversity was lowest at chr13p, chrXp, chr21p, and chr22p, and highest at chr2p, chr12p, chrXq, and chr16p. The TGAGGG and TCAGGG motifs were enriched, whereas TTAGGGG was depleted, at the short-arm telomeres of all five acrocentric chromosomes. The same pattern was observed at chrXp, chr8q, and chr12q as well (Supplementary Fig. 20). Subtelomeric methylation was lowest at chr17p, chr22p, chr14p, and chr8q, and highest at chr4p, chr5p, chr1p, and chr14q. Although the three metrics were not globally correlated across chromosome ends (Supplementary Fig. 21), acrocentric ends shared a distinctive profile, displaying longer telomeres alongside lower TVR motif diversity and subtelomeric methylation levels (Fig. d–f and Supplementary Fig. 22).

### Global diversity and evolutionary dynamics of TVRs

To understand how proximal telomere architecture varies across the global human population, we examined the chromosome-end-specific TVR haplotype diversity in 73 ancestrally diverse HPRC2 individuals sequenced using ONT R10. Five chromosome ends, chr2p, chr5p, chr12p, chr1q and chr7q, were each dominated by a distinct major TVR haplotype with a global frequency exceeding 50% (Fig. 6a and Supplementary Fig. 23). Six additional ends, chr1p, chr4p, chr17p, chr10q, chr13q and chr22q, had major haplotype frequencies of at least 40%. By contrast, chr18p and chr11q showed the lowest conservation, with their most frequent haplotypes accounting for only approximately 10% of observations. We detected no systematic correlation between TVR haplotypes and geographic or ancestral origins. The only exception was a specific chr11p haplotype detected exclusively in East Asian populations, specifically Chinese Dai (CDX) and Japanese (JPT), and absent in other sampled groups (Supplementary Fig. 24). Future studies featuring broader sampling will be required to elucidate its geographical distribution and evolutionary origin.

**Figure 6.**
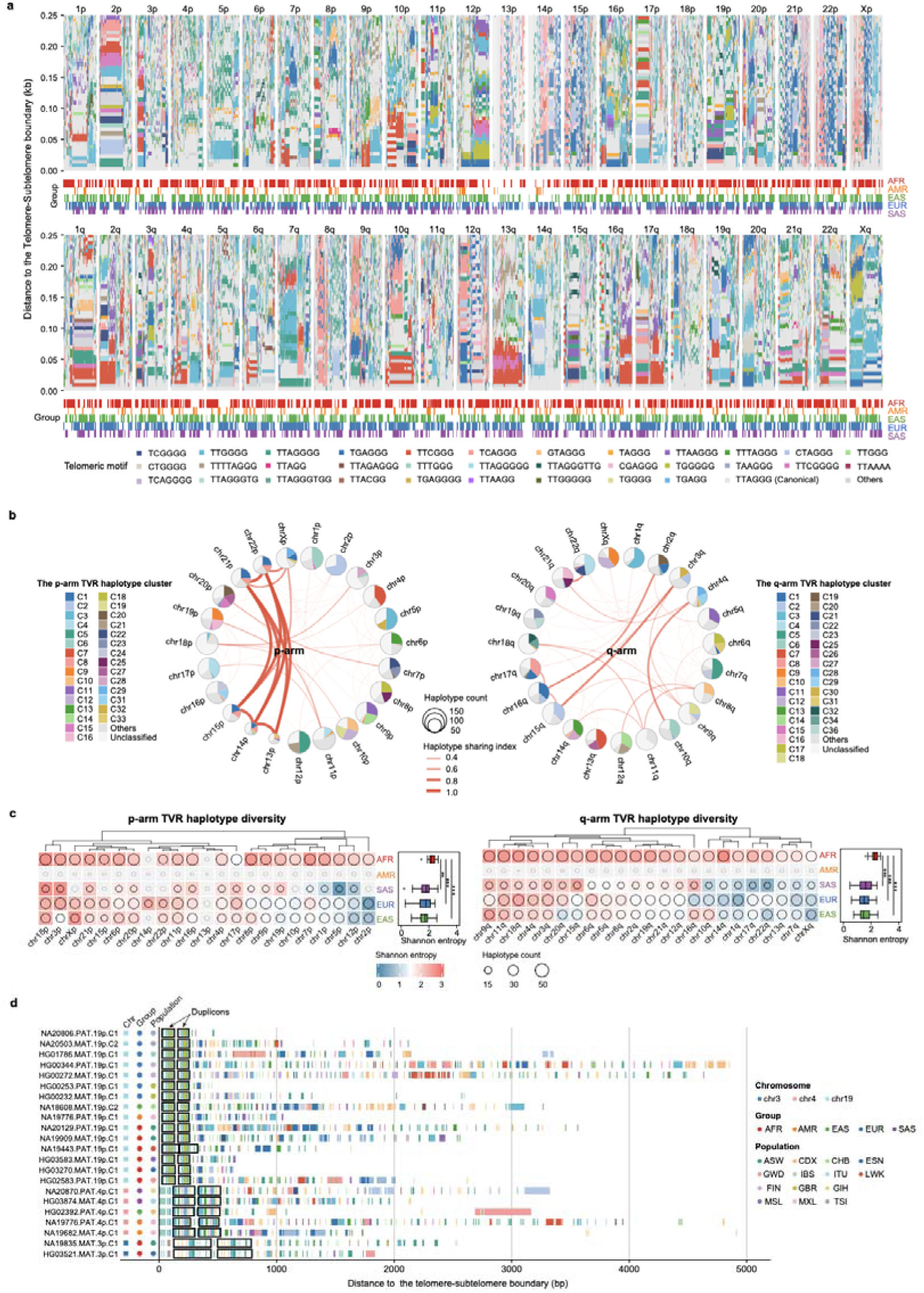
Evolutionary diversity and dynamics of TVRs across the HPRC2 cohort. **(a)** Overview of TVR haplotype diversity across the HPRC2 cohort, with canonical telomere repeats shown in light grey as the background and distinct TVR motifs color-coded. Continental group annotations for each individual are indicated below (AFR, African; AMR, Admixed American; EAS, East Asian; EUR, European; SAS, South Asian). **(b)** Population-wide composition of TVR haplotypes across individual chromosome ends. Interchromosomal similarities in haplotype composition are highlighted with connecting curved arcs. **(c)** Chromosome-end-specific TVR haplotype diversity (quantified by Shannon entropy) across continental groups, with total haplotype counts denoted by open circles. Statistical significance was evaluated using two-sided Wilcoxon signed-rank test (**: *P* < 0.05; ***: *P* < 0.01; ***: *P* < 0.001). **(d)** Examples of cross-population TVR duplicons identified at chromosome ends 19p 4p, and 3p.

Comparison across chromosome ends revealed extensive interchromosomal sharing of TVR architectures (Fig. 6b). The p arms of all five acrocentric chromosomes showed substantial haplotype similarity to one another and also shared TVR architectures with chrXp. Among q arms, prominent sharing occurred between chr2q and chr16q, chr3q and chr15q, and chr4q and chr10q. These patterns are consistent with the previously reported sequence homology among acrocentric short arms^37^, and are compatible with recurrent interchromosomal exchange through non-allelic homologous recombination or crossover.

To quantify population-level TVR haplotype diversity, we calculated Shannon entropy for each chromosome end within continental ancestry groups (Fig. 6c). The Admixed American group was excluded due to insufficient sample representation. Individuals of African ancestry exhibit significantly higher TVR haplotype diversity than individuals of South Asian, European, and East Asian ancestry. This pattern is consistent with broader patterns of human genomic diversity shaped by serial founder effects during migration out of Africa.

Finally, we identified telomeric segmental duplications comprising pairs of highly similar TVR blocks, which we termed TVR duplicons. Note that pervasive simple TVR duplication patterns exist in eight chromosome ends, including the five acrocentric short arms (13p, 14p, 15p, 21p, and 22p), as well as Xp, 8q, and 12q, which confound an accurate duplicon quantification. For the other chromosome ends, our systematic screening of the 73 individuals identified 234 TVR duplicons (Supplementary Fig. 25, 26; Supplementary Table 7). Individual duplication blocks ranged from 56 to 1132 bp, and inter-block distance was positively correlated with block size (Supplementary Fig. 27). Telomeres containing TVR duplicons were significantly longer than those without detected duplicons (two-sided Wilcoxon rank sum, *P* = 3×10^-13^). Notably, a contemporaneous analysis of overlapping HPRC2 data similarly identified recurrent internal TVR duplications and their association with longer telomere arrays^38^. This association links indicating an intrinsic link between segmental duplication and telomere elongation. It remains to be investigated whether duplication promotes elongation or preferentially accumulates in longer arrays. While the vast majority of these telomeric duplications appear to be individual- and haplotype-specific, we identified three cases (on chr3p, chr4p, and chr19p respectively) shared across multiple individuals from diverse continental groups and populations, suggesting that these telomeric duplication events are evolutionarily ancient, likely predating recent human migration events (Fig. 6d).

### *Cis*-impact of subtelomeric repeat element on adjacent telomeres

Building upon our findings that TAR1 elements dictate local DNA methylation peak near telomere–subtelomere boundaries, we investigated whether TAR1 and other subtelomeric repeat elements were associated in cis with adjacent telomere features. Using 464 HPRC2 phased human assemblies, we systematically profiled repeat content within 500 kb of each telomere–subtelomere boundary (Supplementary Fig. 28). Among all repeat classes, TAR1 (Satellite/subtelo) was positioned closest to the telomeres, with a median boundary distance of 58 bp. TAR1 was detected in 89.0% of accessible chromosome ends (18,793 of 21,106). Despite this overall prevalence, TAR1 was depleted at all five acrocentric p-arm ends and was nearly absent from chrXp and chr8q (Supplementary Fig. 29). Additional TAR1 insertions and duplications further into the subtelomere occurred at multiple chromosome ends. These included chr6p, chr8p, chr11p, chr17p, chr18p, chr19p, chr20p, chr6q, chr9q, and chr20q (Supplementary Fig. 30).

Focusing on the immediately adjacent 20-kb subtelomeric regions, the prevalence and cumulative span of different repeat classes stratified chromosome ends into distinct groups (Fig. 7a). These groupings clearly distinguished acrocentric from non-acrocentric ends, and aligned closely with recently defined subtelomere communities^39^. Principal component analysis (PCA) of repeat composition similarly separated all five acrocentric p-arm ends from the remaining chromosome ends along PC1, which explained 22.1% of the variance (Fig. 7b). Satellite/acro, rDNA, DNA-TE/hAT-Tip100, TAR1 and DNA-TE/TcMar-Mariner were among the largest contributors to PC1 and PC2 (Fig. 7c).

**Figure 7.**
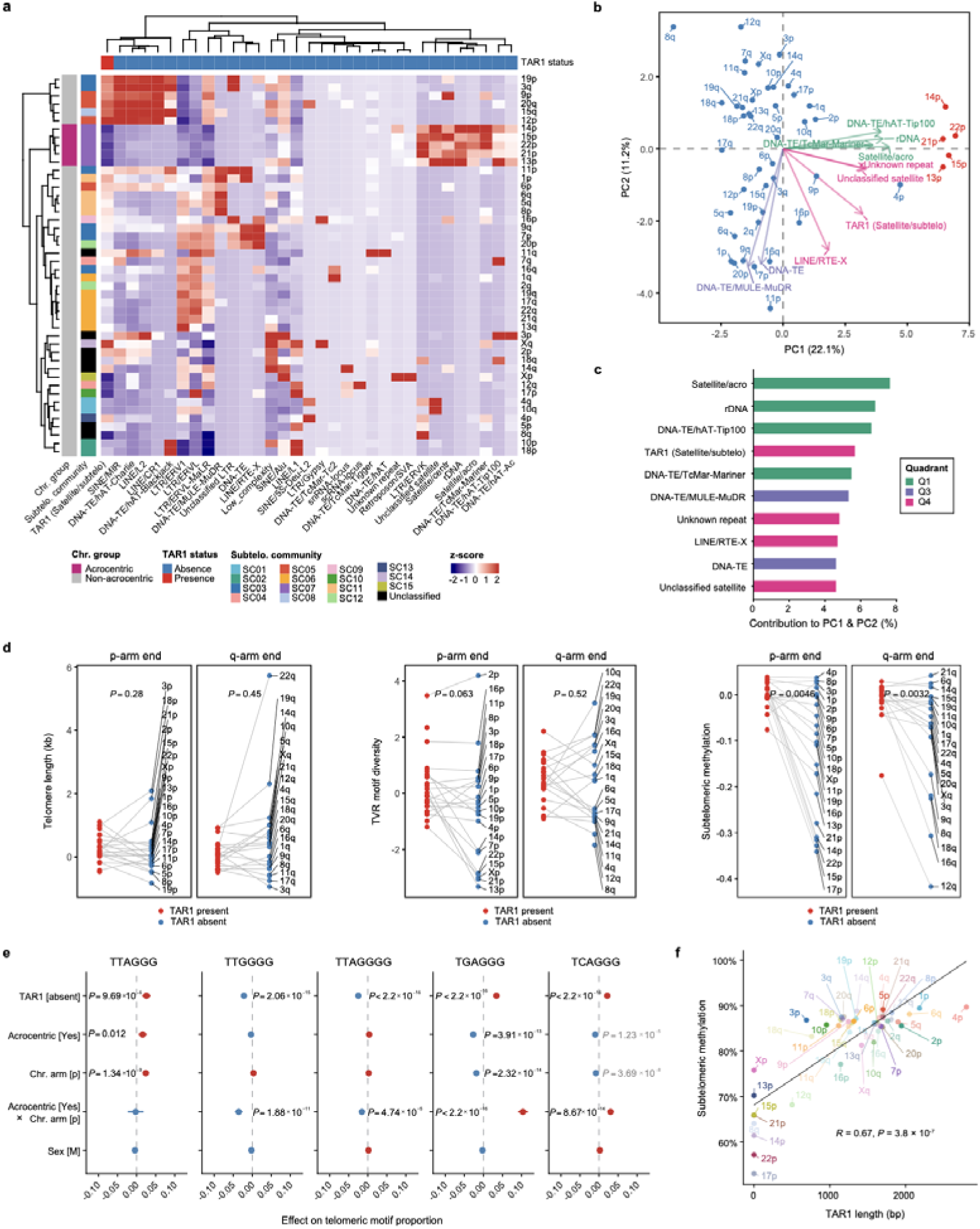
Impact of subtelomeric repeat elements on adjacent telomeres. **(a)** Enrichment of subtelomeric repeat classes at each chromosome end (within 20 kb of telomere–subtelomere boundaries). Chromosome ends are annotated as acrocentric or non-acrocentric and color-coded by subtelomeric community membership. Heatmap values depict *z*-score normalized repeat length for each repeat class. Rows and columns were hierarchical clustered based on Euclidean distance. **(b)** Principal component analysis (PCA) of subtelomeric repeat class composition across chromosome ends. Acrocentric chromosome ends are highlighted in red. Arrows denote the top ten repeat classes contributing to PC1 and PC2, with arrow colors indicating the respective quadrant. **(c)** Percentage contribution of the top ten repeat classes to PC1 and PC2, colored using the same scheme as in (b). **(d)** Effect of TAR1 status (absent vs. present) on telomere length (left), TVR motif diversity (middle), and subtelomeric methylation (right). Grey lines connect measurements from the same chromosome end. *P*-values were calculated using two-sided Wilcoxon rank-sum tests. **(e)** Multivariable linear mixed model effect estimates (with 95% confidence intervals) evaluating the impact of TAR1 status (presence vs. absence within 20-kb subtelomeric region adjacent to the telomere–subtelomere boundary), chromosome type (acrocentric vs. non-acrocentric), chromosome arm (q vs. p), acrocentric chromosome-end (acrocentric and p-arm vs. the others), sex (male vs. female) on the proportions of canonical (TTAGGG) and common non-canonical (TTGGGG, TTAGGGG, TGAGGG, TCAGGG) telomere repeats. Features with |coefficient| ≥ 0.01 were defined as having a high effect size. Significant *P*-values (Adjusted *P* ≤ 0.05) combined with a high effect size are highlighted in black, whereas those with an effect size below this threshold are shown in grey. Non-significant *P*-values are omitted. **(f)** Spearman rank correlation between TAR1 length and subtelomeric methylation level.

TAR1 harbors promoter-like sequences and transcription factor binding sites implicated in TERRA transcription, both of which can regulate telomerase activity. We therefore tested whether local TAR1 status (presence vs. absence) was associated with telomere length, TVR composition, or subtelomeric DNA methylation (Fig. 7d and Supplementary Fig. 31–33). While TAR1 seems not associated with telomere length, we found its presence is associated with significantly greater TVR motif diversity across chr12q and all five acrocentric p-arm ends. At these ends, telomeres lacking adjacent TAR1 contained higher proportions of the canonical TTAGGG repeat as well as common TVRs such as TGAGGG, TCAGGG, and CGAGGG (Fig. 7e and Supplementary Fig. 34). In contrast, other common TVRs including TTGGGG and TTAGGGG were significantly depleted. The broader association between TAR1 presence and increased TVR abundance and diversity is consistent with a contemporaneous HPRC2 analysis^38^.

TAR1 presence also consistently linked to elevated subtelomeric methylation at 16 of the 46 accessible chromosome ends, including chr12q and all five acrocentric p-arm ends. TAR1 length varied markedly among chromosome ends, ranging from truncated 10-bp fragments to extended 3.76-kb arrays. We identified a significant positive correlation between TAR1 length and subtelomeric methylation across chromosome ends (Fig. 7f), echoing our earlier finding in the trio data that TAR1 strongly dictates the local methylation peak at telomere-subtelomere boundaries. Collectively, these results identify TAR1 as a major *cis*-linked feature associated with subtelomeric methylation and telomeric repeat diversity, but not telomere length.

## Discussion

Telomere length, sequence variants, and DNA methylation define complementary dimensions of chromosome-end architecture, reflecting distinct facets of telomere diversity and maintenance. Jointly characterizing these features is therefore essential to understand why individual telomeres differ in function, inheritance and evolutionary history. Historically, however, telomeres have predominantly been represented as bulk, genome-wide length distributions, obscuring critical variation among specific chromosome ends and homologous chromosomes. Leveraging long-read sequencing data, TeloXplorer bridges this resolution gap by integrating chromosome-end-specific and haplotype-resolved profiling of telomere length, sequence composition and DNA methylation into a unified framework. Its robust performance across simulated and empirical datasets from humans, *Arabidopsis* and yeast highlights its power and versatility for high-resolution, multimodal telomere analysis across diverse species with markedly different telomere lengths and repeat architectures. Furthermore, by resolving these features jointly, TeloXplorer reveals multi-layered view demonstrating that telomeres operate as discrete, chromosome-end-specific genomic compartments at both genetic and epigenetic levels, rather than generic, homogeneous terminal arrays.

In addition to relying on long-read sequencing, our analyses demonstrate that the choice of reference genome is integral to achieving accurate, high-resolution telomere analysis. For the human HG002 dataset, replacing the native and haplotype-phased HG002 assembly with the generic CHM13 reference reassigned nearly 35% of telomeric reads to different chromosome ends, with acrocentric chromosomes particularly affected. Thus, reference choice does not merely introduce quantitative noise; it can fundamentally alter the inferred identity of the telomeres being measured. The variable effects observed across additional *Arabidopsis* and yeast samples further suggest that this bias escalates with heterozygosity and subtelomeric structural diversity. Consequently, sample-matched, haplotype-resolved assemblies are expected to provide the most reliable reference system for chromosome-end analysis. Where such assemblies are unavailable, results obtained using generic references should be interpreted cautiously, particularly when evaluating structurally polymorphic chromosome ends.

Our analysis of two ancestrally diverse human trios supports a hierarchical model of telomere inheritance, in which sequence architecture serves as a stable substrate underlying both length control and epigenetic states. At the sequence level, TVR haplotypes exhibited predominant Mendelian transmission across all accessible chromosome ends, preserving parental haplotype-specific signatures. Although sample-specific differences in cell-passage history confounded absolute length comparisons, relative chromosome-end-specific length rankings remained concordant between offspring and parents. Telomere length can be reset during mammalian preimplantation development^40^, and reciprocal-cross experiments in mice show that the direction of early telomere change depends on which parent contributes long or short telomeres^41^. Our haplotype-resolved observations at individual chromosome ends raise the possibility that parental chromosome-end information strongly influences the length set points established during embryonic reprogramming. Such an effect could arise from telomeric or subtelomeric features acting in *cis*. While these two trios are insufficient to establish this mechanism, their length concordance warrants future studies with larger pedigrees and primary tissues. Methylation showed a clearer sequence dependence: telomeric methylation was confined to CpG-containing TVRs, whereas subtelomeric methylation peaked over CpG-rich TAR1 elements. Parent–offspring methylation similarities may therefore reflect sequence-directed reconstruction rather than independent transgenerational inheritance of epigenetic states. Together, these findings suggest that telomere inheritance integrates stable sequence transmission with locally constrained establishment of length and methylation.

At the population level, the conserved chromosome-end-specific telomere length ranking observed across globally diverse human populations recapitulates previous findings from independent cohorts, supporting this pattern as a broadly shared feature of human telomeres. Stable chromosome-end-specific patterns also extended to TVR motif diversity and subtelomeric methylation. Together, these recurrent patterns suggest that local chromosome context substantially influences all three dimensions. Acrocentric chromosome ends are particularly distinctive, combining relatively long telomeres with reduced TVR motif diversity and lower subtelomeric methylation. This combination may reflect a regulatory context that distinguishes acrocentric telomeres from other chromosome ends. Crucially, however, chromosome ends did not follow the same ordering across length, TVR motif diversity and methylation. These dimensions are therefore unlikely to be governed by a single regulatory axis, but instead appear shaped by partially overlapping yet distinct *cis*-acting determinants.

Given their predominant Mendelian transmission across the evaluated trios, TVR haplotypes provide a complementary record of chromosome-end evolution. Across global populations, five chromosome ends (2p, 5p, 12p, 1q, and 7q) each maintained a unique, dominant TVR haplotype, indicative of deep evolutionary conservation. Notably, four of these ends were independently classified as having minimal subtelomeric structural variation in optical mapping data from 154 human genomes^42^. This concordance suggests that a stable subtelomeric background may preserve an ancestral, *cis*-linked proximal TVR architecture. Under this model, distal repeat turnover could alter overall telomere length while preserving proximal haplotype identity. However, this relationship is not universal: 7q retains a dominant TVR architecture despite substantial subtelomeric variation, whereas 18p and 11q exhibit the converse pattern. Subtelomeric stability can therefore explain only part of the observed TVR conservation. A second evolutionary process is implied by extensive interchromosomal TVR haplotype sharing among the *p* arms of all five acrocentric chromosomes, as well as between several non-acrocentric end pairs. These signatures are consistent with recurrent ectopic exchange between sequence-similar subtelomeres via non-allelic homologous recombination or gene conversion^43^. During meiosis, the zygotene bouquet transiently clusters telomeres at the nuclear envelope^44^, providing a plausible spatial context for such interchromosomal encounters^39^. Together, these findings indicate that TVR evolution is shaped by an interplay between local, *cis*-linked stability and episodic interchromosomal exchange. Resolving why the relative contributions of these processes differ among individual chromosome ends remains a key mechanistic challenge.

The elevated TVR haplotype diversity in individuals of African ancestry parallels broader genome-wide patterns, partly reflecting diversity loss in non-African populations during the out-of-Africa bottleneck^45^. However, the present data cannot distinguish demographic history from telomere-specific evolutionary processes. The association between TVR duplication blocks and longer telomeres is similarly non-causal. Duplications could promote elongation, accumulate preferentially within longer tracts, or both. Nevertheless, the coexistence of individual-specific and widely shared duplications suggests that telomere restructuring has occurred over both recent and deeper evolutionary timescales.

Subtelomeric sequences provide a plausible *cis*-acting substrate for this chromosome-end specificity. TAR1 elements are positioned near most human telomere–subtelomere boundaries, where their presence is associated with increased TVR motif diversity at specific ends and elevated subtelomeric methylation. TAR1 length also correlated with methylation level, whereas no corresponding association with telomere length was detected. These results suggest that local sequence composition and local chromatin state co-vary without directly determining telomere tract length. The localization of methylation peaks to TAR1-associated CpG clusters supports a direct structural contribution from the repeat sequence itself, with TAR1-associated chromatin organization and TERRA regulation providing a plausible functional mechanism. Nevertheless, because these associations were evaluated across naturally varying haplotypes, they cannot disentangle TAR1-specific effects from those of co-inherited, linked subtelomeric features. Targeted deletion, insertion, or replacement of TAR1 in otherwise isogenic backgrounds will be required to establish direct causality.

In summary, TeloXplorer provides an integrated framework for multidimensional profiling of telomere length, repeat architecture, and DNA methylation at chromosome-end and haplotype resolution. Its application across species, pedigrees, and global populations reveals telomeres as genetically and epigenetically distinct genomic compartments, defined by a unique interplay among Mendelian transmission of sequence architecture, *cis*-regulatory control of length and methylation, and chromosome-end-specific evolutionary dynamics. Extending TeloXplorer to larger pedigrees, disease cohorts, and controlled perturbation systems will enable direct identification and functional characterization of genetic determinants and *cis*-regulators governing telomere biology. Ultimately, these applications will translate the multidimensional patterns characterized here into causal, mechanistic models of chromosome-end regulation, inheritance, and evolution.

## Methods

### Algorithm design and software implementation

TeloXplorer is a modular computational framework for chromosome-end-resolved telomere analysis using long-read sequencing data and genome assemblies. The framework supports user-defined telomere motifs and includes presets for human, mouse, yeast and *Arabidopsis* telomeres. TeloXplorer was implemented primarily in Python, with C++ routines and utilities integrated to accelerate read screening, telomere-signal smoothing and haplotype-consensus refinement. Under the hood, TeloXplorer comprises six main modules.

1. telox-align. Input long reads were first screened using telogrep, a multithreaded C++ utility that identifies reads containing at least five consecutive copies of a telomeric repeat motif in either orientation. Candidate telomeric reads are subsequently aligned to the sample-matched or generic reference genome assembly using minimap2. Only primary alignments with a read length of ≥1 kb and a mapping quality score of ≥20 were retained for downstream analysis. Reads aligned within predefined chromosome terminal regions are then assigned to the corresponding chromosome ends.
2. telox-length. For each read assigned to a chromosome end, exact motif matches are converted into a base-resolution occupancy profile, from which local motif densities are calculated. The resulting profile is iteratively smoothed using BLOOM, a C++ utility that implements a density-based algorithm for merging adjacent segments with similar signals. Fuzzy motif matching is subsequently used to recover local signal losses attributable to sequencing errors or telomere variant repeats. A directional penalty-and-reward scoring model is then applied from the distal end of the read, and the position yielding the maximum cumulative score is defined as the telomere–subtelomere boundary. Reads are classified as chromosome-end telomeres, internal neotelomeres or minitelomeres according to their genomic mapping positions and terminal motif orientations. The module reports telomere length for each read, together with the mean and median telomere lengths for each chromosome end.
3. telox-tvr. Within the telomeric regions defined by telox-length, sequences are normalized to a common orientation and decomposed into ordered repeat units. Candidate TVRs are first identified by scanning predefined *k*-mer lengths and reading phases for recurrent tandem units, with cyclically equivalent units standardized relative to the canonical telomere motif. Repeat decomposition is then performed using anchor-guided local dynamic programming. Resolved repeat units are encoded as single-character tokens, and reads from the same chromosome end are clustered using hierarchical density*-*based spatial clustering of applications with noise (HDBSCAN) algorithm^46^ based on normalized Levenshtein distances between their token strings. For each cluster, up to 150 of the longest reads are used to construct an initial consensus by adaptive banded partial order alignment (abPOA)^47^, which is subsequently refined through a custom hidden Markov model (HMM). The resulting chromosome-end-level haplotypes are reported as consensus sequences and run-length-encoded repeat architectures.
4. telox-methyl. When a modBAM file is provided, telox-methyl extracts MM/ML-encoded base-modification probabilities from reads overlapping inferred telomere–subtelomere boundaries. The canonical cytosine probability at each eligible site is calculated as one minus the summed probabilities of the declared cytosine modifications. The state with the highest probability among canonical cytosine (C), 5-methylcytosine (5mC) and 5-hydroxymethylcytosine (5hmC) is retained if it passes the corresponding probability threshold. Default thresholds are 0.66 for canonical cytosine and 0.90 for both modified states. Base modification calls are retrieved across telomeres and their adjacent subtelomeric regions. Position coordinates are normalized relative to read strand and chromosome arm, defining the inferred telomere–subtelomere boundary as position zero.
5. telox-asm. When only a genome assembly is available, telox-asm extracts terminal sequences from both ends of each assembled chromosome or contig. Each terminal sequence is analyzed using the BLOOM smoothing and directional boundary-scoring procedures implemented in telox-length. Telomere length is defined as the distance from the assembly terminus to the inferred telomere–subtelomere boundary.
6. telox-reads. In the absence of a suitable reference assembly, telox-reads provides a reference-free mode for analyzing raw long-read data. Telomeric reads are first identified using the screening procedure implemented in telox-align and are subsequently evaluated in both motif orientations using the boundary-detection algorithm implemented in telox-length. The module reports the length and sequence of the telomeric tract in each read, providing a reference-free telomere-length distribution without chromosome-end-specific assignment.

Additional plotting modules including telox-plot-length, telox-plot-reads, telox-plot-tvr-hap, and telox-plot-methyl are provided for visualizing chromosome-end-specific telomere length distributions, read-level TVR architectures, TVR haplotype consensus sequences and methylation profiles respectively. By integrating these core and auxiliary modules, TeloXplorer supports multidimensional telomere analysis across diverse data types and use scenarios.

### Software selection and customization for benchmarking

We systematically evaluated the performance of TeloXplorer alongside four established methods, namely TeloBP^15^ (v1.0.0), Topsicle^48^ (v1.1.0), Telometer^49^ (v1.1), and Telogator2^33^ (v2.2.3), for telomere read identification and telomere length estimation. While TeloXplorer and Topsicle can be directly applied across different species, TeloBP requires custom parameterization in its source code. To enable multi-species benchmarking in TeloBP, we modified its primary execution script (teloBPCmd.py) to expose the compositionGStrand and compositionCStrand parameters, allowing custom non-human telomeric motifs to be specified.

Together, these three methods were included in our benchmarking across all simulated human, *Arabidopsis*, and yeast datasets. In contrast, Telometer employs a regular-expression-based approach strictly optimized for canonical human telomeric motifs, restricting its application to human datasets. Similarly, although Telogator2 is theoretically species-independent, it relies on a pre-curated TVR library for optimal performance. Because comprehensive TVR profiles are currently unavailable for *Arabidopsis* and yeast, we restricted Telogator2 benchmarking to human data.

### Long-read simulation for benchmarking

For the benchmarking analysis, we employed pbsim3^50^ (v3.0.5) to simulate WGS reads for human, *Arabidopsis*, and yeast based on their respective T2T (or near T2T) reference genomes (Supplementary Table 3). For pbsim3, real-data-derived error hidden Markov models (ERRHMMs) and read length distributions were used for simulating ONT (model: ERRHMM-ONT-HQ.model; mean length: 20 kb; standard deviation: 15 kb) and PacBio (model: ERRHMM-RSII.model; mean length: 15 kb; standard deviation: 4 kb) reads respectively. We further modeled combinations of sequencing depths (5×, 10×, 30×, 50×, and 100×) and error rates (1%, 5%, and 10%) within our simulation framework. To account for edge effects in linear genome sampling, we appended a continuous block of Ns (human and *Arabidopsis*: 20 kb; yeast: 10 kb) to all chromosomal termini prior to read simulation. These terminal N-blocks were subsequently trimmed from the simulated reads to ensure they would not interfere with downstream telomere length estimation.

### Benchmarking for telomeric read identification

To rigorously evaluate telomeric read identification, we first employed TeloXplorer’s telox-asm module (v0.5.0) to delineate the telomere–subtelomere boundary for each chromosome end of all three reference genomes used in our benchmarking analysis. This step was executed using the command: telox asm --preset <human/yeast/arabidopsis> --assembly <ref_genome>. Additional manual curation was performed to confirm boundary accuracy.

Simulated reads whose reference-based coordinates overlapped both the telomeric tract (≥ 100 bp for human and *Arabidopsis*; ≥ 30 bp for yeast) and the subtelomeric anchoring region (≥ 200 bp) were designated as bona fide telomeric reads, with their true sampling coordinates serving as the ground truth for chromosome-end assignment. Reads originating from telomeric regions that lacked sufficient subtelomeric anchor overlap were excluded from this analysis. A notable challenge in our benchmarking telomere length estimation across these methods is that several existing methods (e.g., TeloBP, Topsicle, and Telometer) lack an integrated read-mapping module, instead requiring pre-aligned or chromosome-end-extracted reads as input. To ensure fair comparison, we applied TeloXplorer’s telox-align module to map simulated reads back to their corresponding reference genome. This step was executed using the following command: telox align --preset <human/yeast/arabidopsis> --fastq <fastq> --ref --mm2-opts <-ax map-ont/-ax map-pb> --min-mapq 20. Minimap2^51^ (v2.30-r1287) was invoked internally by TeloXplorer for this task. We benchmarked telomeric read identification performance across all tools using precision, recall, and F1 score: true positives (TP) were bona fide telomeric reads correctly identified; false positives (FP) were non-telomeric reads misclassified as telomeric; and false negatives (FN) were bona fide telomeric reads left unidentified. Precision, recall, and F1 score were computed as TP/(TP+FP), TP/(TP+FN), and 2TP/(2TP+FP+FN), respectively.

### Benchmarking for telomere length calculation

Based on the read alignments, we applied identical read filtering criteria during telomere length calculation across all tools: 1) MAPQ ≥ 20; 2) read length ≥ 1000 bp; 3) subtelomeric anchor length ≥ 200 bp; and 4) estimated per-read telomere length ≥ 100 bp for human and *Arabidopsis*, or ≥ 30 bp for yeast. For each chromosome end, we evaluated both the absolute and relative deviation of estimated median telomere lengths from the ground truth, summarizing standard errors of the mean (SEM) for these estimation errors across all chromosome ends.

### HG002 cell culture

The human B-lymphoblastoid cell line HG002 (originally from the Coriell Institute for Medical Research, cat. no. GM24385) was kindly provided by Dr. Fuchou Tang (Peking University). Prior to use, the cell line was authenticated by short tandem repeat (STR) profiling (Shanghai Biowing Applied Biotechnology Co. Ltd, Shanghai, China), to confirm its identity and was routinely confirmed to be free from mycoplasma contamination. Cells were cultured in suspension in RPMI-1640 medium (Gibco, cat. no. C11875500BT) supplemented with 15% fetal bovine serum (FBS, ExCell Bio), and 1% penicillin-streptomycin (Gibco, cat. no. 15140122). The cultures were maintained at 37°C in a humidified incubator with 5% CO_2_ and passaged regularly to maintain exponential growth. Prior to DNA extraction, cell viability was confirmed to be >95% using trypan blue exclusion.

### HMW DNA extraction

High molecular weight genomic DNA of the HG002 cell line was extracted using the QIAGEN Genomic-tip (Cat. no. 13343). DNA purity and concentration were assessed using a NanoDrop One spectrophotometer (Thermo Fisher Scientific) and a Qubit 4 fluorometer (Invitrogen), respectively. DNA integrity and fragment size distribution were confirmed via agarose gel electrophoresis. Following quality control, long DNA fragments were size-selected using the Blue Pippin HT system (Sage Science).

### Nanopore library preparation, sequencing, and basecalling

Sequencing libraries were prepared utilizing the ONT Ligation Sequencing Kit V14 (Cat. no. SQK-LSK114). DNA damage repair and end-preparation were performed using the NEBNext FFPE DNA Repair Mix (New England Biolabs, Cat. no. M6630) and the NEBNext Ultra II End Repair/dA-Tailing Module (New England Biolabs, Cat. no. E7546). The final prepared libraries were loaded onto PromethION R10.4.1 (ONT, Cat. no. FLO-PRO114M) flow cells and sequenced on a Nanopore PromethION P48 platform. Raw POD5 signal data were basecalled with Dorado^52^ (v0.7.2) using the super accuracy (SUP) model (dna_r10.4.1_e8.2_400bps_sup@v5.0.0) together with its compatible all-context 5mC/5hmC modified-base model (dna_r10.4.1_e8.2_400bps_sup@v5.0.0_5mC_5hmC@v1).

### Integrated multidimensional telomere characterization for emprical datasets

We ran TeloXplorer on both our in-house HG002 ONT R10 WGS dataset and various public human, *Arabidopsis*, and yeast sequencing datasets for comprehensive telomere analysis and comparison (Supplementary Table 4). Unless otherwise specified, sample-matched native genome assemblies were used (Supplementary Table 3). Telomere characterization for human, *Arabidopsis*, and yeast was performed using TeloXplorer (v0.5.0) using the following command: telox run --preset <human/arabidopsis/yeast> --fastq <fastq> (or --bam <bam> or --modbam <modbam> depending on the input read format) --ref <ref_genome> --mm2-opts “<-ax map-ont/-ax map-pb/-ax map-hifi>” --min-read-qual 10. For TVR analysis, an additional quality threshold of --min-tel-qual 20 was applied across all datasets, except for PB CLR and ONT R9 data, where more relaxed cutoffs of 0 and 10 were used, respectively, given their lower baseline read qualities. The resulting telomere lengths, sequence compositions, and DNA methylation profiles (if available) were subsequently utilized for downstream analyses.

For the telomere length comparison, we included only chromosome arms with a minimum coverage of five telomeric reads in each compared dataset. Pearson correlation was used for pairwise telomere-length comparison. For TVR comparisons, overall TVR compositions were quantified as normalized motif frequencies (the proportion of each motif relative to total TVRs), and the ten most abundant motifs within each dataset were plotted. Read-level TVR profiles were plotted using TeloXplorer’s plot-reads module.

### Similarity-based TVR haplotype clustering

Pairwise TVR haplotype similarity was calculated using a custom motif-aware Levenshtein algorithm implemented in TeloXplorer. Haplotypes were represented as ordered, length-encoded motif blocks, with comparisons restricted to the first 250 bp distal to the telomere–subtelomere boundary. Dynamic programming was used to align haplotypes with logarithmically scaled penalties for motif copy-number differences, motif substitutions and block insertions or deletions. Default penalty weights were set to 1.0 for copy-number differences, 3.0 for substitution opening, and 0.5 for substitution extension, 2.0 for gap opening, and 0.5 for gap extension, respectively. The resulting alignment cost was normalized to a distance (D) between 0 and 1, and pairwise similarity (S) was defined as (S = 1 − D). TeloXplorer’s plot-tvr-hap module was used for visualization.

In parallel with the HG002 TVR similarity analysis, subtelomeric sequences spanning 1 kb immediately upstream of the telomere–subtelomere boundary were extracted across all chromosome ends. Pairwise sequence identities were calculated using global alignments generated by EMBOSS Needle^53^ (v6.5.7.0; parameters: -gapopen 10 -gapextend 0.5).

### Comparison with estimates from TRF experiments

Previously published telomere estimates from TRF experiments were used to compare with those from TeloXplorer’s sequencing-based estimates. The TRF results of 29 *Arabidopsis* ecotypes and 8 yeast strains were retrieved from previous studies^12,48^. To account for variable subtelomeric structure in yeast chromosome ends, the published TRF values were re-analyzed against strain-specific genome assemblies retrieved from the ScRAPdb database^54^ (Supplementary Fig. 10 and Supplementary Table 5). Specifically, Δ*d* was defined as the distance between the telomere–subtelomere boundary and its nearest upstream XhoI restriction site for each chromsome end, based on which a strain-specific adjusting factor (Δ*d*_mode_) was further calculated as the mode of Δ*d*across all chromosome ends. The adjusted TRF-based telomere estimate was derived as: Adjusted telomere length original TRF length Δ*d*_mode_. These adjusted TRF values were then compared against TeloXplorer estimates using Spearman rank correlation analysis.

### Human trio assembly and read processing

Two widely used human family trios were used for this analysis, which include an Ashkenazi Jewish trio (HG002, son; HG003, father; and HG004, mother), and a Chinese trio (HG005, son; HG006, father; and HG007, mother). Their ONT reads were retrieved from the publicly accessible ONT Open Data repository (release giab_2025.01; Supplementary Tables 3,4). For the HG002/HG003/HG004 trio, the recently published fully phased HG002 assembly was used for the downstream analysis^31^. As for the HG005/HG006/HG007 trio, we retrieved its phased assembly from the HPRC portal (https://data.humanpangenome.org/assemblies) and performed reference-guided scaffolding by RagTag^55^ (v2.1.0) with the following command: ragtag.py scaffold -t <threads> -f 100000 --remove-small -q 60 -a 0.7 -r -u -w -C --aligner minimap2 --mm2-params ’-x asm5 --secondary=no’ -o <output_dir> <ref_genome> <input_assembly>. The human CHM13 reference genome^56^ (v2.0) was used as the reference for this scaffolding analysis. The scaffolded HG005 maternal and paternal haplotype assemblies were used for the downstream analysis. For both HG002 and HG005 assemblies, repeat annotation was conducted following the T2T Consortium protocol. Briefly, we built a custom repeat library by merging the precompiled human library from RepeatMasker^57^ (v4.1.5) with repeat models generated from human Y chromosome and ape autosomal and X/Y chromosome analyses (these repeat models are provided at: https://github.com/jessicaStorer88/RepeatMasker_library_CHM13). The HG002 and HG005 assemblies were subsequently annotated using RepeatMasker (v4.1.5; options: -no_is -xsmall -gff -libdir <custome_repeat_library> -s -species human <input_assembly>).

### HPRC2 genome assembly and read processing

All haplotype-resolved HPRC2 genome assemblies were retrieved from the HPRC portal (https://data.humanpangenome.org/assemblies). For each sample, we performed CHM13-guided scaffolding on the maternal and paternal haplotype assemblies using RagTag^55^ (v2.1.0) following the same protocol used for the HG005 assembly in the trio-based analysis. Similarly, repeat annotation was performed in the same way as described earlier. For each HPRC sample, all BAM-formatted ONT reads (comprising both R9 and R10 chemistries) were downloaded from the HPRC portal (https://data.humanpangenome.org/raw-sequencing-data). Reads originating from the sample were subsequently merged using samtools^58^ (v1.22.1; options: merge -cp).

### Applying TeloXplorer analysis to the HPRC2 cohort

For each HPRC2 sample, we performed full-suite telomere characterization using TeloXplorer (v0.5.0) with the following command: telox run --preset human --modbam <input_modbam> --min-read-qual 10 --min-tel-qual 20 --ref <input_genome_assembly> --mm2-opts “-ax map-ont” --plot-length -W 8 -H 3 --start-step 1 --threads $threads –prefix <output_sample_prefix> --outdir <output_dir>. Additionally, telomere analysis results for chromosome end 18q in the paternal haplotype assembly of sample NA18982 were excluded from downstream analysis due to potential assembly chimerism reported in a recent study^39^.

### Telomere length heterogeneity evaluation

To assess telomere-length heterogeneity in relation to telomerase-positive (TERT+) and alternative lengthening of telomeres (ALT+) cancer-cell profiles, we calculated the coefficient of variation (CV) for each haplotype from the distribution of read-level telomere-length estimates. The CV was defined as the standard deviation divided by the mean telomere length. Based on established thresholds from a previous study^32^, TERT-positive cells exhibit CV < 0.55, reflecting relatively uniform telomere lengths maintained by telomerase activity, while ALT-positive cells display CV > 0.80, consistent with the highly heterogeneous telomere-length distributions characteristic of alternative lengthening mechanisms.

### Identification of telomere fusion events

We detected telomere fusions in long-read sequencing datasets using TelFusDetector ^59^ (v1.0.0). Candidate reads spanning fusion junctions were required to harbor telomeric repeats in both orientations. Specifically, we classified reads as fusion-supporting if they simultaneously contained ≥ 15 TTAGGG and ≥ 15 CCCTAA repeats, including ≥ 5 occurrences each of the tandem dimers (TTAGGG) and (CCCTAA) , as previously defined ^59^. The telomere fusion rate per sample was calculated as the number of fusion-supporting reads divided by the total count of telomere-containing reads.

### Chromosome-end-specific TVR motif diversity quantification

TVR content was quantified over the first 1 kb of telomeric sequence from the subtelomere–telomere boundary towards the chromosome terminus. Motif blocks extending beyond this window were truncated at the 1-kb boundary. The TVR diversity score for a chromosome end was defined as

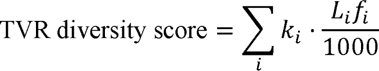

Where *i* indexes the consensus sequences recovered at a given chromosome end,*k_i_* is the number of distinct TVR motifs in consensus *i, L_i_* is the total clipped length (bp) of its TVR motif blocks, and *f_i_* is the number of reads supporting consensus *i* divided by the total number of reads mapped to that chromosome end. The canonical TTAGGG repeat contributes zero to both *K_i_*and *L_i_*. The score was calculated separately for each consensus before summation across consensuses.

### Chromosome-end-specific subtelomeric methylation level quantification

TeloXplorer reports valid CpG positions and per-read methylation calls relative to the subtelomere–telomere boundary. Valid CpG sites within ±5 kb of the boundary were considered. Sites were classified as telomeric or subtelomeric according to their signed position and chromosome arm. For each chromosome end of each haplotype and each region, methylation level was calculated as the number of methylated sites divided by the number of valid CpG sites, yielding the CpG methylation level within the corresponding region.

### Telomere-associated metric normalization

To evaluate relative differences across chromosome ends, we performed median-centered normalization for the three telomere-associated metrics (i.e., telomere length, TVR motif diversity, and subtelomeric methylation level) calculated for each haplotype-resolved HPRC2 genome assembly. For each of these metrics, the median value across all chromosome ends within the maternal or paternal haplotype assembly was subtracted from each individual chromosome-end estimate. This normalization reduces between-haplotype baseline shifts while preserving within-haplotype variation among chromosome ends.

### TVR duplication block detection

Within each telomere consensus sequence, TVR blocks were tokenized based on motif identity alone, explicitly omitting motif copy number. The short arms of the five acrocentric chromosomes (13p, 14p, 15p, 21p, and 22p), as well as Xp, 8q, and 12q, were excluded in this analysis due to pervasive TVR duplication that confound accurate quantification. A sliding window of five consecutive tokens with a step size of one token was applied to each remaining sequence. Candidate window pairs were required to display identical five-token sequences, starting indices separated by at least five tokens, and a minimum of three consecutively shifted matching windows. Consecutive matching windows were merged into paired blocks; pairs whose genomic lengths differed by more than 20% relative to the longer copy were filtered out. Finally, manually curated false-positive duplication calls were removed.

### HPRC2 TVR haplotype filtering

Before cross-cohort analysis, TVR haplotypes identified by TeloXplorer across individual HPRC2 haplotype assemblies were filtered to restrict representation to a maximum of two top-frequency haplotypes per chromosome end. For chromosome ends resolved to a single TVR haplotype, the haplotype was retained only if it achieved an allele frequency of at least 0.4. For chromosome ends harboring multiple distinct TVR haplotypes, a minimum allele frequency threshold of 0.2 was applied to each individual haplotype.

### HPRC2 TVR haplotype clustering

To define recurrent TVR haplotype structures across the HPRC2 cohort, we performed unsupervised density-based clustering using the Hierarchical Density*-*Based Spatial Clustering of Applications with Noise (HDBSCAN) algorithm^46^. For each chromosome end, pairwise TVR haplotype similarity scores were calculated from the filtered haplotypes using the plot-tvr-hap module of TeloXplorer, producing an end-specific haplotype similarity matrix. Similarity matrices were converted to distance matrices by taking the complement of each similarity score (distance = 1 − similarity). HDBSCAN was then applied directly to the precomputed distance matrices using a minimum cluster size of 10, a minimum samples value of 3, and a cluster selection epsilon of 0.2. Haplotypes that were not assigned to a discrete structural cluster by HDBSCAN were designated as “Unclassified”.

### Inter-chromosomal TVR haplotype sharing index

To quantify the extent to which different chromosomes shared structurally similar TVR haplotypes, we constructed a weighted *k*-nearest-neighbor graph from the global pairwise similarity matrix. Each haplotype was connected to its 30 most structurally similar haplotypes, and reciprocal connections were combined to generate an undirected graph. For each chromosome end pair, the observed total weight of inter-end connections (W_obs_) was compared with that expected from the weighted degrees (W_exp_) of the two arms under a random distribution model. The resulting enrichment (E = W_obs_/W_exp_) was transformed into a bounded inter-chromosomal haplotype sharing index (HSI) using the following equation: HSI = 1 − exp(−0.5 × max(0, E − 1)). For network visualization, each chromosome-end node was depicted as a pie chart illustrating its composition of HDBSCAN-defined haplotype clusters, with edge width scaled to represent the inter-chromosomal haplotype-sharing index.

### Population-level diversity analysis of TVR haplotypes

TVR haplotype diversity across five HPRC continental groups (AFR, AMR, SAS, EUR, and EAS) was evaluated by computing the Shannon entropy index using the R package vegan^60^ (v2.7-2). Within each continental group, diversity was estimated independently for each chromosome end, restricting analysis to ends represented by at least 15 haplotypes in the corresponding continental group. Classified haplotypes were categorized according to their HDBSCAN cluster assignments, whereas each unclassified haplotype was treated as a unique variant to preserve the contribution of rare, private structures to overall diversity.

To control for sampling biases arising from unequal haplotype counts across populations, we performed rarefaction by randomly sampling 15 haplotypes without replacement per eligible chromosome arm in each population over 1,000 iterations. The Shannon entropy was calculated for each iteration, and the mean index across iterations was defined as the final diversity estimate. Differences across continental groups were evaluated per chromosome end using paired Wilcoxon signed-rank tests, with AFR serving as the baseline reference.

### Subtelomeric repeat composition and clustering

We analyzed subtelomeric repeat compositions using annotations retained within the 20-kb region immediately upstream of the telomere boundary. To determine repeat composition, we calculated the class-specific repeat proportions per haplotype by dividing each class’s cumulative length by the total repeat length within the 20-kb window. We then averaged these proportions across all haplotypes for each chromosome arm. The resulting matrix was -score standardized by repeat class. Chromosome ends were designated as acrocentric or non-acrocentric and assigned to subtelomeric communities as defined by a recent study^39^. Hierarchical clustering of rows and columns was performed using Euclidean distance and Ward’s D2 linkage with ComplexHeatmap^61^ (v2.22.0). For PCA, each chromosome end was represented by the mean interval length per repeat class within the 20-kb window. After excluding zero-variance classes and centering/scaling the data, PCA was executed using the prcomp function in R^62^ (v4.4.2). The top 10 repeat classes with the highest contribution scores to PC1 and PC2 were identified.

### Multivariable linear mixed-effects modeling of TVR composition

For each telomere repeat motif, a separate multivariable linear mixed model was fitted to estimate the effects of TAR1 status (TAR1-present vs. TAR1-absent within the telomere-proximal 20 kb), chromosome group (acrocentric vs. non-acrocentric), chromosome arm (p vs. q), the interaction between chromosome group and arm, and sex (male vs. female) on the proportion contributed by that motif at each chromosome end. All terms were entered simultaneously, with a random intercept per individual to account for repeated measurements across chromosome ends and haplotypes. The reference levels were TAR1-present, non-acrocentric, q arm and female. Models were fitted by restricted maximum likelihood (REML) using the lmer function from the R package lmerTest^63^ (v3.2-1). Fixed-effect estimates and 95% confidence intervals were reported on the motif-proportion scale. Raw coefficient *P* values were pooled across all motif-specific models and all non-intercept fixed-effect coefficients, then adjusted using the Benjamini–Hochberg procedure to control the false discovery rate; adjusted *P* values ≤ 0.05 were considered significant. An absolute fixed-effect coefficient ≥ 0.01 on the motif proportion was considered as a high effect.

### Generative AI use

Generative AI tools were used exclusively to refine grammar, clarity, and readability. All AI-generated suggestions were critically reviewed and edited by the authors, who take full responsibility for the final manuscript.

## Data and code availability

The sequencing data of the HG002 cell line generated by this study is deposited at Sequence Read Archive (SRA) under accession numbers of PRJNA1511489. Information about all previously published sequencing data analyzed in this study can be found in Supplementary Tables 3, 4, and 7. TeloXplorer is free for use under the GNU GPL v3.0 License, with the source code available on GitHub ( https://github.com/hhuili/TeloXplorer ).

## Funding

This work is supported by National Natural Science Foundation of China (32470663 to J.-X.Y.), Guangdong Provincial Pearl River Talents Program (2019QN01Y183 to J.-X.Y.), China Postdoctoral Science Foundation (2023M744080 to C.C.), and Young Talents Program of Sun Yat-sen University Cancer Center (YTP-SYSUCC-0042 to J.-X.Y.). The funders have not played any role in the study design, data collection and analysis, decision to publish, or preparation of the manuscript.

## Author contributions

J.-X.Y. designed and supervised this study. H.L., Z.M., L.Y. developed the software. H.L., C.C., L.Y. and Z.M. analyzed the data and visualized the results. J.-X.Y., C.C., L.Y., Y.S. and W.B. tested the software. J.-X.Y. drafted the initial manuscript and all authors contributed to the manuscript. All authors read and approved the manuscript.

## Competing interests

The authors declare no competing interests.

## Acknowledgements

We thank the valuable discussion on yeast telomere TRF analysis from Dr. Melania Jennifer D’Angiolo (University College London) and Dr. Xue-Ting Zhu (CAS Center for Excellence in Molecular Cell Science). We thank Dr. Gianni Liti (Institute for Research on Cancer and Aging, Nice), Dr. Jin-Qiu Zhou (CAS Center for Excellence in Molecular Cell Science), and Dr. Aaron Mendez-Bermudez (Institute for Research on Cancer and Aging, Nice) for critically reading this manuscript and providing insightful feedback. We are grateful to Dr. Fuchou Tang (Peking University) for sharing the HG002 cell line for our sequencing experiment. We thank the facility support from the Single-Molecule Sequencing Platform and the Bioinformatics Platform at Sun Yat-sen University Cancer Center. We would also like to acknowledge the National Human Genome Research Institute (NHGRI) for funding the following grants supporting the creation of the human pangenome reference: U41HG010972, U01HG010971, U01HG013760, U01HG013755, U01HG013748, U01HG013744, R01HG011274, and the Human Pangenome Reference Consortium (BioProject ID: PRJNA730823).

## Human Pangenome Reference Consortium

Derek Albracht^2^, Ivan A. Alexandrov^3^, Jamie Allen^4^, Alawi A. Alsheikh-Ali^5^, Nicolas Altemose^6^, Casey Andrews^7^, Dmitry Antipov^8^, Lucinda Antonacci-Fulton^2^, Alexander Arguello^9^, Mobin Asri^10^, Marcelo Ayllon^11^, Jennifer R. Balacco^12^, Floris P. Barthel^13^, Edward A. Belter Jr^2^, Halle D. Bender^10^, Andrew P. Blair^10^, Davide Bolognini^14^, Katherine E. Bonini^15^, Christina Boucher^16^, Guillaume Bourque^17,18,19^, Silvia Buonaiuto^20^, Shuo Cao^20^, Andrew Carroll^21^, Ann M. Mc Cartney^22^, Monika Cechova^10^, Mark J.P. Chaisson^23^, Pi-Chuan Chang^21^, Xian Chang^10^, Jitender Cheema^4^, Haoyu Cheng^24^, Claudio Ciofi^25^, Hiram Clawson^10^, Sarah Cody^2^, Vincenza Colonna^20^, Holland C. Conwell^26^, Robert Cook-Deegan^27^, Mark Diekhans^10^, Maria Angela Diroma^25^, Daniel Doerr^28,29,30^, Zheng Dong^7^, Danilo Dubocanin^6^, Richard Durbin^31,32^, Jana Ebler^28,33^, Evan E. Eichler^11,34^, Jordan M. Eizenga^10^, Parsa Eskandar^10^, Eddie Ferro^16^, Anna-Sophie Fiston-Lavier^35,36^, Sarah M. Ford^26^, Willard W. Ford^37^, Giulio Formenti^12^, Adam Frankish^4^, Mallory A. Freeberg^4^, Qichen Fu^7^, Stephanie M. Fullerton^38^, Robert S. Fulton^2^, Shenghan Gao^39^, Yan Gao^40^, Gage H. Garcia^11^, Obed A. Garcia^41^, Joshua M.V. Gardner^10^, Shilpa Garg^42^, Erik Garrison^20^, Nanibaa’ A. Garrison^43,44,45^, John E. Garza^2^, Margarita Geleta^46,47^, Mohammadmersad Ghorbani^48^, Tina A. Graves-Lindsay^2^, Richard E. Green^26^, Carol W. Greider^49^, Cristian Groza^50^, Bida Gu^23^, Andrea Guarracino^13,20^, Melissa Gymrek^51^, Maximilian Haeussler^10^, Leanne Haggerty^4^, Ira M. Hall^52,53^, Nancy F. Hansen^8^, Yue Hao^13^, Mohammad Amiruddin Hashmi^5^, David Haussler^10^, Prajna Hebbar^10^, Peter Heringer^28,29,30^, Glenn Hickey^10^, Todd L. Hillaker^10^, S. Nakib Hossain^4^, Neng Huang^40,54^, Sarah E. Hunt^4^, Toby Hunt^4^, Alexander G. Ioannidis^6,10,47^, Nafiseh Jafarzadeh^10^, Nivesh Jain^12^, Erich D. Jarvis^12,34^, Maryam Jehangir^13^, Juan Jiang^7^, Eimear E. Kenny^15^, Juhyun Kim^8^, Bonhwang Koo^12^, Sergey Koren^8^, Milinn Kremitzki^2,7^, Charles H. Langley^55^, Ben Langmead^56^, Heather A. Lawson^7^, Daofeng Li^7^, Heng Li^40,54^, Ronghan Li^7^, Wen-Wei Liao^52,53^, Jiadong Lin^11^, Tianjie Liu^7^, Glennis A. Logsdon^39^, Ryan Lorig-Roach^10^, Jonathan LoTempio Jr^22,57^, Hailey Loucks^10^, Jane E. Loveland^4^, Jianguo Lu^58^, Shuangjia Lu^52,53^, Julian K. Lucas^10^, Walfred Ma^23^, Juan F. Macias-Velasco^2,7,59^, Kateryna D. Makova^60^, Maximillian G. Marin^40,54^, Christopher Markovic^2^, Tobias Marschall^28,33^, Franco L. Marsico^20^, Fergal J. Martin^4^, Mira Mastoras^10^, Capucine Mayoud^35^, Brandy McNulty^10^, Jack A. Medico^12^, Julian M. Menendez^10^, Karen H. Miga^10^, Anna Minkina^61^, Matthew W. Mitchell^62^, Saswat K. Mohanty^63^, Younes Mokrab^48,64,65^, Jean Monlong^66^, Shabir Moosa^48^, Avelina Moreno-Ochando^67,68^, Shinichi Morishita^69^, Jonathan M. Mudge^4^, Katherine M. Munson^11^, Njagi Mwaniki^70^, Nasna Nassir^5^, Chiara Natali^25^, Shloka Negi^10^, Lingbin Ni^11^, Adam M. Novak^10^, Faith Okamoto^10^, Keisuke K. Oshima^39^, Pilar N. Ossorio^71,72^, Chie Owa^69^, Sadye Paez^12^, Benedict Paten^10^, Clelia Peano^14,73^, Adam M. Phillippy^8,56,74,75^, Brandon D. Pickett^8^, Laura Pignata^20^, Nadia Pisanti^70^, David Porubsky^11,76^, Pjotr Prins^20^, Timofey Prodanov^28,33^, Anandi Radhakrishnan^10^, T. Rhyker Ranallo-Benavidez^13^, Brian J. Raney^10^, Mikko Rautiainen^77^, Alessandro Raveane^14^, Andreas Rechtsteiner^49^, Luyao Ren^11,34^, Arang Rhie^8^, Fedor Ryabov^78,79^, Samuel Sacco^26^, Farnaz Salehi^20^, Michael C. Schatz^56,80^, Laura B. Scheinfeldt^81^, Aarushi Sehgal^37^, William E. Seligmann^26^, Mahsa Shabani^82^, Kishwar Shafin^21^, Shadi Shahatit^35^, Ruhollah Shemirani^15^, Vikram S. Shivakumar^56^, Swati Sinha^4^, Jouni Sirén^10^, Linnéa Smeds^63^, Steven J. Solar^8^, Marco Sollitto^12,25^, Nicole Soranzo^14,31,83^, Andrew B. Stergachis^11,61^, Marie-Marthe Suner^4^, Yoshihiko Suzuki^69^, Arda Söylev^28,33^, Ahmad Abou Tayoun^84,85^, Jack A.S. Tierney^4^, Chad Tomlinson^2^, Francesca Floriana Tricomi^4^, Mohammed Uddin^5,86^, Matteo Tommaso Ungaro^26,87^, Rahul Varki^16^, Flavia Villani^20^, Ivo Violich^10^, Mitchell R. Vollger^88^, Brian P. Walenz^8^, Charles Wang^89^, Lisa E. Wang^15^, Ting Wang^2,7,59^, Aaron M. Wenger^90^, Conor V. Whelan^12^, Zilan Xin^7^, Zheng Xu^7^, Kai Ye^91^, DongAhn Yoo^11^, Wenjin Zhang^7^, Ying Zhou^40^, Xiaoyu Zhuo^7^, Giulia Zunino^14^

^2^ McDonnell Genome Institute, Washington University School of Medicine, St. Louis, MO 63108, USA

^3^ Department of Human Molecular Genetics and Biochemistry, Faculty of Medical and Health Sciences, Tel Aviv University, Tel Aviv 69978, Israel

^4^ European Molecular Biology Laboratory, European Bioinformatics Institute (EMBL-EBI), Wellcome Genome Campus, Hinxton, Cambridge CB10 1SD, UK

^5^ Center for Applied and Translational Genomics (CATG), Mohammed Bin Rashid University of Medicine and Health Sciences, Dubai Health, Dubai, UAE

^6^ Department of Genetics, Stanford University, Palo Alto, CA 94304 USA

^7^ Department of Genetics, Washington University School of Medicine, St. Louis, MO 63110, USA

^8^ Genome Informatics Section, Center for Genomics and Data Science Research, National Human Genome Research Institute, National Institutes of Health, Bethesda, MD 20892, USA

^9^ Division of Genome Sciences, National Human Genome Research Institute, Bethesda, MD 20871 USA

^10^ UC Santa Cruz Genomics Institute, University of California, Santa Cruz, CA 95060, USA

^11^ Department of Genome Sciences, University of Washington School of Medicine, Seattle, WA 98195, USA

^12^ The Vertebrate Genome Laboratory, The Rockefeller University, New York, NY 10065, USA

^13^ Bioinnovation and Genome Sciences, The Translational Genomics Research Institute (TGen), Phoenix, AZ 85004, USA

^14^ Human Technopole, Milan, Italy

^15^ Institute for Genomic Health, Icahn School of Medicine at Mount Sinai, New York, NY 10029, USA

^16^ Department of Computer and Information Science and Engineering, University of Florida, Gainesville, FL 32611, USA

^17^ Canadian Center for Computational Genomics, McGill University, Montréal, QC H3A 0G1, Canada

^18^ Department of Human Genetics, McGill University, Montréal, QC H3A 0G1, Canada

^19^ Victor Phillip Dahdaleh Institute of Genomic Medicine, Montréal, QC H3A 0G1, Canada

^20^ Department of Genetics, Genomics and Informatics, University of Tennessee Health Science Center, Memphis, TN 38163, USA

^21^ Google LLC, Mountain View, CA 94043, USA

^22^ Institute of Clinical and Translational Sciences, University of California, Irvine, CA 92697, USA

^23^ Quantitative and Computational Biology, University of Southern California, Los Angeles, CA 90089, USA

^24^ Department of Biomedical Informatics and Data Science, Yale School of Medicine, New Haven, CT 06510, USA

^25^ Department of Biology, University of Florence, Sesto Fiorentino, FI 50019, Italy

^26^ Department of Ecology and Evolutionary Biology, University of California, Santa Cruz, CA 95060, USA

^27^ Arizona State University, Consortium for Science, Policy & Outcomes, Washington, DC 20006, USA

^28^ Center for Digital Medicine, Heinrich Heine University Düsseldorf, Düsseldorf, NRW, DE

^29^ Department for Endocrinology and Diabetology at the Medical Faculty and University Hospital Düsseldorf, Heinrich Heine University Düsseldorf, Düsseldorf, NRW, DE

^30^ Paul-Langerhans-Group Computational Diabetology, German Diabetes Center (DDZ) and Leibniz Institute for Diabetes Research, Düsseldorf, NRW, DE

^31^ Wellcome Sanger Institute, Genome Campus, Hinxton, CB10 1RQ, UK

^32^ Department of Genetics, University of Cambridge, Cambridge, CB2 3EH, UK

^33^ Institute for Medical Biometry and Bioinformatics, Medical Faculty and University Hospital Düsseldorf, Heinrich Heine University, Düsseldorf, NRW, DE

^34^ Howard Hughes Medical Institute, Chevy Chase, MD 20815, USA

^35^ ISEM, Univ Montpellier, CNRS, IRD, Montpellier, FR

^36^ Institut Universitaire de France, Paris, FR

^37^ Department of Computer Science and Engineering, University of California San Diego, La Jolla, CA 92093, USA

^38^ Department of Bioethics & Humanities, University of Washington School of Medicine, Seattle, WA 98195, USA

^39^ Department of Genetics, Epigenetics Institute, Perelman School of Medicine, University of Pennsylvania, Philadelphia, PA 19104, USA

^40^ Department of Data Science, Dana-Farber Cancer Institute, Boston, MA 02215, USA

^41^ Department of Anthropology, University of Kansas, Lawrence, KS 66045, USA

^42^ School of Health Sciences, University of Manchester, Manchester M13 9PL, UK

^43^ Traditional, ancestral and unceded territory of the Gabrielino/Tongva peoples, Institute for Society & Genetics, University of California, Los Angeles, Los Angeles, CA 90095, USA

^44^ Traditional, ancestral and unceded territory of the Gabrielino/Tongva peoples, Institute for Precision Health, David Geffen School of Medicine, University of California, Los Angeles, Los Angeles, CA 90095, USA

^45^ Traditional, ancestral and unceded territory of the Gabrielino/Tongva peoples, Division of General Internal Medicine & Health Services Research, David Geffen School of Medicine, University of California, Los Angeles, Los Angeles, CA 90095, USA

^46^ Department of Electrical Engineering and Computer Science, University of California, Berkeley, Berkeley, CA 94720, USA

^47^ Department of Biomedical Data Science, Stanford University School of Medicine, Stanford, CA 94305, USA

^48^ Medical and Population Genomics Lab, Sidra Medicine, Doha, Qatar

^49^ Department of Molecular Cell and Developmental Biology, University of California, Santa Cruz, CA, USA

^50^ Montreal Heart Institute, Montréal, QC, Canada

^51^ Department of Pediatrics, University of California San Diego, La Jolla, CA 92093, USA

^52^ Center for Genomic Health, Yale University School of Medicine, New Haven, CT 06510, USA

^53^ Department of Genetics, Yale University School of Medicine, New Haven, CT 06510, USA

^54^ Department of Biomedical Informatics, Harvard Medical School, Boston, MA 02115, USA

^55^ Department of Evolution and Ecology and the Center for Population Biology, University of California, One Shields, Davis, CA 95616, USA

^56^ Department of Computer Science, Johns Hopkins University, Baltimore, MD 21218, USA

^57^ Department of Pediatrics, Division of Genetics, School of Medicine, University of California, Irvine, CA 92697, USA

^58^ Sun Yat-sen University, Guangzhou, China

^59^ Edison Family Center for Genome Sciences & Systems Biology, Washington University School of Medicine, St. Louis, MO 63110, USA

^60^ Department of Biology and Center for Medical Genomics, Penn State University, University Park, PA 16802, USA

^61^ Division of Medical Genetics, Department of Medicine, University of Washington School of Medicine, Seattle, WA 98195, USA

^62^ The Jackson Laboratory for Genomic Medicine, Farmington, CT 06032, USA

^63^ Department of Biology, Penn State University, University Park, PA 16802, USA

^64^ Department of Biomedical Science, College of Health Sciences, Qatar University, Doha, Qatar

^65^ Department of Genetic Medicine, Weill Cornell Medicine-Qatar, Doha, Qatar

^66^ IRSD - Digestive Health Research Institute, University of Toulouse, INSERM, INRAE, ENVT, UPS, Toulouse, FR

^67^ MATCH biosystems, S.L., Elche, Spain

^68^ Universidad Miguel Hernández de Elche, Elche, Spain

^69^ Department of Computational Biology and Medical Sciences, The University of Tokyo, Kashiwa, Chiba 277-8561, Japan

^70^ Department of Computer Science, University of Pisa, Pisa, Italy

^71^ Law School, University of Wisconsin-Madison, Madison, WI 53706, USA

^72^ Morgridge Institute for Research, Madison, WI 53715, USA

^73^ Institute of Genetics and Biomedical Research, UoS of Milan, National Research Council, Milan, Italy

^74^ Department of Biomedical Engineering, Johns Hopkins University, Baltimore, MD 21218, USA

^75^ Department of Genetic Medicine, Johns Hopkins University School of Medicine, Baltimore, MD 21205, USA

^76^ Genome Biology Unit, European Molecular Biology Laboratory (EMBL), Heidelberg, DE

^77^ Institute for Molecular Medicine Finland, Helsinki Institute of Life Science, University of Helsinki, Helsinki, Finland

^78^ The Center for Bio- and Medical Technologies, Moscow, RUS

^79^ Centre for Biomedical Research and Technology, HSE University, Moscow, RUS

^80^ Department of Biology, Johns Hopkins University, Baltimore, MD 21218, USA

^81^ Coriell Institute for Medical Research, Camden, NJ 08103, USA

^82^ University of Amsterdam, Amsterdam, Netherlands

^83^ School of Clinical Medicine, University of Cambridge, Cambridge, CB2 0SP, UK

^84^ Center for Genomic Discovery, Mohammed Bin Rashid University, Dubai Health, UAE

^85^ Dubai Health Genomic Medicine Center, Dubai Health, UAE

^86^ GenomeArc Inc, Mississauga, ON, Canada

^87^ Department of Biology and Biotechnologies “Charles Darwin”, University of Rome “La Sapienza”, Rome 00185, IT

^88^ Department of Human Genetics and Utah Center for Genetic Discovery, University of Utah, Salt Lake City, UT, USA

^89^ Center for Genomics, Loma Linda University School of Medicine, Loma Linda, CA 92350, USA

^90^ PacBio, Menlo Park, CA 94025, USA

^91^ The first affiliated hospital of Xi’an Jiaotong University, Xi’an Jiaotong University, Xi’an, Shaanxi, 710049, China

## Notes

### Competing Interest Statement

The authors have declared no competing interest.

### Summary of Updates

The first and last name of co-author Yan Shuai (First: Yan, Last: Shuai) were accidentally inverted in the original version, which has been corrected now.

