## Supplementary Fig. for "Multidimensional telomere diversity and inheritance at individual and population scales"

### Supplementary Figures


**Figure S1. Assembly-based identification of telomere–subtelomere boundaries and chromosome-end-specific telomere length estimates.** 3

**Figure S2. Precision of identifying chromosome-end-specific telomere containing reads using simulated long-read data.** 4

**Figure S3. Recall of identifying chromosome-end-specific telomere containing reads using simulated long-read data.** 5

**Figure S4. F1 score of identifying chromosome-end-specific telomere containing reads using simulated long-read data.** 6

**Figure S5. Benchmarking telomere length estimation by absolute and relative telomere length difference using simulated long-read data.** 7

**Figure S6. Comparison of TeloXplorer and Telogator2 for read-level and haplotype-level TVR analysis in human telomeres.** 8

**Figure S7. Correlation of chromosome-end-specific telomere lengths across human HG002 sequencing datasets.** 9

**Figure S8. Cross-platform read-level telomere variant repeat (TVR) profiling of human telomeres.** 10

**Figure S9. Genome-wide telomere length comparison between TeloXplorer and terminal restriction fragment (TRF) profiling.** 11

**Figure S10. Adjusting yeast telomere length estimates from Telomere Restriction Fragment (TRF) data by accounting for subtelomeric variation.** 12

**Figure S11. Cross-platform read-level telomere variant repeat (TVR) profiling of *Arabidopsis* telomeres.** 13

**Figure S12. Cross-platform read-level telomere variant repeat (TVR) profiling of yeast telomeres.** 14

**Figure S13. Comparison of chromosome-end-specific telomere length estimates using native versus generic reference genomes.** 15

**Figure S14. Representative examples of TVR mutations in the child (HG002) in comparison to his parents (father: HG003, mother: HG004).** 16

**Figure S15. Allele-specific inheritance of telomere variant repeat (TVR) composition across the HG005/HG006/HG007 trio.** 17

**Figure S16. Allele-specific inheritance of telomeric and subtelomeric DNA methylation across the HG005/HG006/HG007 trio.** 18

**Figure S17. Telomere length distribution of the HPRC2 cohort.** 19

**Figure S18. Screening for Alternative Lengthening of Telomeres (ALT) candidates across the HPRC2 cohort.** 20

**Figure S19. Cross-cohort consistency of chromosome-end-specific telomere length rankings.** 21

**Figure S20. Chromosome-end-specific telomeric motif composition across the HPRC2 cohort.** 22

**Figure S21. Interrelationships among chromosome-end-specific telomere**

length, telomere variant repeat (TVR) motif diversity, and subtelomeric methylation in the HPRC2 cohort. 23
**Figure S22. Comparison of Telomeric and subtelomeric features between acrocentric and non-acrocentric chromosome ends in the HPRC2 cohort.** 24
**Figure S23. Cumulative global frequencies of chromosome-end-specific telomere variant repeat (TVR) haplotype cluster in the HPRC2 cohort.** 25
**Figure S24. Haplotype-resolved telomere variant repeat (TVR) profile for the HPRC2 cohort.** 26
**Figure S25. Duplication blocks of telomere variant repeat (TVR) identified in the p-arm telomeres across the HPRC2 cohort.** 27
**Figure S26. Duplication blocks of telomere variant repeat (TVR) identified in the q-arm telomeres across the HPRC2 cohort.** 28
**Figure S27. Structural characteristics of telomere variant repeat (TVR) duplication blocks.** 29
**Figure S28. Spatial distribution of subtelomeric repeat elements relative to the telomere–subtelomere boundary across the HPRC2 cohort.** 30
**Figure S29. Prevalence of TAR1 subtelomeric elements across the HPRC2 cohort.** 31
**Figure S30. Physical distribution of TAR1 elements across individual chromosome ends in the HPRC2 cohort.** 32
**Figure S31. Impact of subtelomeric TAR1 presence on chromosome-end-specific telomere length in the HPRC2 cohort.** 33
**Figure S32. Impact of subtelomeric TAR1 presence on chromosome-end-specific telomere variant repeat (TVR) motif diversity in the HPRC2 cohort.** 34
**Figure S33. Impact of subtelomeric TAR1 presence on chromosome-end-specific subtelomeric methylation levels in the HPRC2 cohort.** 35
**Figure S34. Multivariable associations of subtelomeric and demographic features with telomere variant repeat composition in the HPRC2 cohort.** 36

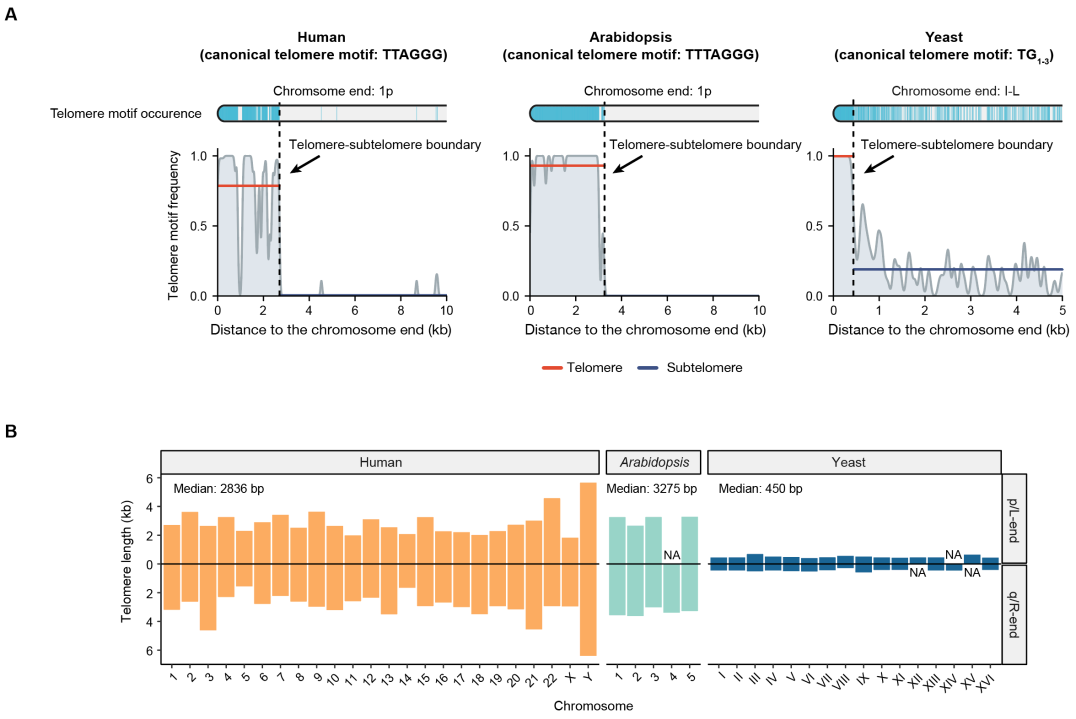


**Figure S1. Assembly-based identification of telomere–subtelomere boundaries and chromosome-end-specific telomere length estimates.**

(A) Telomere–subtelomere boundaries identified by TeloXplorer across human, *Arabidopsis*, and yeast genome assemblies. Occurrences of species-specific canonical telomere motifs are highlighted for representative chromosome ends: human chromosome 1p, *Arabidopsis* chromosome 1p, and yeast chromosome I left end (I-L). Corresponding motif frequencies calculated using a sliding-window approach are shown below. (B) Chromosome-end-specific telomere lengths quantified by TeloXplorer using the same genome assemblies for human, *Arabidopsis*, and yeast.

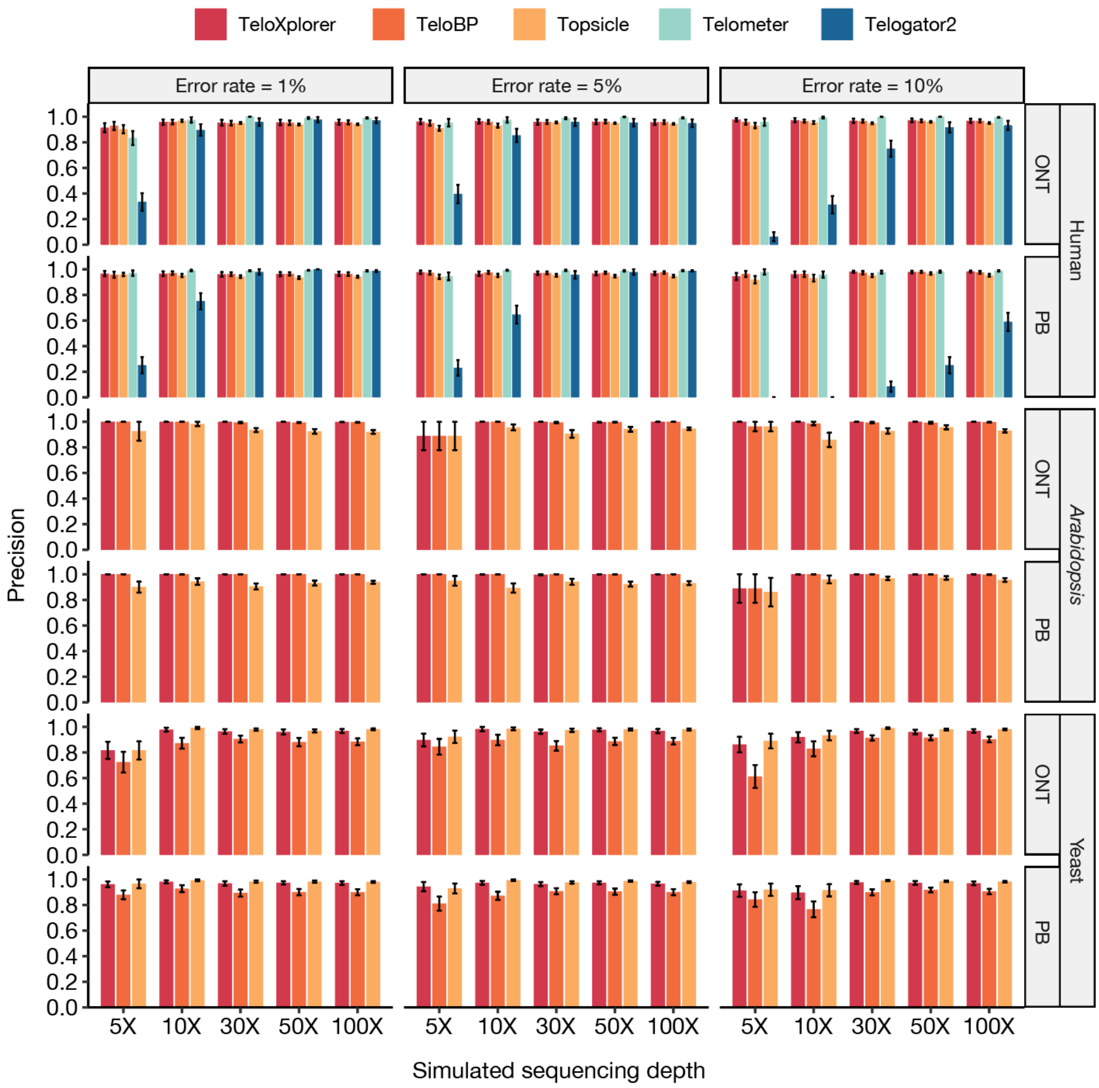


**Figure S2. Precision of identifying chromosome-end-specific telomere containing reads using simulated long-read data.**

Performance of TeloXplorer was evaluated alongside existing tools, including TeloBP, Topsicle, Telometer, and Telogator2, based on simulated PacBio (PB) and Oxford Nanopore Technologies (ONT) reads. Read sets were generated for human, *Arabidopsis*, and yeast across varying sequencing depths (5×–100×) and error rates (1%, 5%, and 10%). Telometer and Telogator2 lack native support for *Arabidopsis* and yeast; hence, performance metrics for these species were not evaluated.

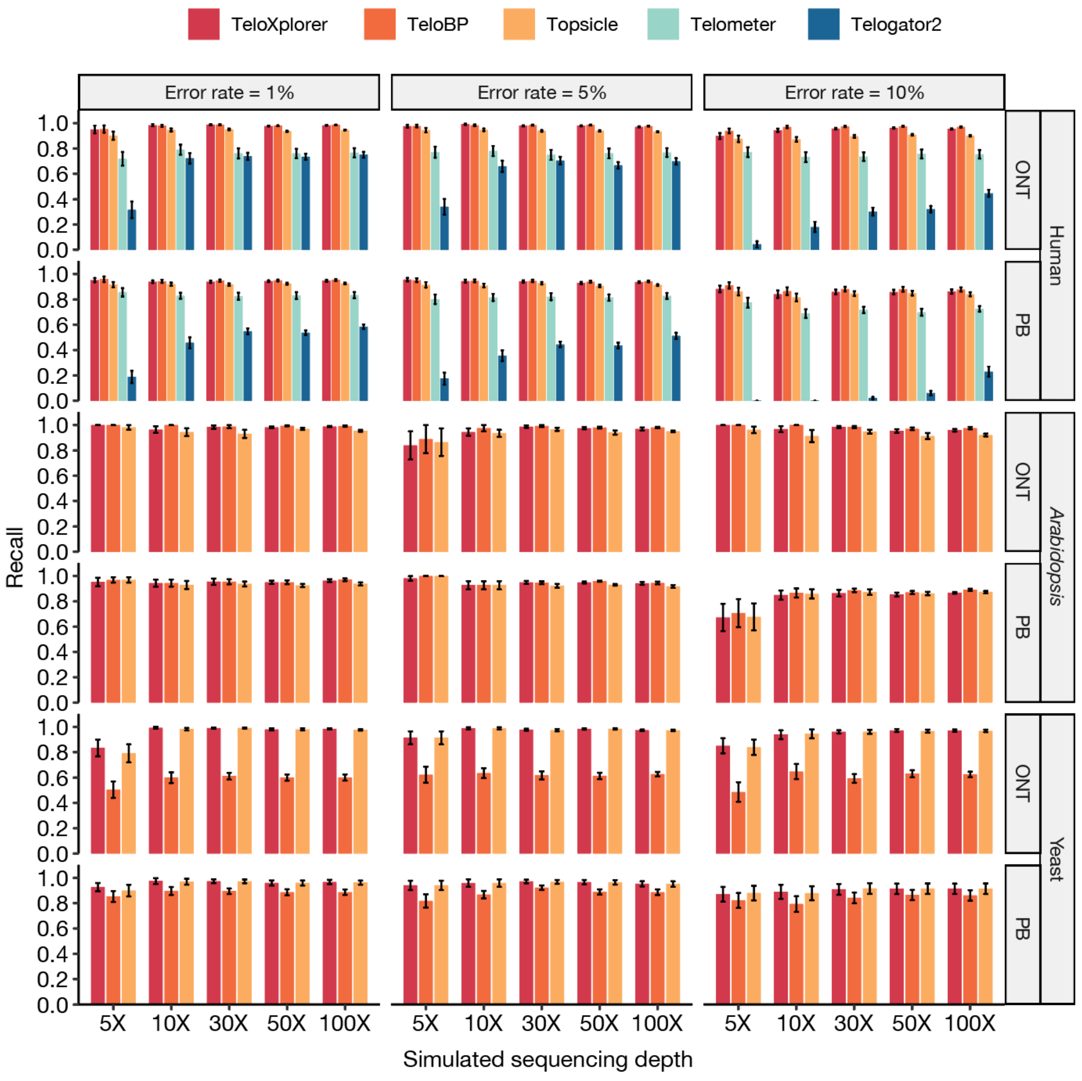


**Figure S3. Recall of identifying chromosome-end-specific telomere containing reads using simulated long-read data.**

Performance of TeloXplorer was evaluated alongside existing tools, including TeloBP, Topsicle, Telometer, and Telogator2, based on simulated PacBio (PB) and Oxford Nanopore Technologies (ONT) reads. Read sets were generated for human, *Arabidopsis*, and yeast across varying sequencing depths (5×–100×) and error rates (1%, 5%, and 10%). Telometer and Telogator2 lack native support for *Arabidopsis* and yeast; hence, performance metrics for these species were not evaluated.

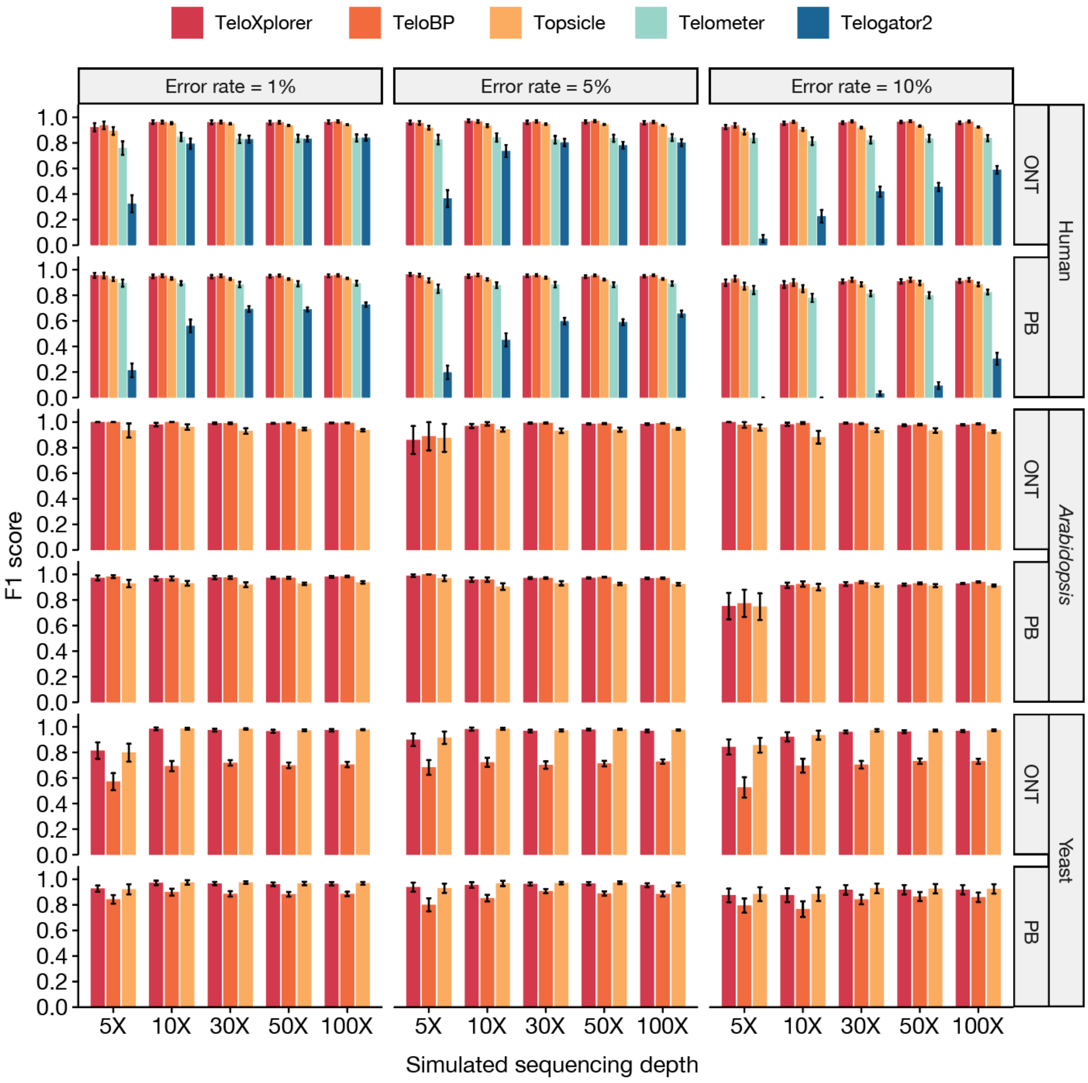


**Figure S4. F1 score of identifying chromosome-end-specific telomere containing reads using simulated long-read data.**

Performance of TeloXplorer was evaluated alongside existing tools, including TeloBP, Topsicle, Telometer, and Telogator2, based on simulated PacBio (PB) and Oxford Nanopore Technologies (ONT) reads. Read sets were generated for human, *Arabidopsis*, and yeast across varying sequencing depths (5×–100×) and error rates (1%, 5%, and 10%). Telometer and Telogator2 lack native support for *Arabidopsis* and yeast; hence, performance metrics for these species were not evaluated.

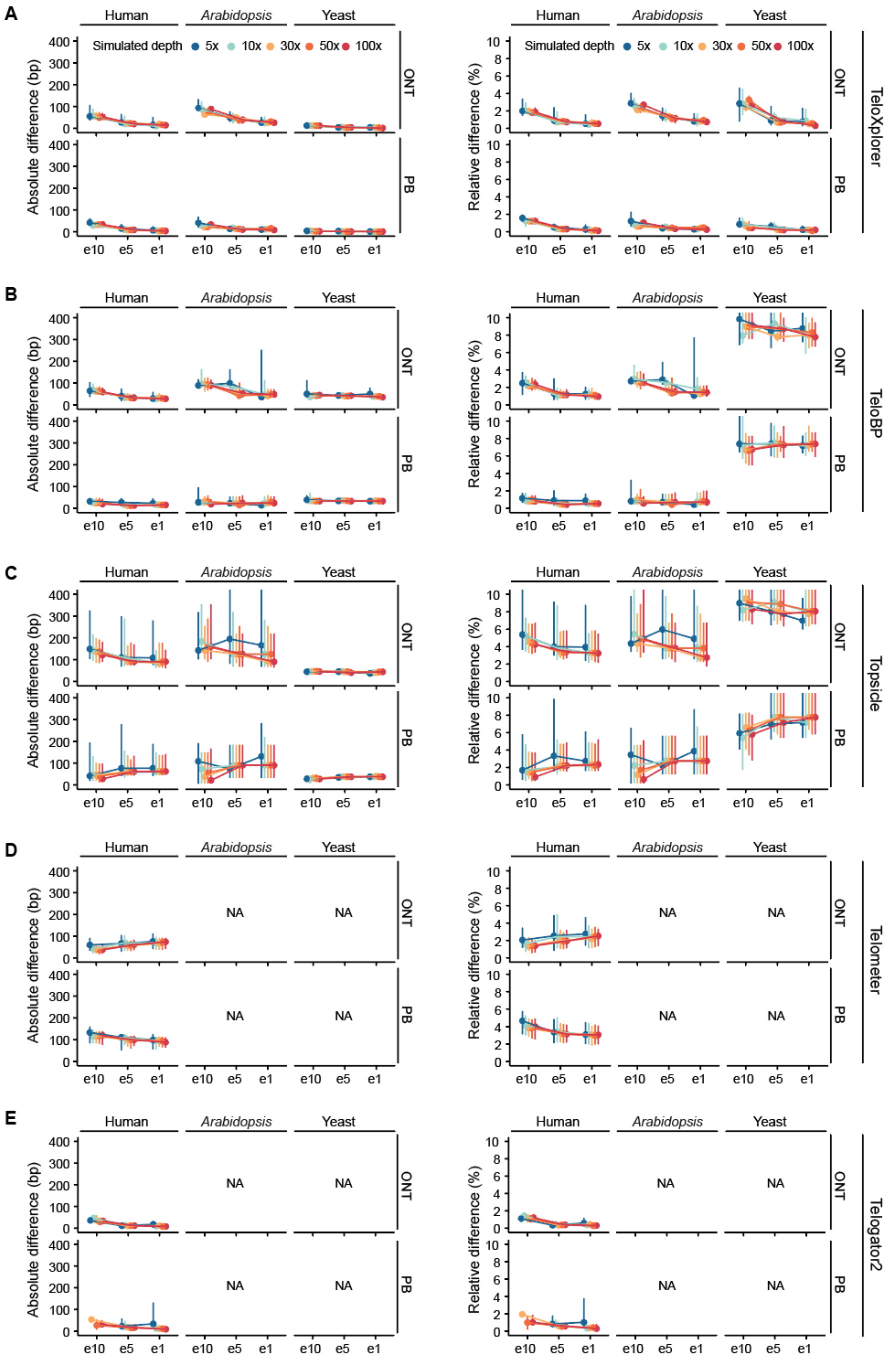


**Figure S5. Benchmarking telomere length estimation by absolute and relative telomere length difference using simulated long-read data.**

Performance of TeloXplorer (A) was evaluated by absolute and relative telomere length difference alongside existing tools, including TeloBP (B), Topsicle (C), Telometer (D), and Telogator2 (E) based on simulated PacBio (PB) and Oxford Nanopore Technologies (ONT) reads. Read sets were generated for human, *Arabidopsis*, and yeast across varying sequencing depths (5×–100×) and error rates (e1: 1%, e5: 5%, and e10: 10%). Telometer and Telogator2 lack native support for *Arabidopsis* and yeast; hence, performance metrics for these species were not evaluated (indicated by NA).

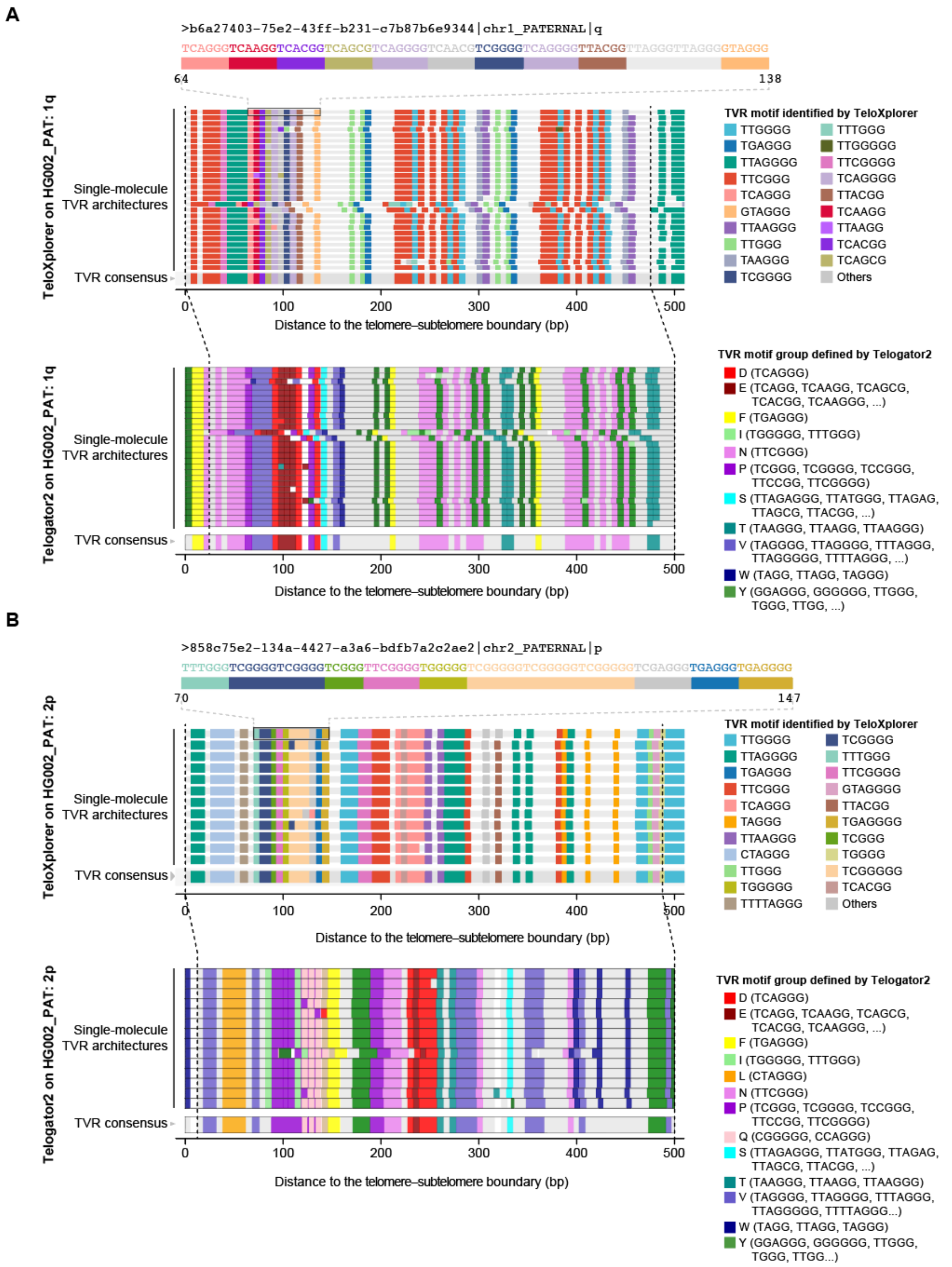


**Figure S6. Comparison of TeloXplorer and Telogator2 for read-level and haplotype-level TVR analysis in human telomeres.**

(A, B) Telomere variant repeat (TVR) maps for the paternal haplotype (PAT) of HG002 chromosome ends 1q (A) and 2p (B), generated using TeloXplorer and Telogator2.

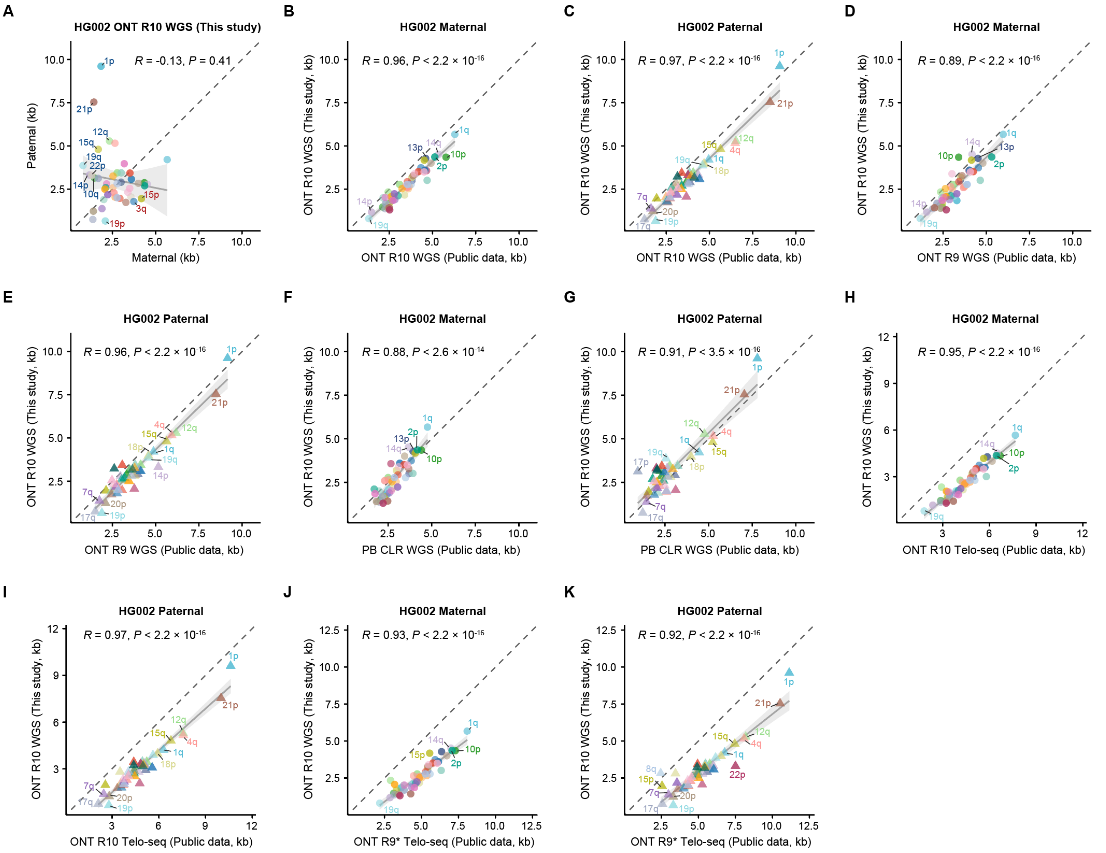


**Figure S7. Correlation of chromosome-end-specific telomere lengths across human HG002 sequencing datasets.**

(A) Correlation between maternal and paternal haplotype-specific telomere length estimates generated in this study via whole-genome sequencing (WGS) using Oxford Nanopore Technologies (ONT) R10 chemistry for the human HG002 cell line. (B, C) Cross-dataset correlation of maternal (B) and paternal (C) telomere length estimates between this study (ONT R10 WGS) and a public ONT R10 WGS dataset for HG002. (D, E) Correlation of maternal (D) and paternal (E) telomere length estimates between this study (ONT R10 WGS) and a public ONT R9 WGS dataset for HG002. (F, G) Correlation of maternal (F) and paternal (G) telomere length estimates for HG002 between this study (ONT R10 WGS) and a public PacBio Continuous Long Read (PB CLR) WGS dataset. (H, I) Correlation of maternal (H) and paternal (I) telomere length estimates for HG002 between this study (ONT R10 WGS) and a public ONT R10 Telo-seq dataset. (J, K) Correlation of maternal (J) and paternal (K) telomere length estimates for HG002 between this study (ONT R10 WGS) and a public ONT R9 Telo-seq dataset featuring retrained basecalling model (denoted by asterisk). Pearson's correlation test was used for all comparison.

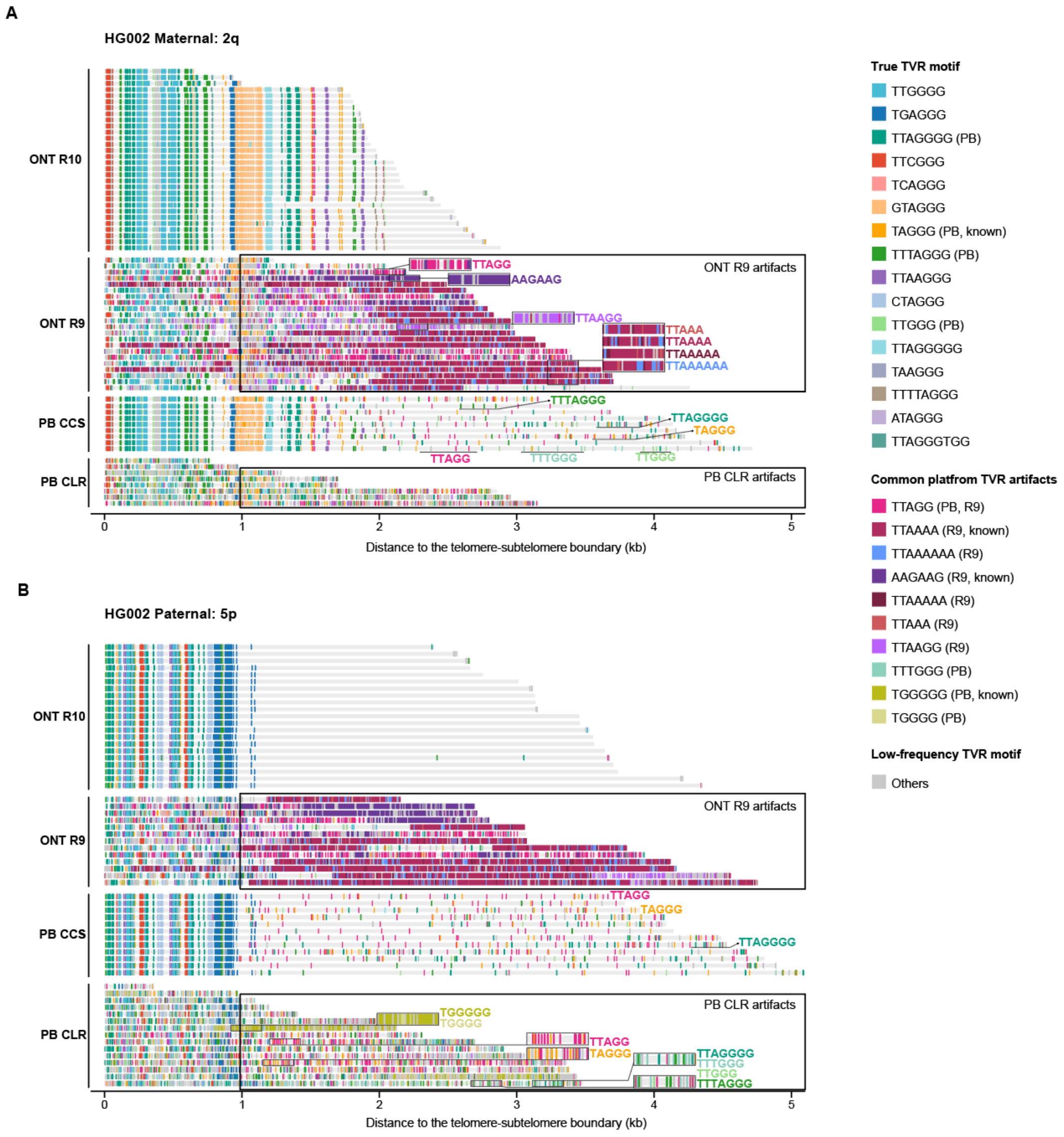


**Figure S8. Cross-platform read-level telomere variant repeat (TVR) profiling of human telomeres.**

(A, B) Comparison of TVR maps generated using Oxford Nanopore Technologies (ONT) R10, ONT R9, PacBio Continuous Long Read (PB CLR), and PB Circular Consensus Sequencing (CCS) platforms for representative HG002 chromosome ends: 2p maternal (A) and 5p paternal (B).

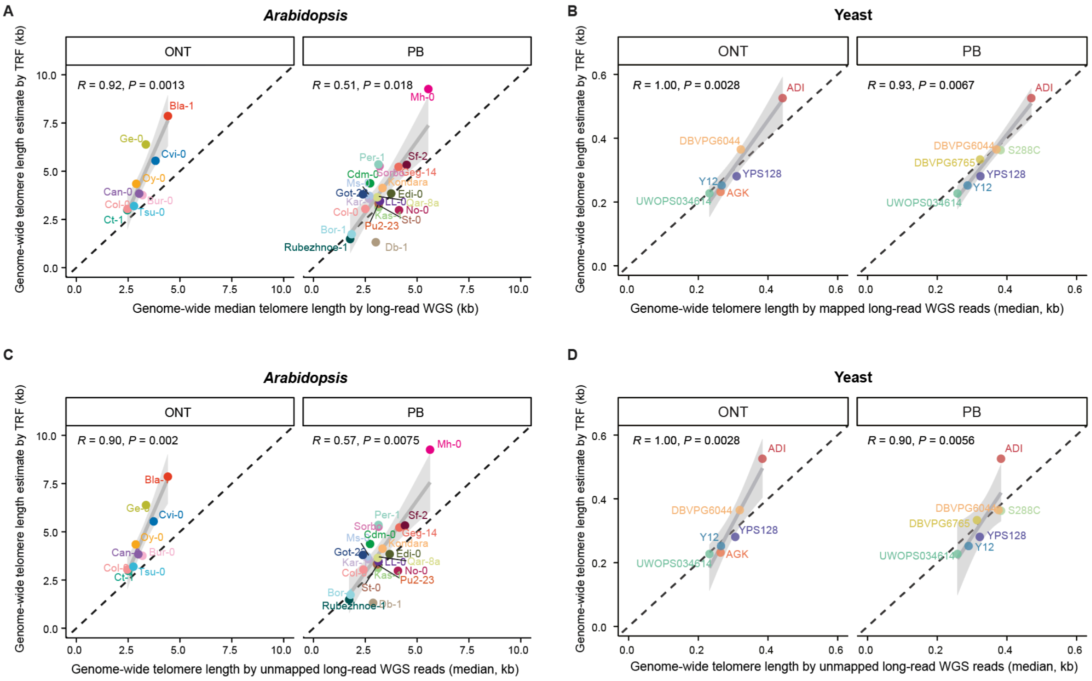


**Figure S9. Genome-wide telomere length comparison between TeloXplorer and terminal restriction fragment (TRF) profiling.**

(A, B) Correlation between TeloXplorer genome-wide estimates (median) with read mapping and genome-wide average telomere lengths determined by TRF across diverse *Arabidopsis thaliana* ecotypes (A) and yeast strains (B). (C, D) Correlation between TeloXplorer genome-wide estimates (median) without read mapping and genome-wide average telomere lengths determined by TRF across diverse *Arabidopsis thaliana* ecotypes (C) and yeast strains (D) Sperman correlation test was used for all comparisons. ONT: Oxford Nanopore Technologies; PB: PacBio.

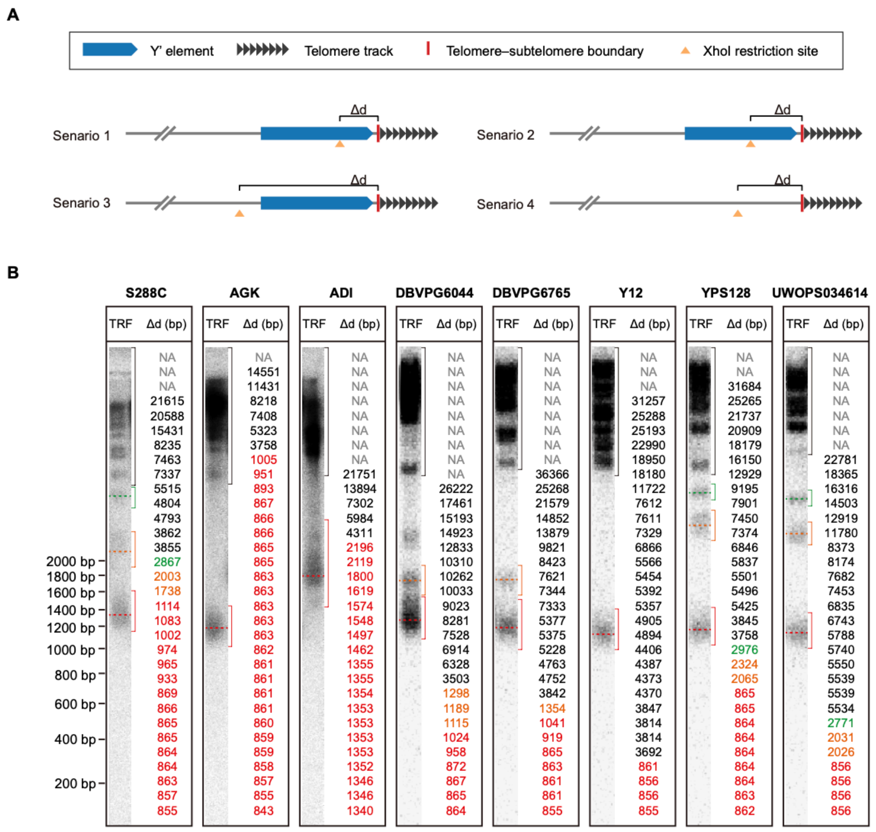


C

| Strain | TRF size[1] (bp) | Mode of Δd[2] (bp) | Adjusted telomere length[3] (bp) |
|---|---|---|---|
| S288C | 1227 | 865 | 362 |
| AGK | 1095 | 863 | 232 |
| ADI | 1879 | 1353 | 526 |
| DBVPG6044 | 1230 | 866 | 363 |
| DBVPG6765 | 1194 | 861 | 333 |
| Y12 | 1108 | 856 | 252 |
| YPS128 | 1145 | 864 | 281 |
| UWOPS034614 | 1083 | 856 | 227 |

Footnote:

1. D'Angiolo, et al. (2023) Genetics.
2. At a given chromosome end, Δd represents the distance from the telomere boundary to the proximal XhoI restriction site.
3. Adjusted telomere length = TRF size - Mode of Δd.

**Figure S10. Adjusting yeast telomere length estimates from Telomere Restriction Fragment (TRF) data by accounting for subtelomeric variation.**

(A) Structural architecture of the yeast *Saccharomyces cerevisiae* chromosome ends under different subtelomeric configurations. Representative scenarios 1–3 contain Y′ elements, whereas scenario 4 represents a Y′-free chromosome end. $\Delta d$ denotes the distance between the telomere–subtelomere boundary and the proximal XhoI site. (B) Re-assessment of published TRF data (D'Angiolo et al. 2023) for eight representative yeast strains by considering their respective genome-assembly-derived $\Delta d$ values. For each strain, the sorted $\Delta d$ values across all chromosome ends are displayed adjacent to their corresponding TRF bands (color-coded), with the modal $\Delta d$ values highlighted in bold. (C) Adjusted telomere lengths for representative yeast strains, calculated by subtracting the mode of $\Delta d$ across all chromosome ends in a given strain from the measured TRF band size.

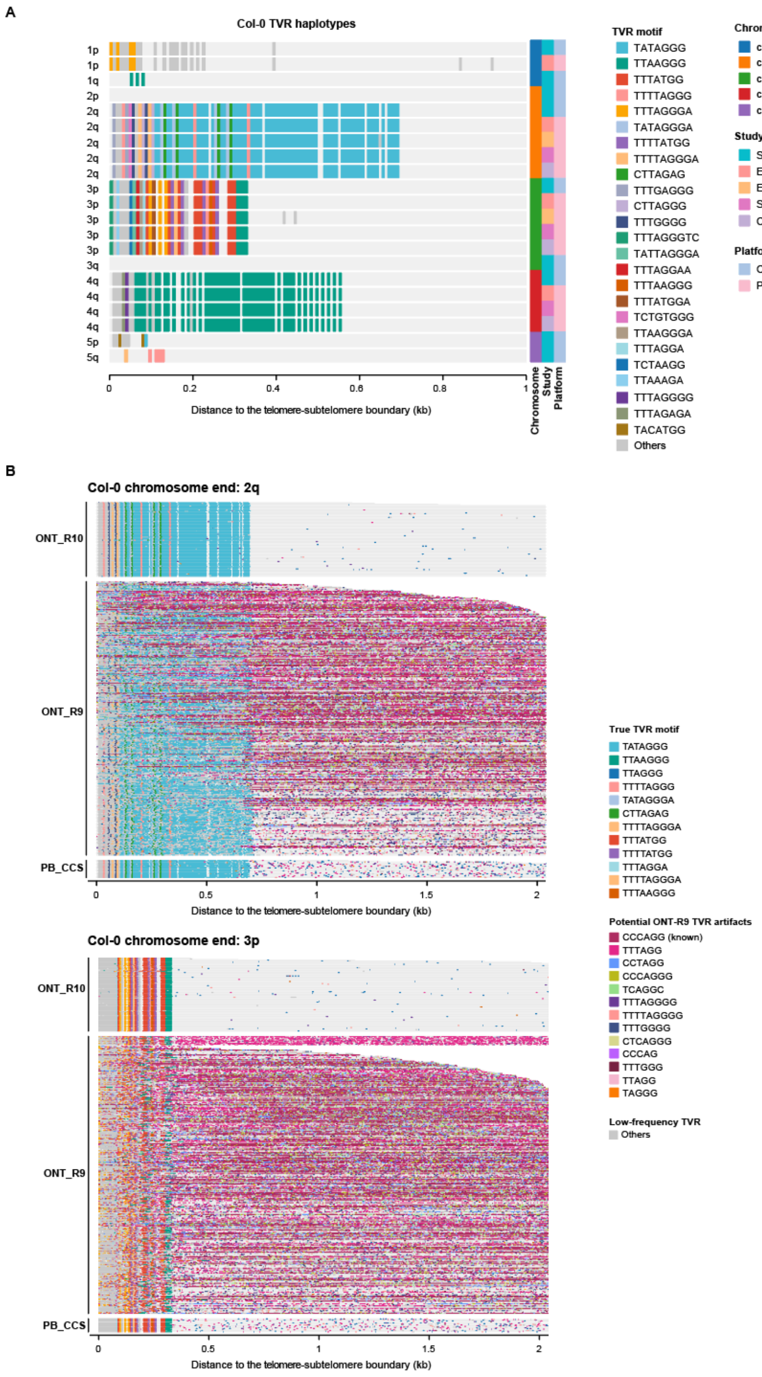


**Figure S11. Cross-platform read-level telomere variant repeat (TVR) profiling of *Arabidopsis* telomeres.**

(A) Comparison of haplotype-level TVR maps for the *Arabidopsis* Col-0 ecotype generated using Oxford Nanopore Technologies (ONT) R10 and PacBio Circular Consensus Sequencing (PB CCS) platforms for all accessible chromosome ends. (B) Comparison of read-level TVR maps for *Arabidopsis* Col-0 ecotype for representative chromosome ends: 2q and 3p.

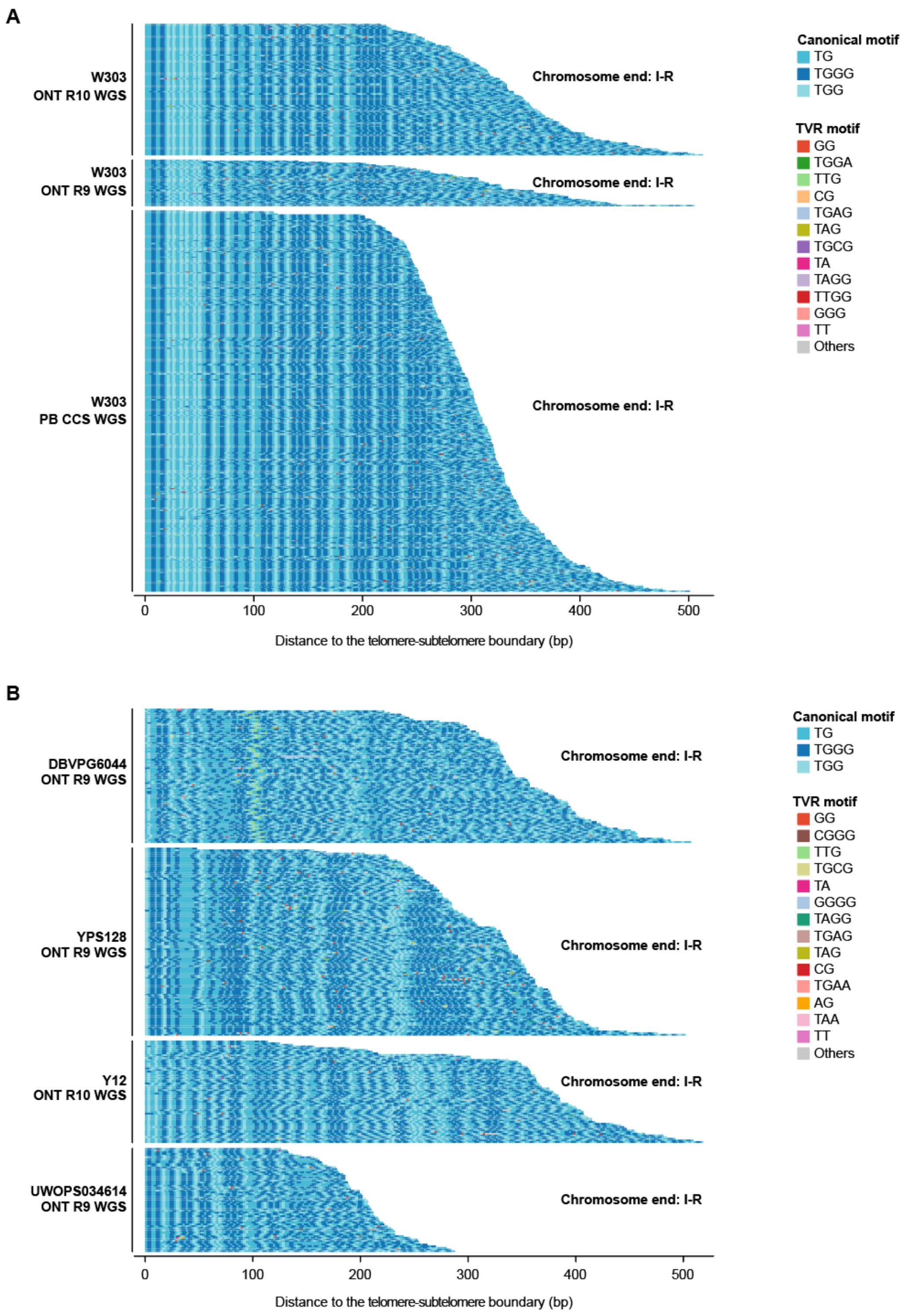


**Figure S12. Cross-platform read-level telomere variant repeat (TVR) profiling of yeast telomeres.**

(A) Comparison of read-level TVR maps for the yeast W303 strain generated using Oxford Nanopore Technologies (ONT) R10, ONT R9, and PacBio Circular Consensus Sequencing (PB CCS) platforms for a representative chromosome end: I-right (I-R). (B) Comparison of read-level TVR maps across diverse yeast strains generated using ONT R10 and R9 platforms for the chromosome end I-R.

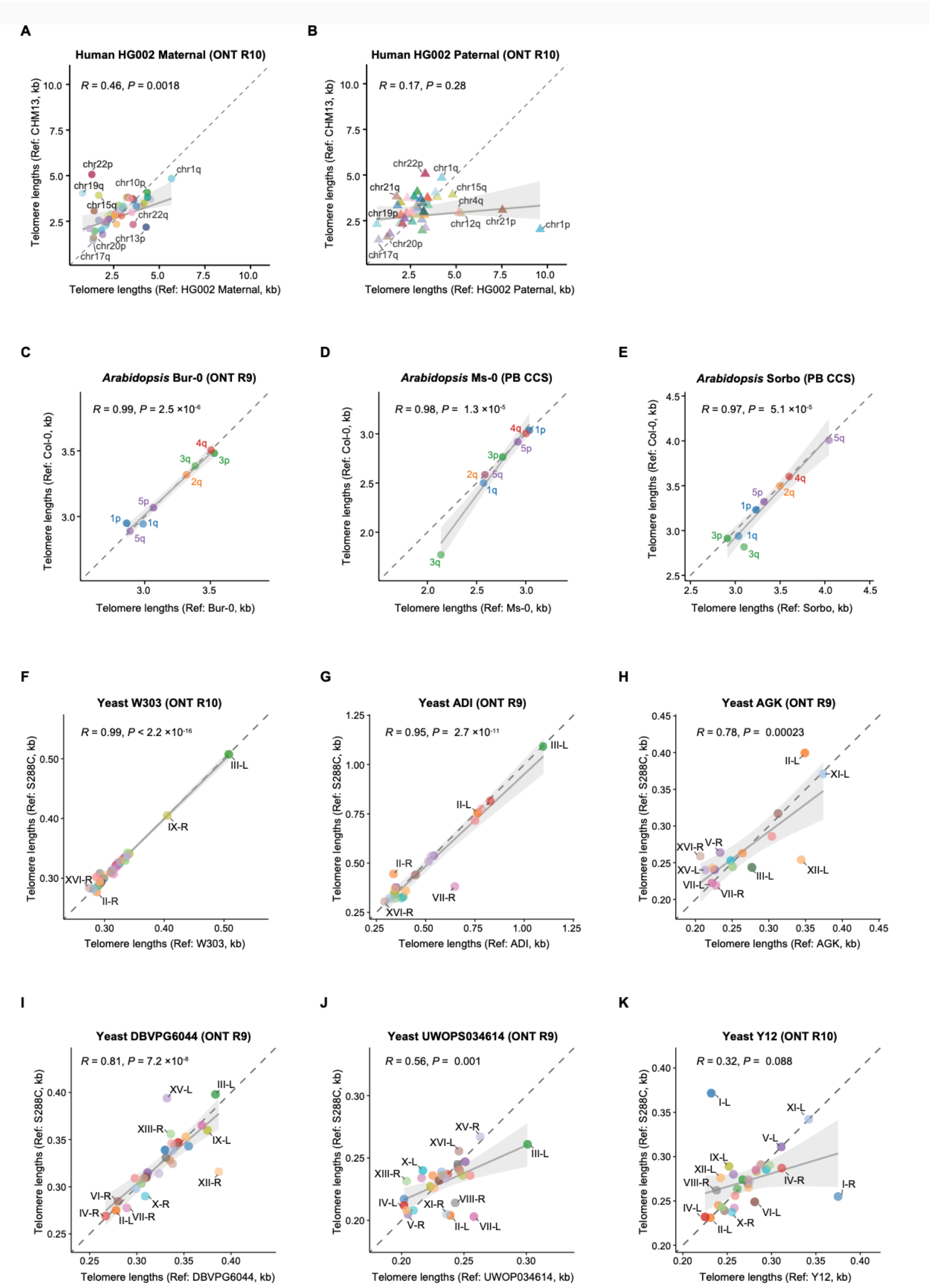


**Figure S13. Comparison of chromosome-end-specific telomere length estimates using native versus generic reference genomes.**

(A, B) Correlation of the human HG002 maternal (A) and paternal (B) telomere length estimates derived using native haplotype-resolved assemblies versus the generic CHM13 reference genome. (C–E) Correlation of *Arabidopsis* telomere length estimates derived using the native assemblies versus the generic Col-0 reference genome for the ecotypes Bur-0 (C), Ms-0 (D), and Sorbo (E). (F–K) Correlation of yeast telomere length estimates derived using the native assemblies versus the generic S288C reference genome for the strains W303 (F), ADI (G), AGK (H), DBVPG6044 (I), UWOPS034614 (J), Y12 (K).

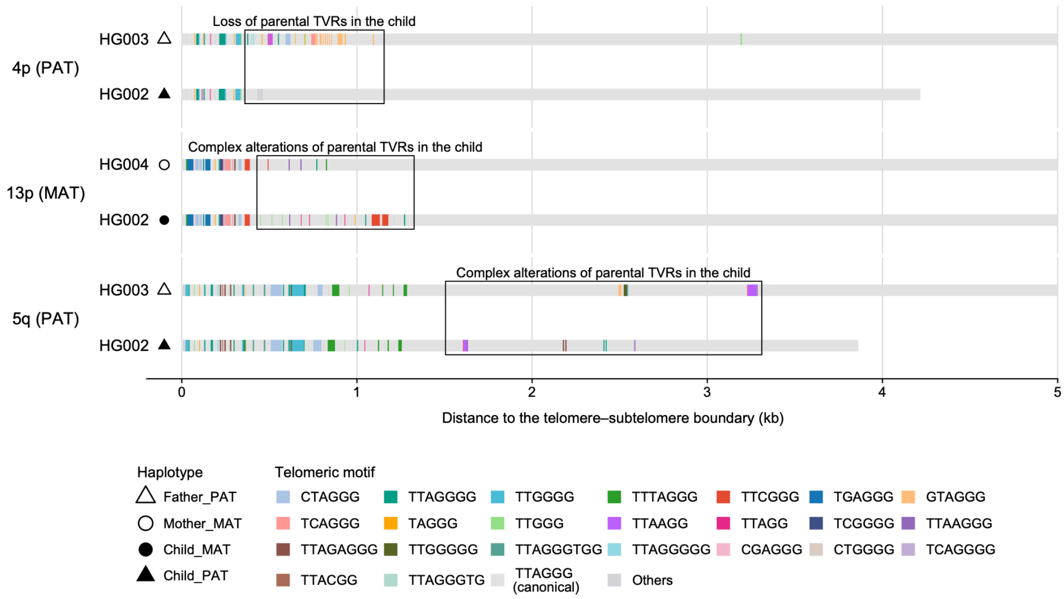


**Figure S14. Representative examples of TVR mutations in the child (HG002) in comparison to his parents (father: HG003, mother: HG004).**
Parental origin of TVR haplotypes in the child (HG005; maternal alleles marked by filled circles, paternal alleles by filled triangles) aligned with matching haplotypes from the mother (HG007; open circles) and father (HG006; open triangles). Canonical telomere repeats are shown in light gray as the background, with colored stripes representing variant repeats (TVRs). MAT: maternal haplotype; PAT: paternal haplotype.

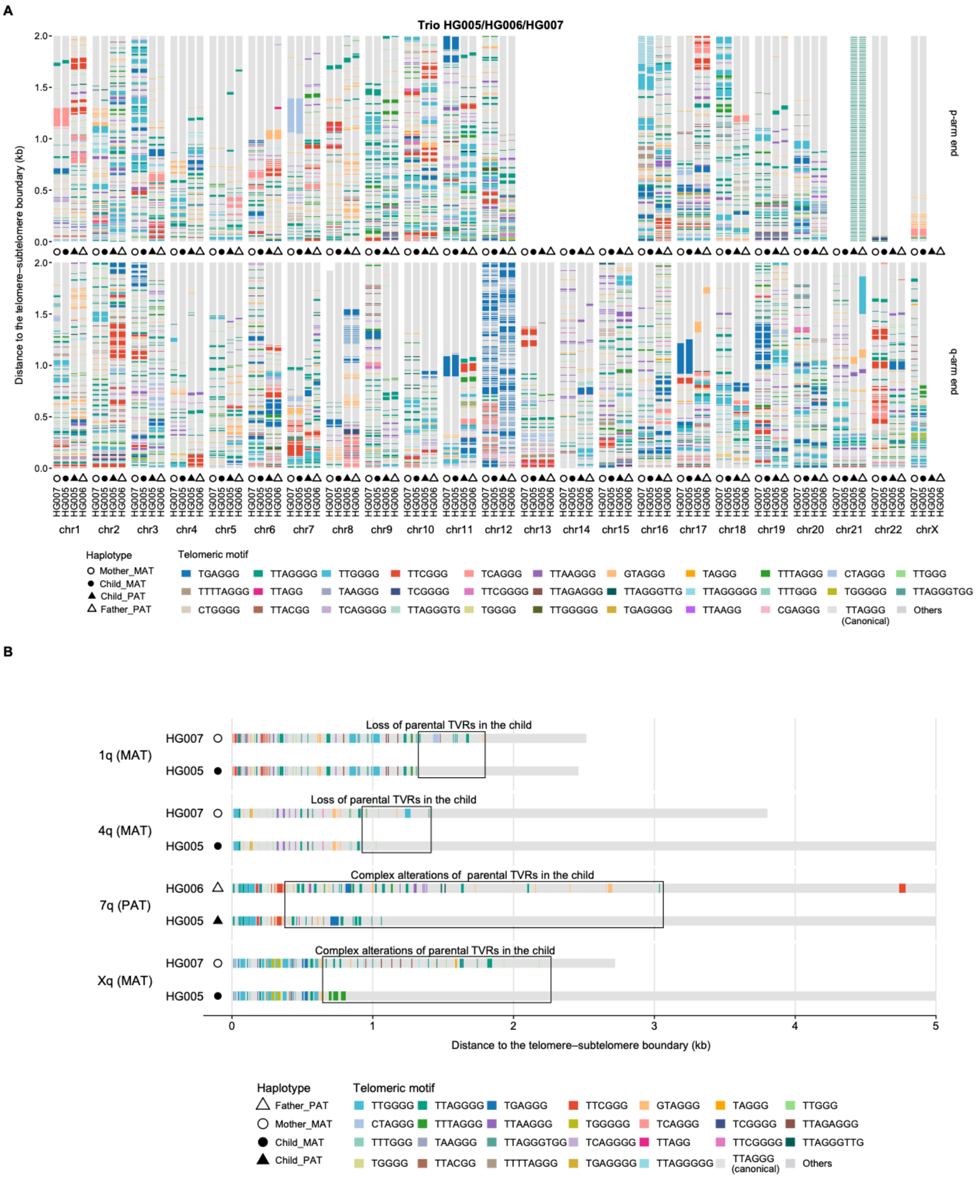

**Figure S15. Allele-specific inheritance of telomere variant repeat (TVR) composition across the HG005/HG006/HG007 trio.**

(A) Parental origin of TVR haplotypes in the child (HG005; maternal alleles marked by filled circles, paternal alleles by filled triangles) aligned with matching haplotypes from the mother (HG007; open circles) and father (HG006; open triangles). Canonical telomere repeats are shown in light gray as the background, with colored stripes representing variant repeats (TVRs). (B) Representative examples of TVR mutations in the child (HG005) in comparison to his parents (father: HG006, mother: HG007). MAT: maternal haplotype; PAT: paternal haplotype.

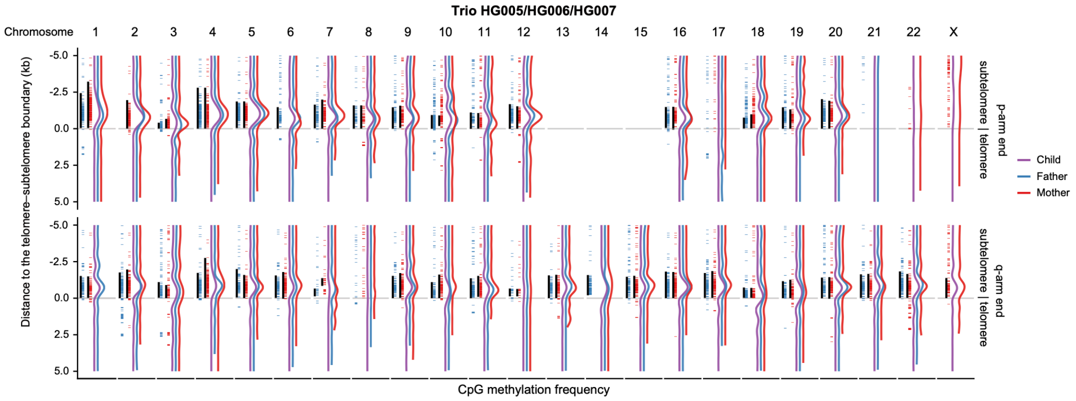


**Figure S16. Allele-specific inheritance of telomeric and subtelomeric DNA methylation across the HG005/HG006/HG007 trio.**

Chromosomal distribution of DNA methylation patterns across the HG005/HG006/HG007 trio (child: purple; mother: red; father: blue), shown alongside annotations for CpG clusters (blue and red stripes) and TAR1 elements (black bars) of the corresponding maternal (red) and paternal (blue) homologous chromosomes.

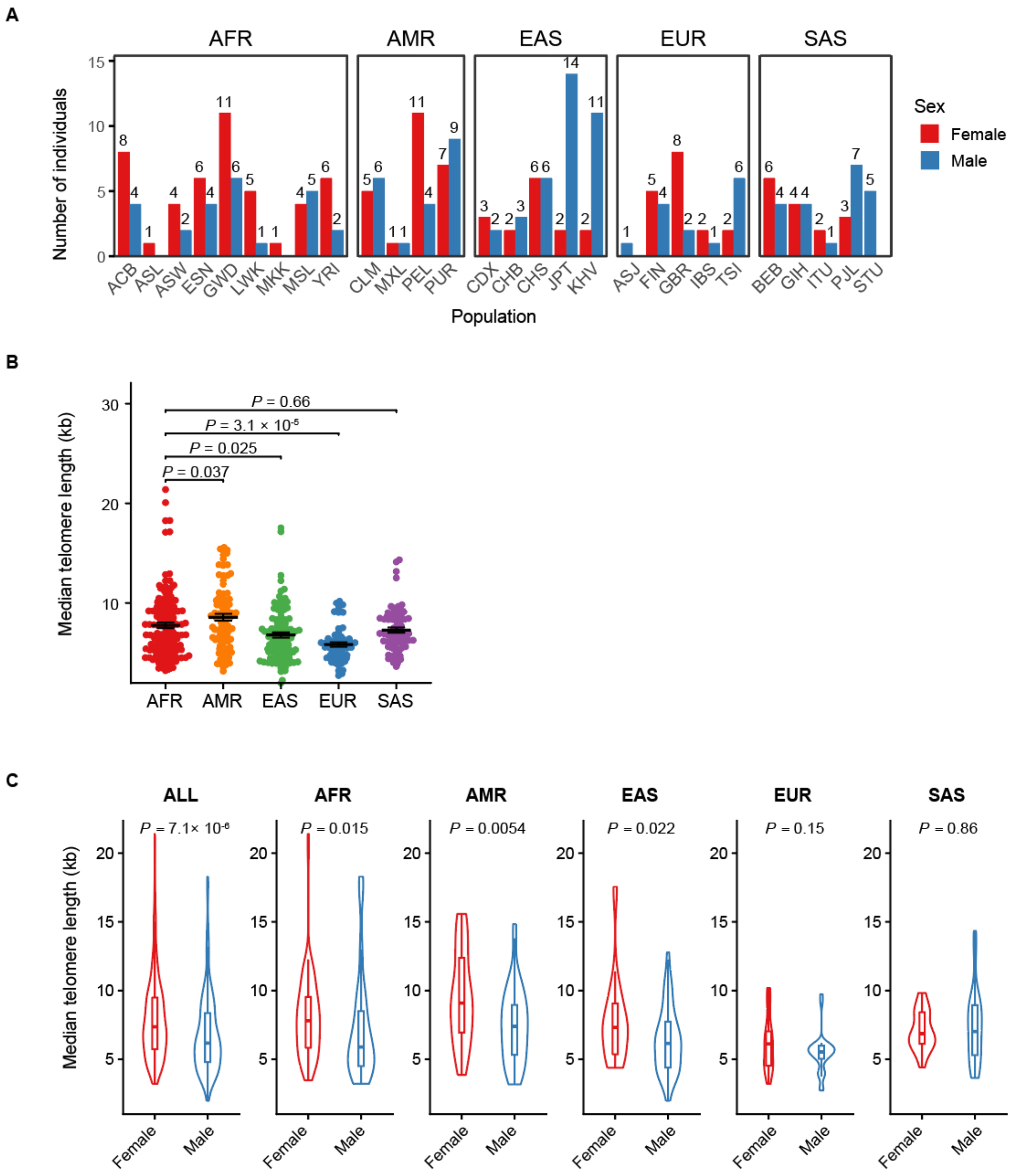


**Figure S17. Telomere length distribution of the HPRC2 cohort.**
(A) Continental, population and sex decomposition of the HPRC2 cohort. (B) Genome-wide median telomere lengths across different continental groups. (C) Comparison of genome-wide median telomere lengths between females and males within each continental group. Continental groups: AFR, African; AMR, Admixed American; EAS, East Asian; EUR, European; SAS, South Asian. *P*-values were calculated using two-sided Wilcoxon rank-sum tests for all comparisons.

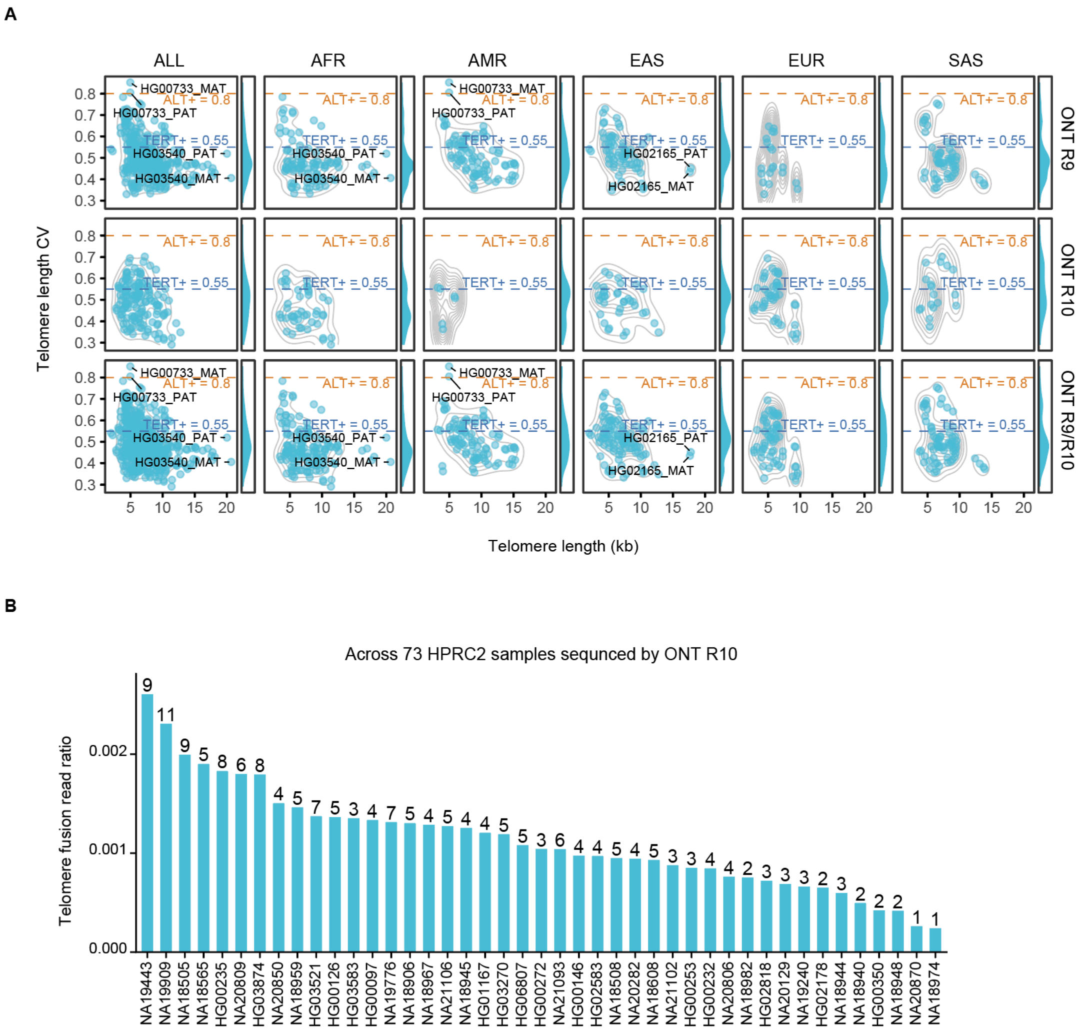


**Figure S18. Screening for Alternative Lengthening of Telomeres (ALT) candidates across the HPRC2 cohort.**

(A) Bivariate distribution of telomere length and heterogeneity (coefficient of variation, CV) across the HPRC2 individuals sequenced using Oxford Nanopore Technologies (ONT R9 vs. R10). Dashed reference lines denote previously reported characteristic CV value for telomerase-positive (TERT+, CV = 0.55) and ALT-positive (ALT+, CV = 0.8) mechanisms (Schmidt, et al. 2024). Continental groups: AFR, African; AMR, Admixed American; EAS, East Asian; EUR, European; SAS, South Asian. (B) Proportion of reads containing telomere fusion signatures (an ALT+ signature) across 73 HPRC2 individuals sequenced on the ONT R10 platform.

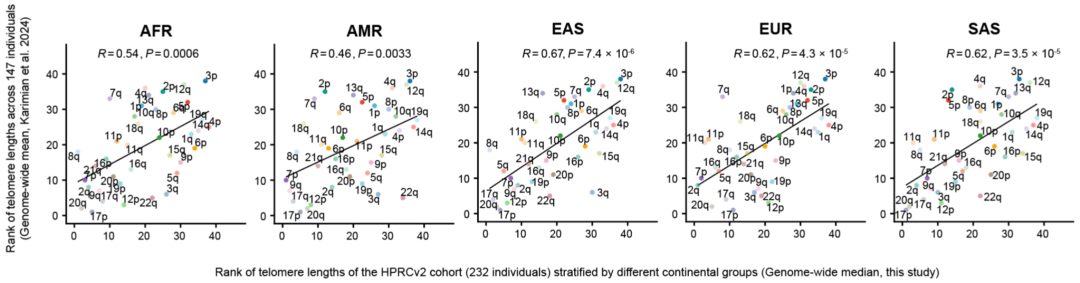


**Figure S19. Cross-cohort consistency of chromosome-end-specific telomere length rankings.**

Spearman rank correlation of chromosome-end-specific telomere lengths between HPRC2 continental groups (AFR, African; AMR, Admixed American; EAS, East Asian; EUR, European; SAS, South Asian) and a previously published independent cohort (Karimian et al. 2024). *P*-values indicate two-sided statistical significance.

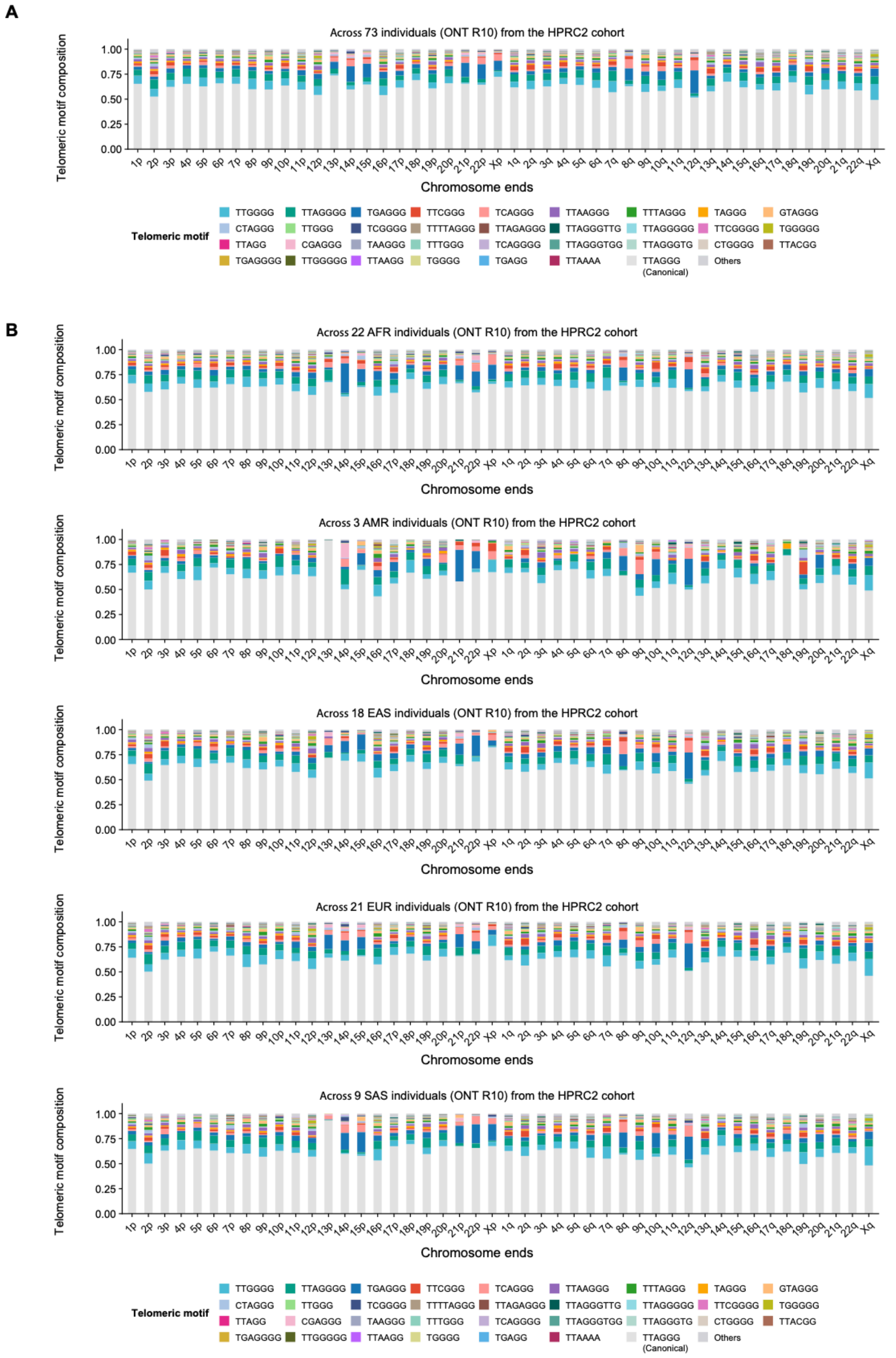


**Figure S20. Chromosome-end-specific telomeric motif composition across the HPRC2 cohort.**

(A) Global profile of telomeric motif composition across all 73 ONT R10-sequenced HPRC2 individuals. (B) telomeric motif composition across the same cohort, stratified by continental groups (AFR, African; AMR, Admixed American; EAS, East Asian; EUR, European; SAS, South Asian).

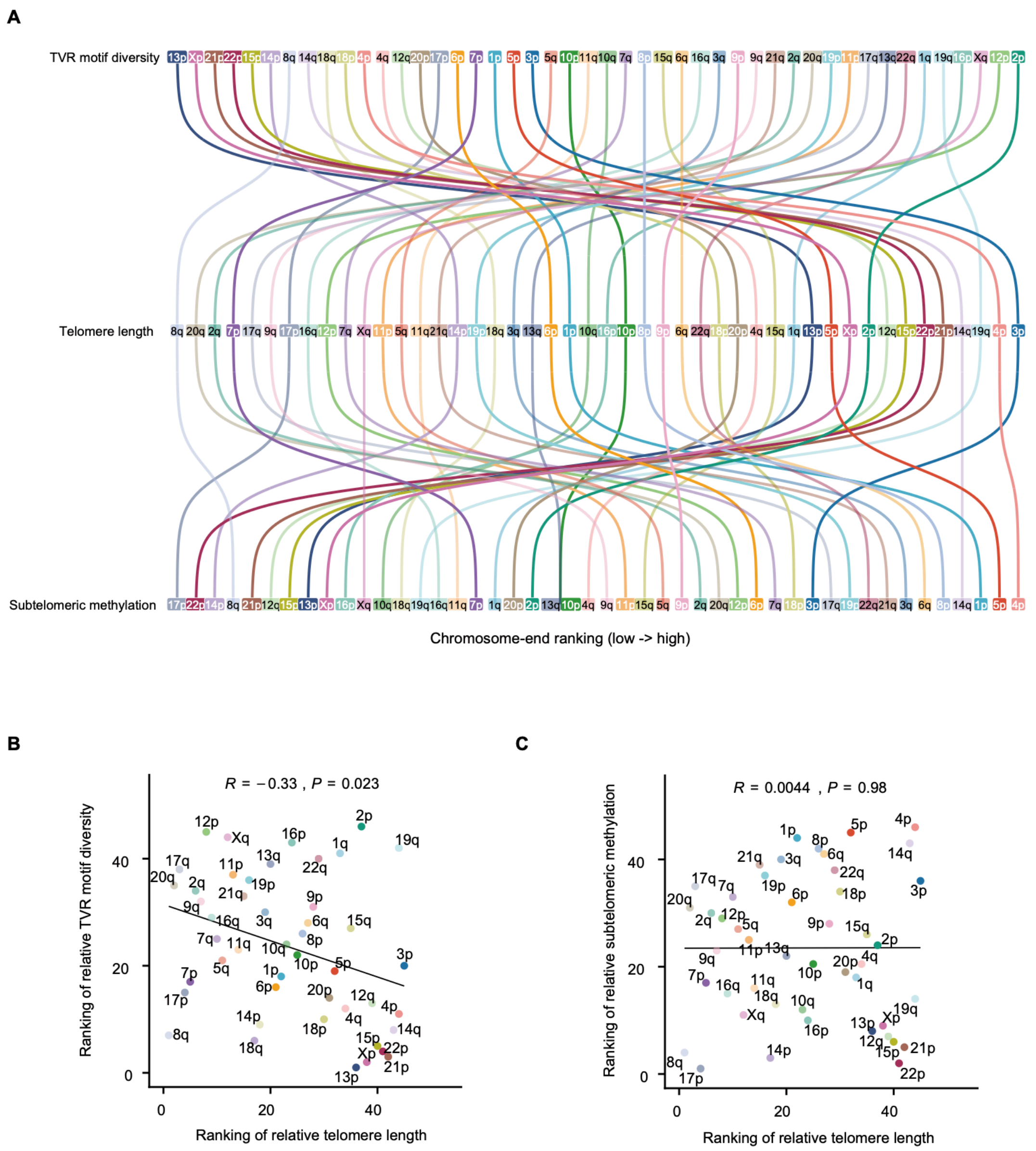

**Figure S21. Interrelationships among chromosome-end-specific telomere length, telomere variant repeat (TVR) motif diversity, and subtelomeric methylation in the HPRC2 cohort.**

(A) Chromosome-end-specific ranking of telomere length, TVR motif diversity, and subtelomeric DNA methylation levels. (B, C) Spearman rank correlations of chromosome-end-specific rankings between telomere length and TVR motif diversity (B) as well as subtelomeric methylation level (C) respectively. *P*-values indicate two-sided statistical significance.

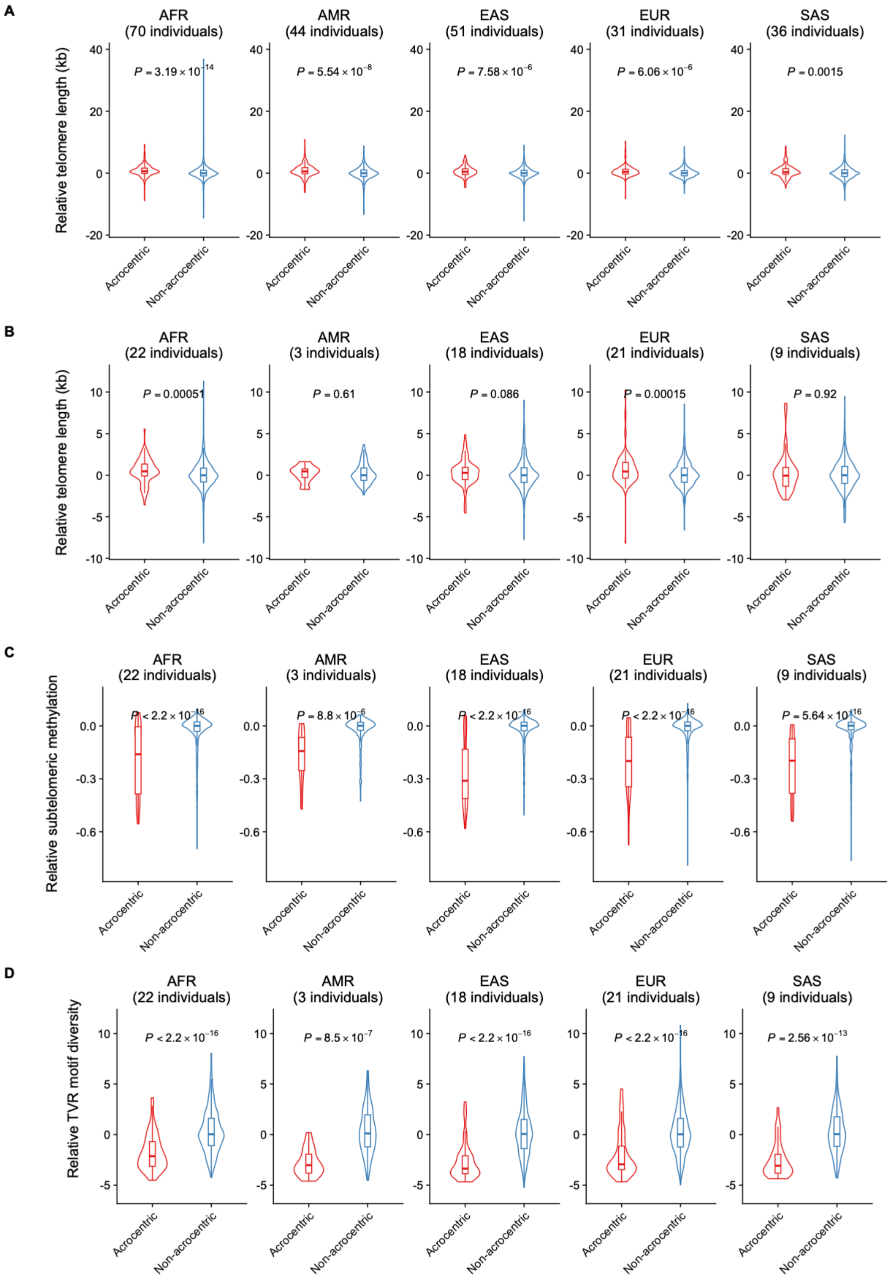


**Figure S22. Comparison of Telomeric and subtelomeric features between acrocentric and non-acrocentric chromosome ends in the HPRC2 cohort.**

(A) Relative telomere length comparison between acrocentric and non-acrocentric chromosome ends across the full HPRC2 cohort (232 individuals). (B) Comparison of relative telomere length, telomere variant repeat (TVR) motif diversity, and subtelomeric DNA methylation levels between acrocentric and non-acrocentric ends across the Oxford Nanopore Technologies (ONT) R10-sequenced HPRC2 subset (73 individuals). *P*-values were calculated using two-sided Wilcoxon rank-sum tests for all comparisons. Continental groups: AFR, African; AMR, Admixed American; EAS, East Asian; EUR, European; SAS, South Asian.

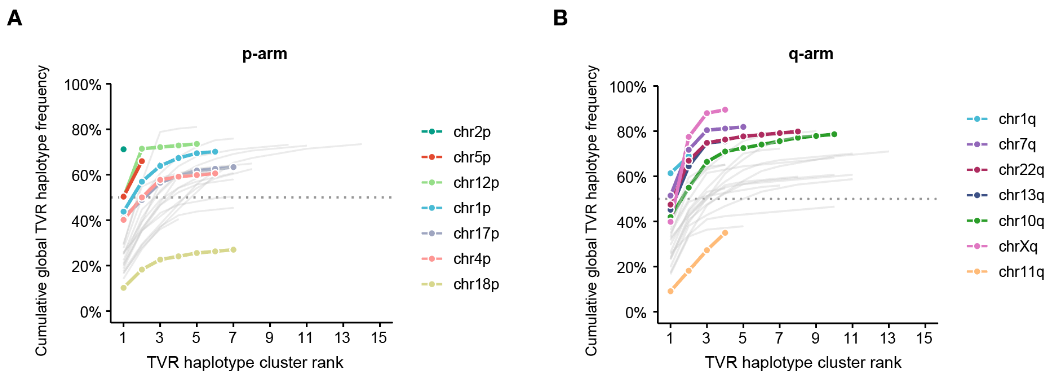


**Figure S23. Cumulative global frequencies of chromosome-end-specific telomere variant repeat (TVR) haplotype cluster in the HPRC2 cohort.**

Relationship between chromosome-end-specific telomere variant repeat (TVR) haplotype cluster ranks (ordered by cluster frequency) and their respective cumulative global frequencies. (A, B) Frequency distribution profiles shown separately for p-arm (A) and q-arm (B) chromosome ends. Horizontal dotted lines indicate the 50% cumulative TVR haplotype frequency reference. This analysis was conducted on the HPRC2 subset of 73 individuals sequenced using the Oxford Nanopore Technologies (ONT) R10 platform.

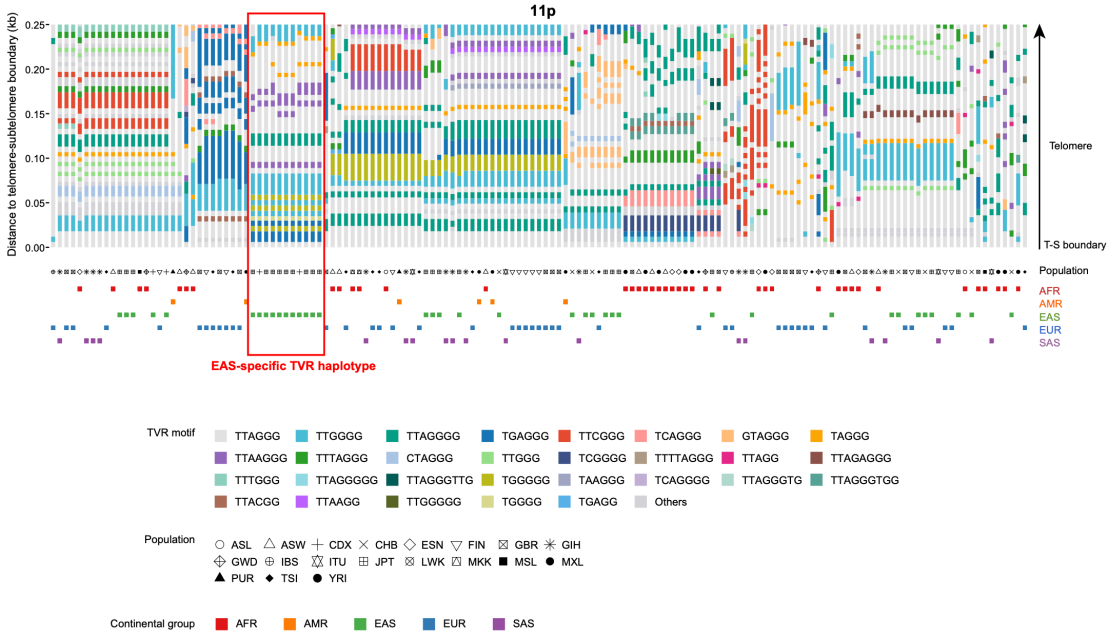


**Figure S24. Haplotype-resolved telomere variant repeat (TVR) profile for the HPRC2 cohort.**

Individual haplotypes of telomere across 73 individuals from the HPRC cohort sequenced using the Oxford Nanopore Technologies (ONT) R10 platform were analyzed and presented in an ordered fashion based their TVR haplotype similarity. Sample metadata, including continental group (AFR, African; AMR, Admixed American; EAS, East Asian; EUR, European; SAS, South Asian) and detailed population of origin, are annotated. The red outline highlights a prominent EAS-specific TVR haplotype cluster. T-S boundary: The telomere–subtelomere boundary.

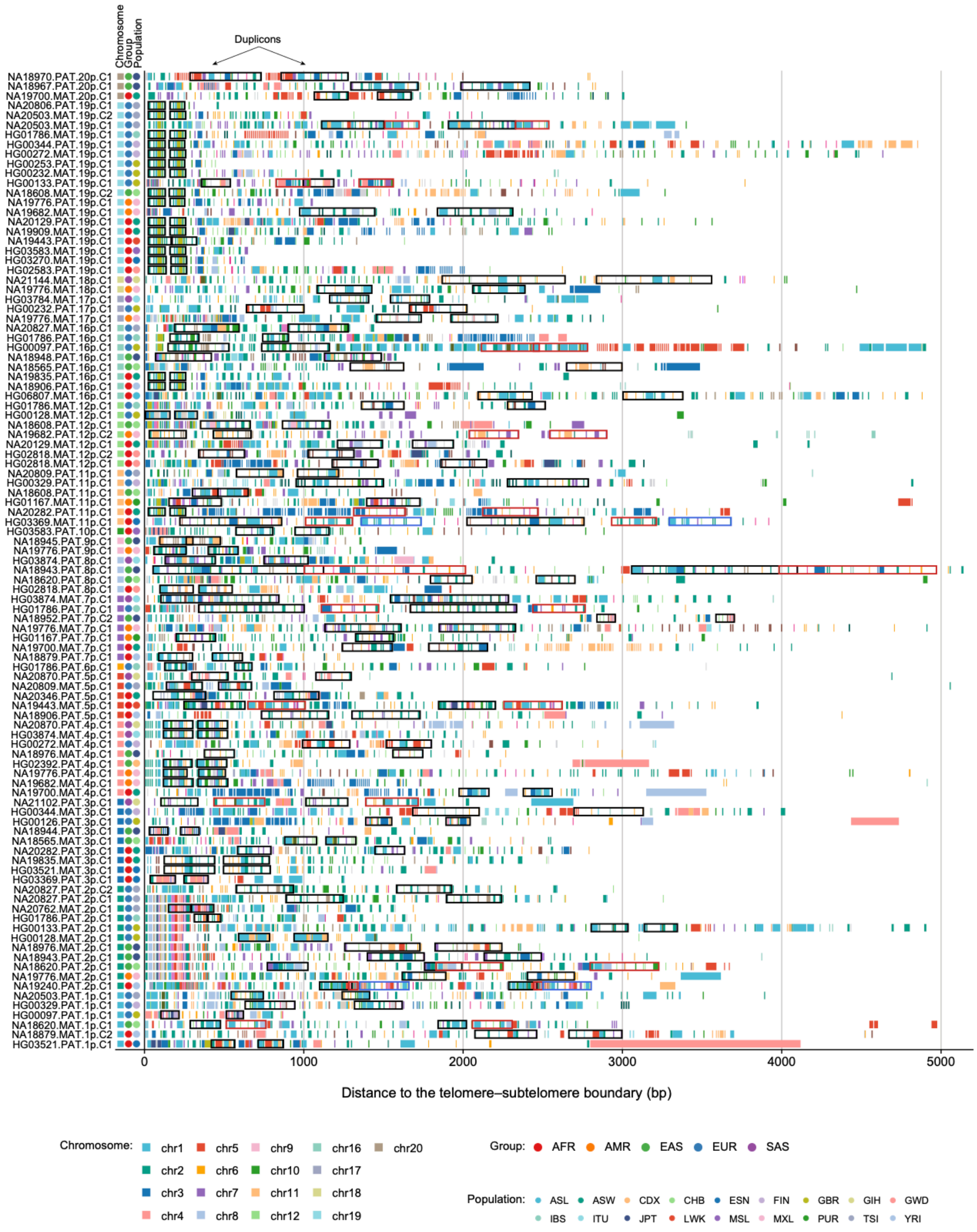


**Figure S25. Duplication blocks of telomere variant repeat (TVR) identified in the p-arm telomeres across the HPRC2 cohort.**

Haplotype-resolved TVR architectures containing identified TVR duplication blocks were are presented. Haplotype identifiers are formatted as: <HPRC2_Sample_ID>.<Maternal (MAT)/Paternal (PAT) haplotype>. <chromosome end>. < TVR haplotype ID (C1/C2)>.

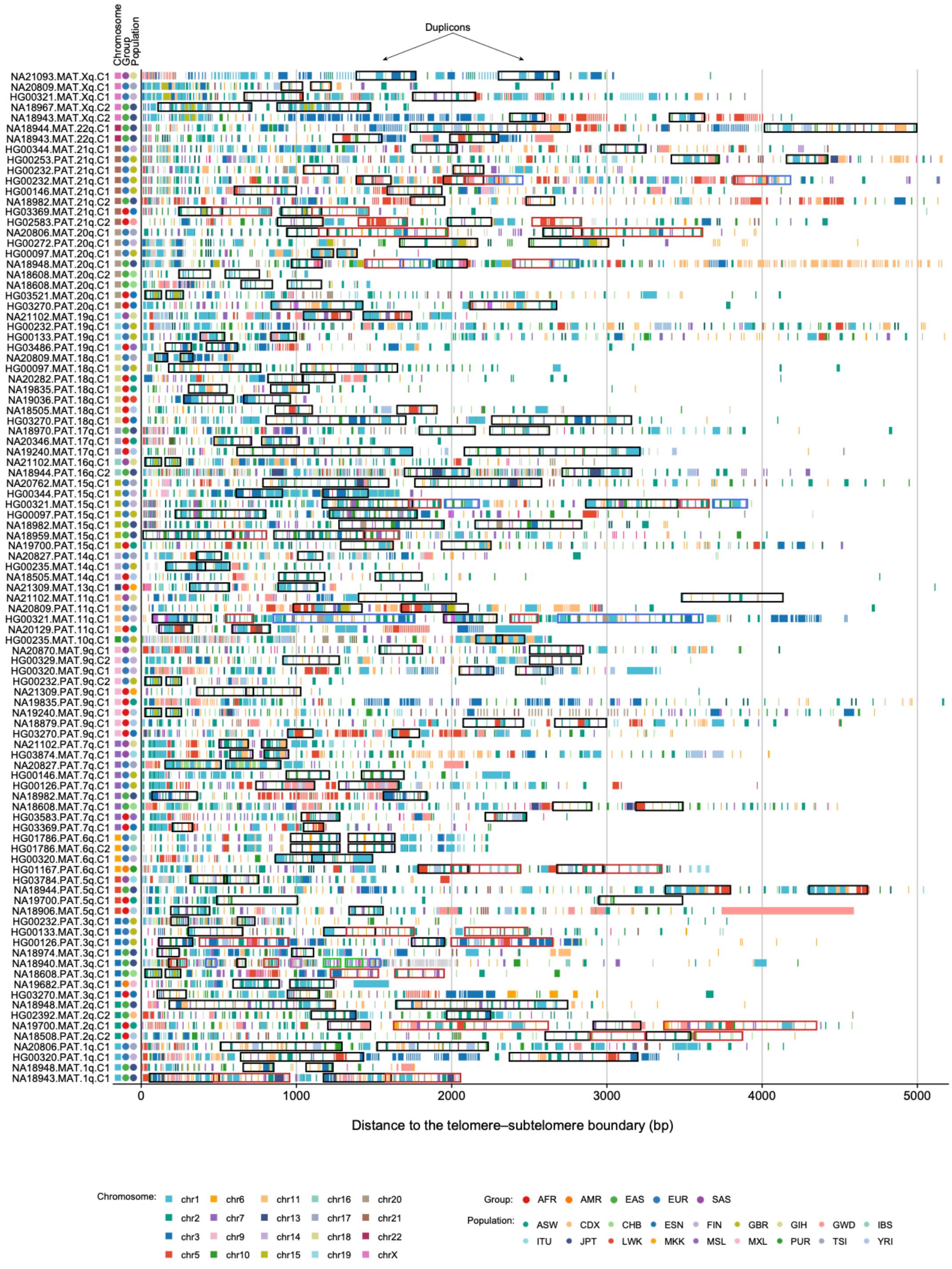


**Figure S26. Duplication blocks of telomere variant repeat (TVR) identified in the q-arm telomeres across the HPRC2 cohort.**

Haplotype-resolved TVR architectures containing identified TVR duplication blocks were are presented. Haplotype identifiers are formatted as: <HPRC2_Sample_ID>.<Maternal (MAT)/Paternal (PAT) haplotype>. <chromosome end>. < TVR haplotype ID (C1/C2)>.

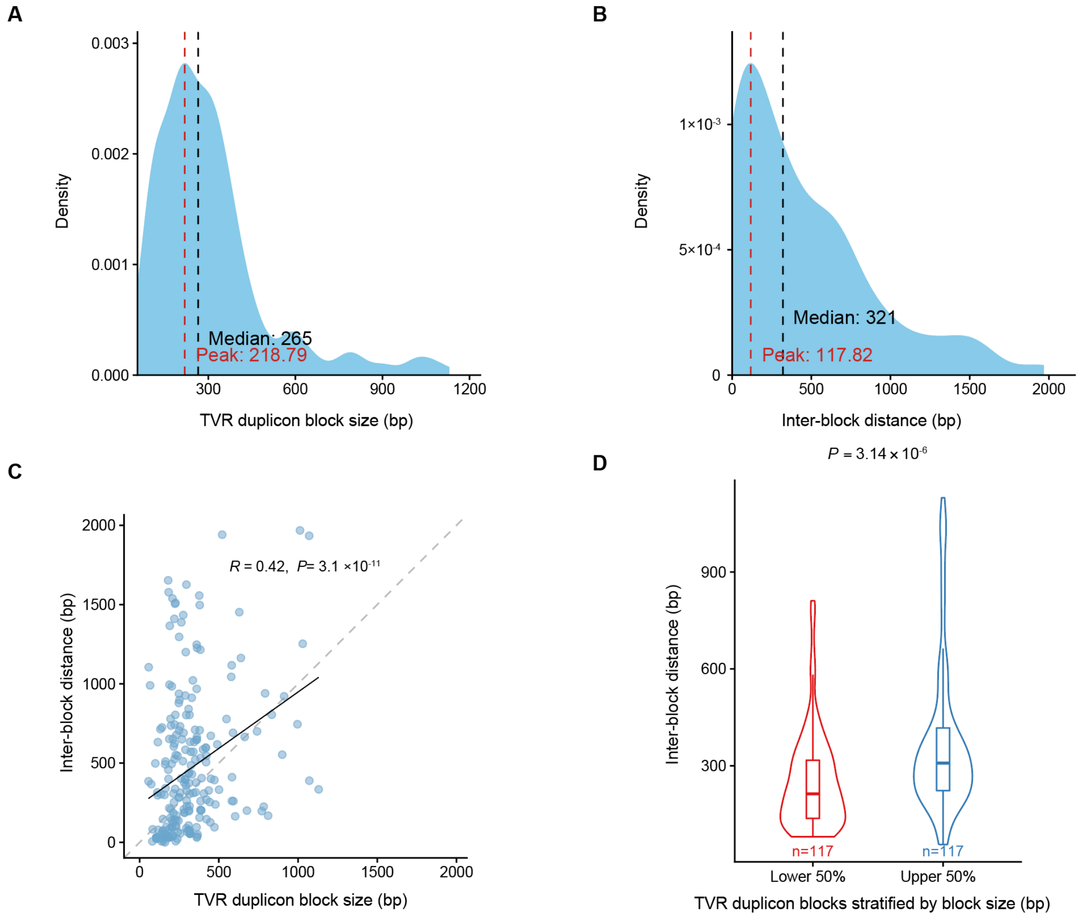


**Figure S27. Structural characteristics of telomere variant repeat (TVR) duplication blocks.**

(A) Distribution of TVR duplication block size (bp). (B) Distribution of inter-block distances between paired TVR duplication blocks. (C) Pearson correlation between TVR duplication block size and inter-block distance. (D) Comparison of inter-block distances between smaller blocks (lower 50%) and larger blocks (upper 50%). Statistical significance was evaluated using a two-sided Wilcoxon rank-sum test.

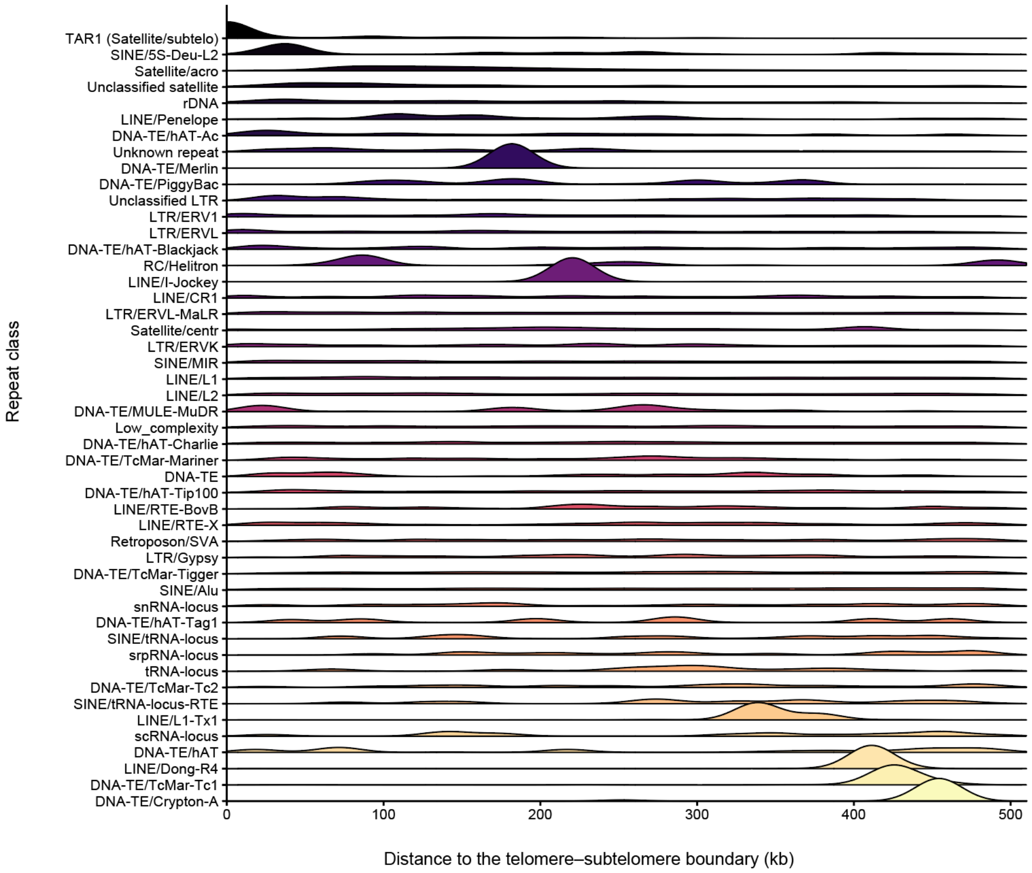


**Figure S28. Spatial distribution of subtelomeric repeat elements relative to the telomere–subtelomere boundary across the HPRC2 cohort.**

Distance distributions of major repeat element classes (annotated by RepeatMasker) across the 500-kb subtelomeric flanking regions.

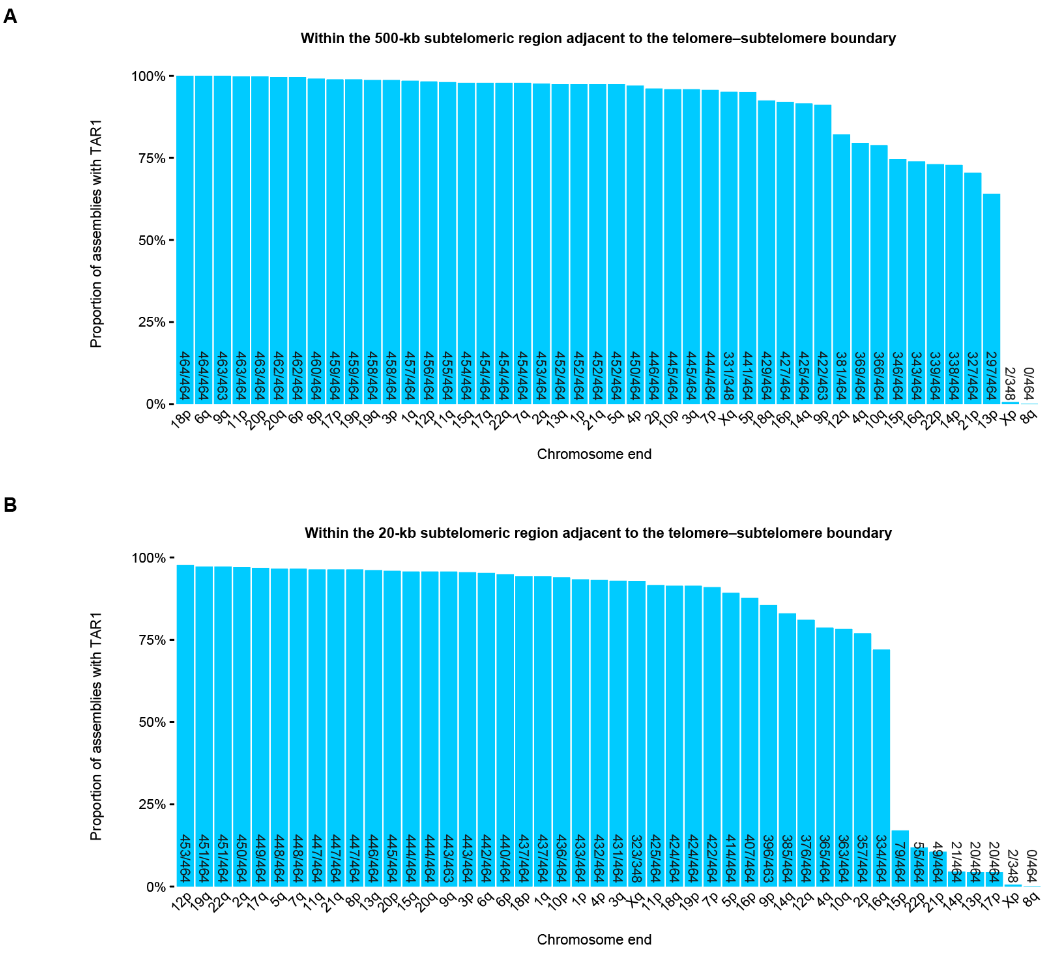


**Figure S29. Prevalence of TAR1 subtelomeric elements across the HPRC2 cohort.** (A, B) Frequency of TAR1 element occurrence at individual chromosome end across HPRC2 assemblies, with the exact number of occurrences further indicated. The calculations were performed within (A) 500-kb and (B) 20-kb subtelomeric regions adjacent to the telomere–subtelomere boundary respectively.

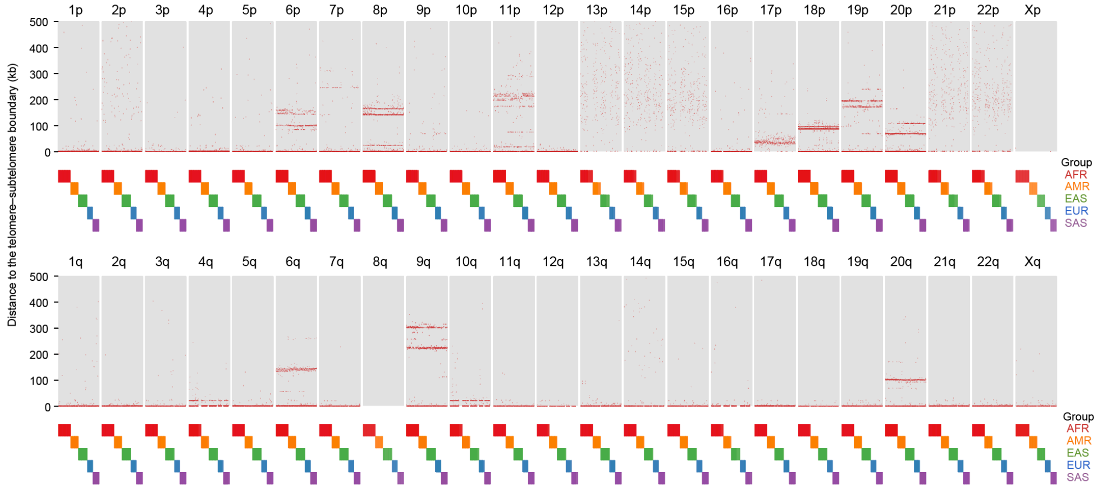


**Figure S30. Physical distribution of TAR1 elements across individual chromosome ends in the HPRC2 cohort.**

Haplotype-resolved presence patterns of TAR1 elements within the 500-kb subtelomeric regions of each chromosome end. Red dots denote assembly-specific TAR1 locations. Assemblies are stratified by continental group (AFR, African; AMR, Admixed American; EAS, East Asian; EUR, European; SAS, South Asian).

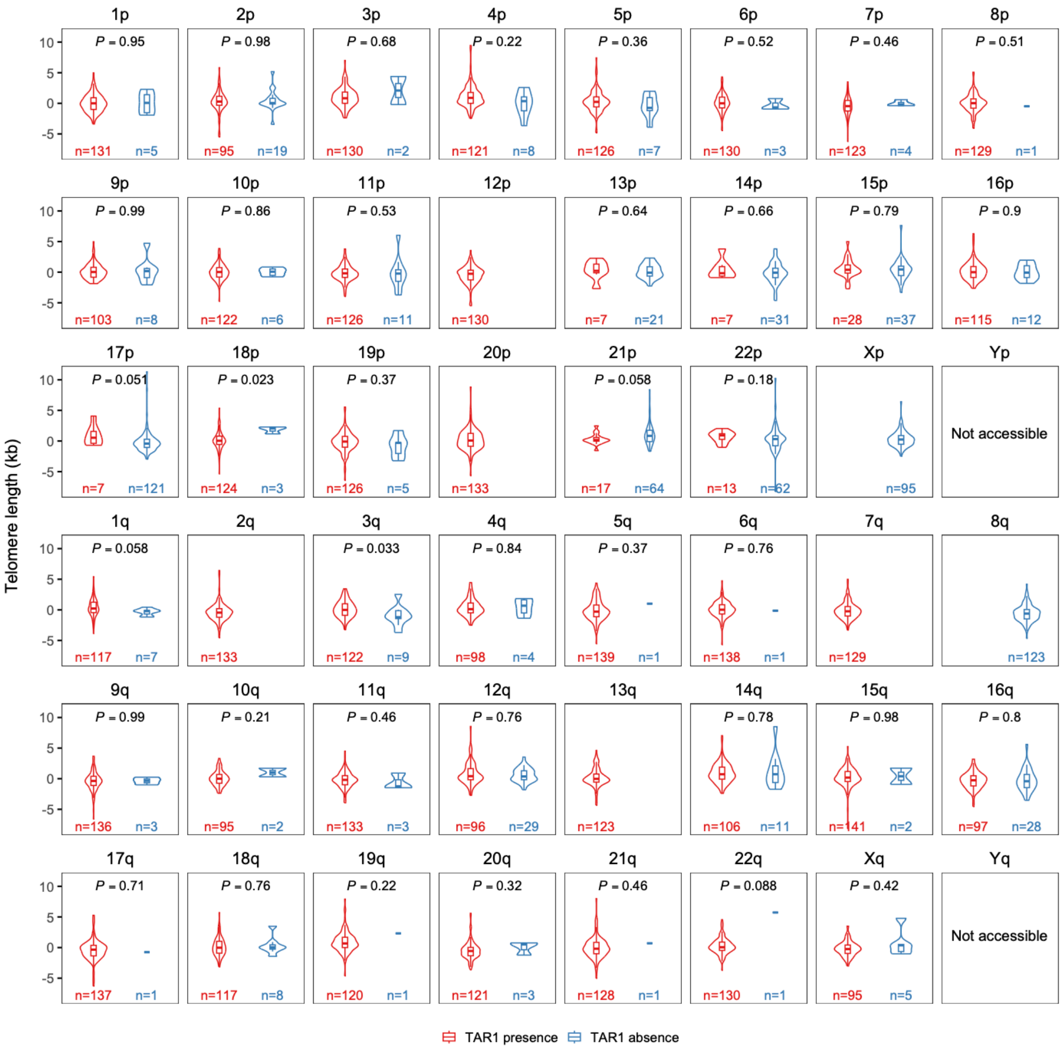


**Figure S31. Impact of subtelomeric TAR1 presence on chromosome-end-specific telomere length in the HPRC2 cohort.**

Comparison of telomere length at individual chromosome ends in the presence versus absence of subtelomeric TAR1 elements within the 20-kb subtelomeric regions adjacent to the telomere–subtelomere boundary. *P*-values were calculated using two-sided Wilcoxon rank-sum tests.

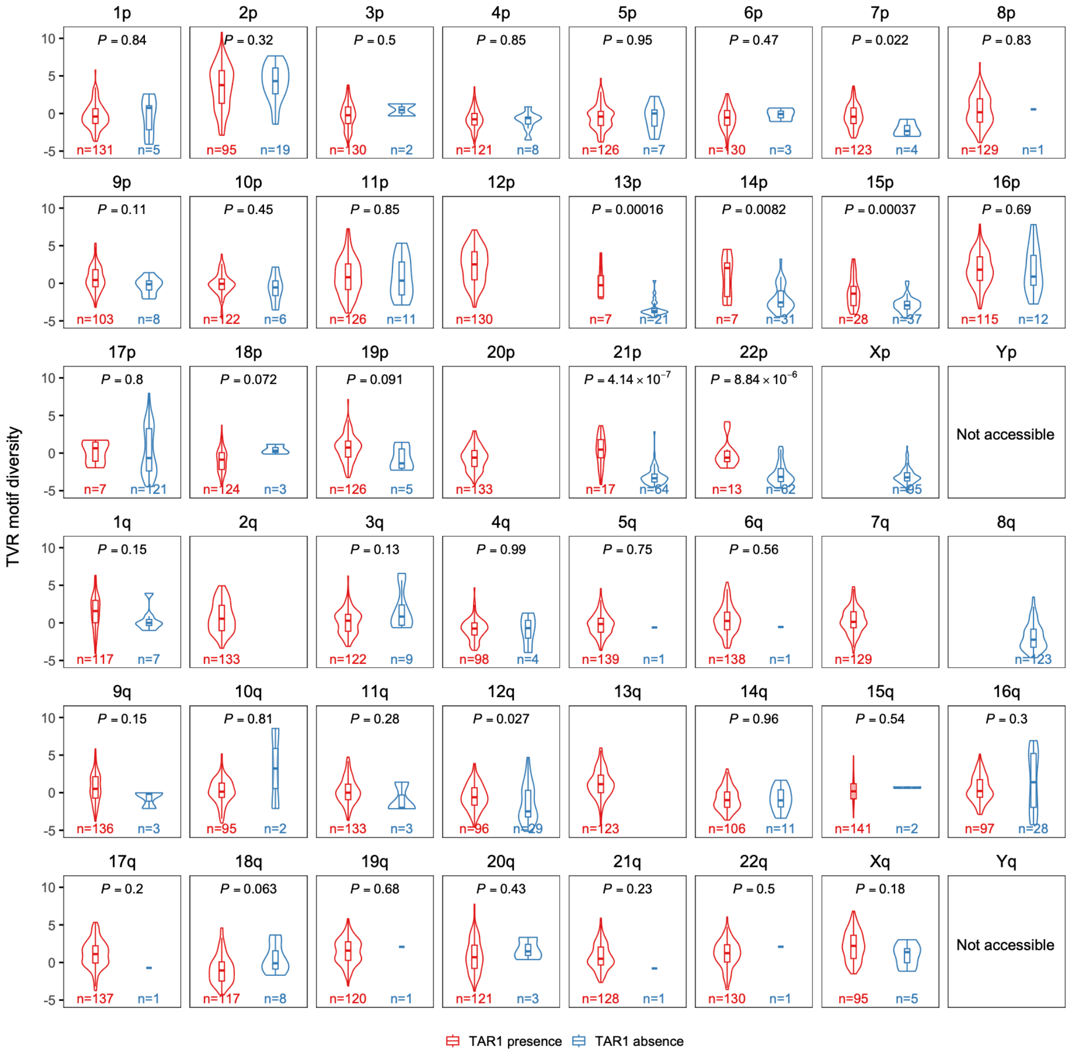


**Figure S32. Impact of subtelomeric TAR1 presence on chromosome-end-specific telomere variant repeat (TVR) motif diversity in the HPRC2 cohort.**

Comparison of TVR motif diversity at individual chromosome ends in the presence versus absence of subtelomeric TAR1 elements within the 20-kb subtelomeric regions adjacent to the telomere–subtelomere boundary. *P*-values were calculated using two-sided Wilcoxon rank-sum tests.

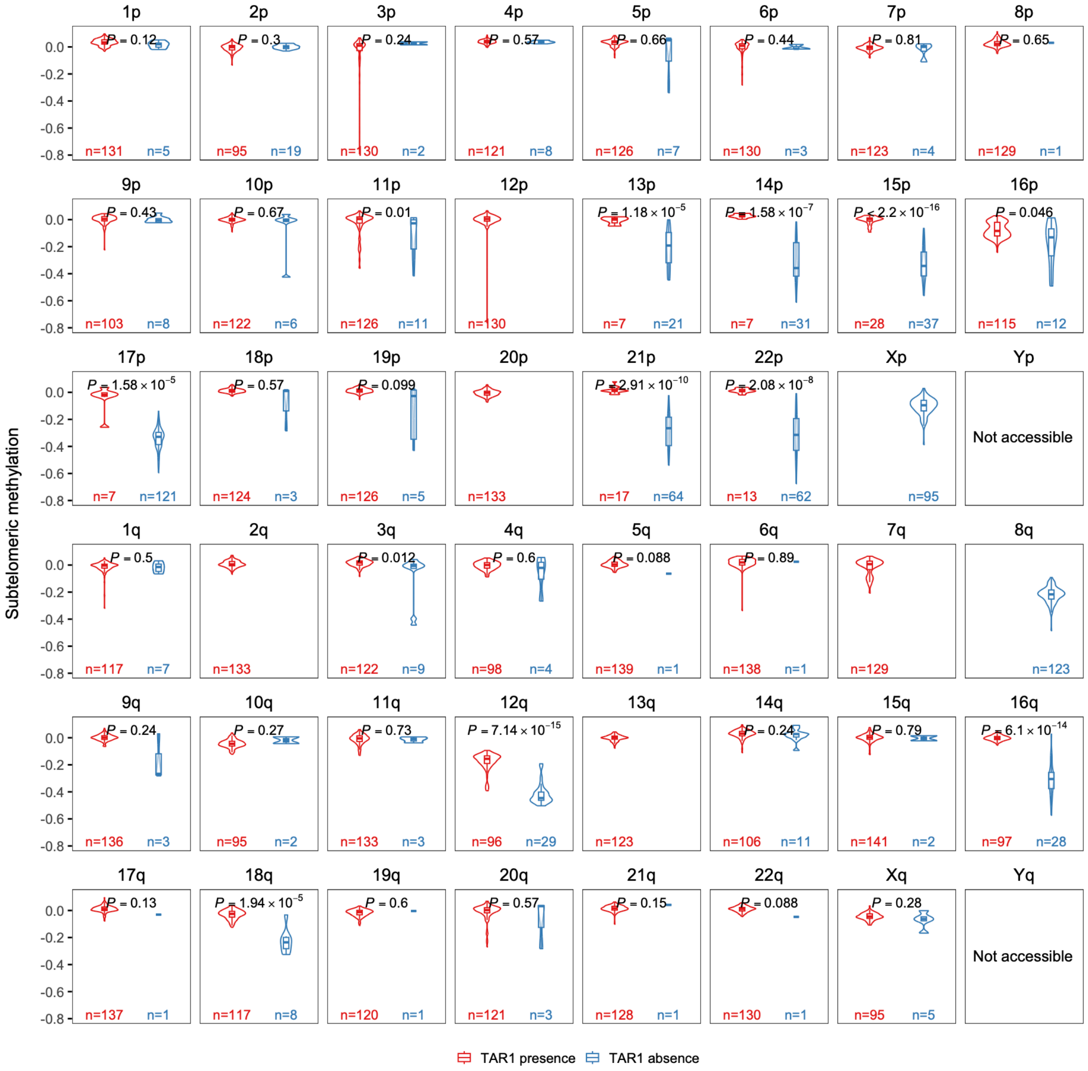


**Figure S33. Impact of subtelomeric TAR1 presence on chromosome-end-specific subtelomeric methylation levels in the HPRC2 cohort.**

Comparison of subtelomeric methylation levels at individual chromosome ends in the presence versus absence of subtelomeric TAR1 elements within the 20-kb subtelomeric region adjacent to the telomere–subtelomere boundary. *P*-values were calculated using two-sided Wilcoxon rank-sum tests.

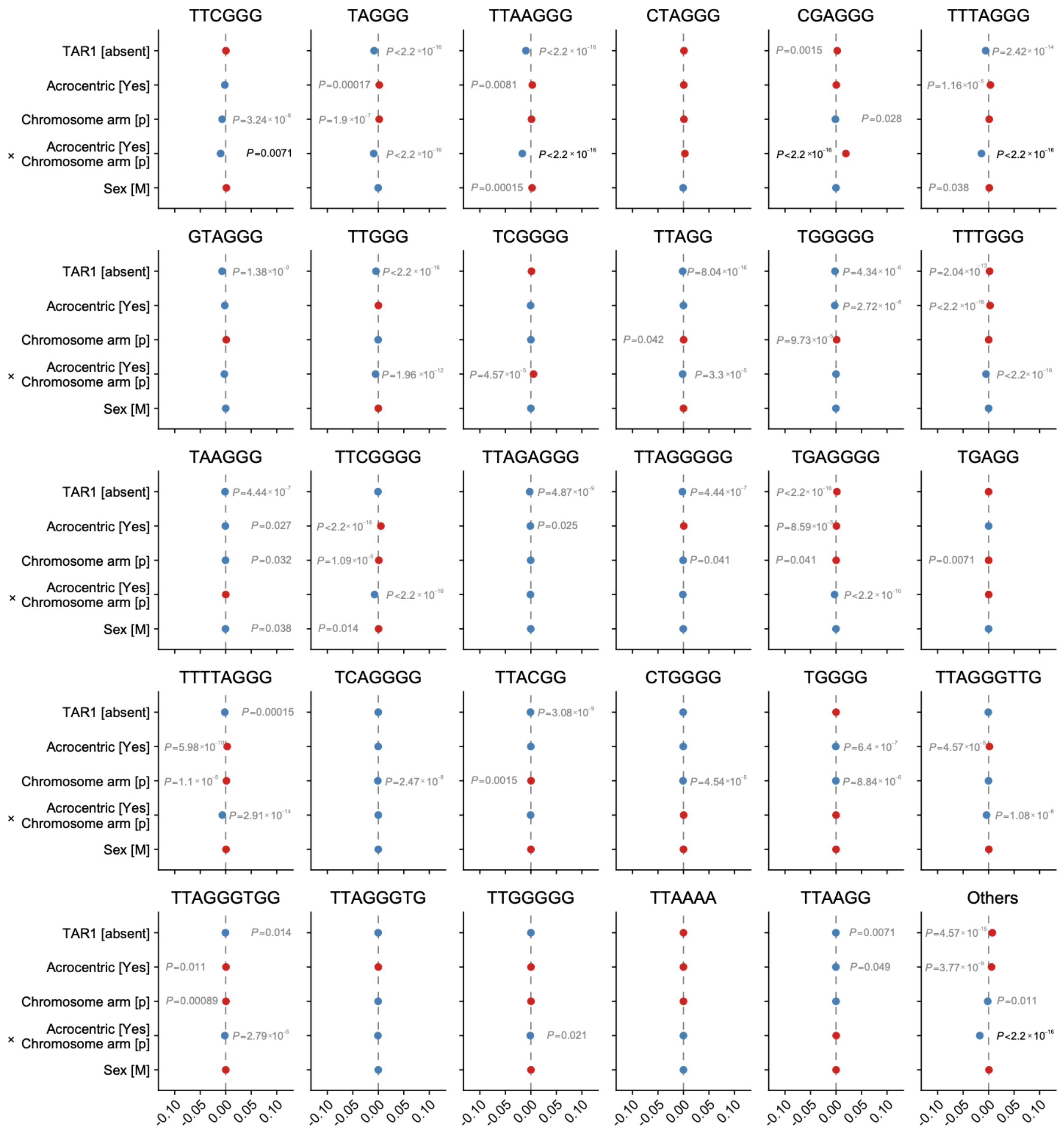


**Figure S34. Multivariable associations of subtelomeric and demographic features with telomere variant repeat composition in the HPRC2 cohort.**

This plot is complementary to Figure 7e. Forest plots show coefficient estimates and 95% confidence intervals from linear mixed-effects models fitted separately to the proportion of each displayed canonical or variant telomere repeat motif across the 73 ONT R10-sequenced HPRC2 individuals. Predictors included TAR1 status (presence/absence within 20-kb subtelomeric region adjacent to the telomere–subtelomere boundary), acrocentric chromosome-end classification, chromosome arm, gender and superpopulation. Reference categories were TAR1 present, non-acrocentric, p arm, female and African (AFR), respectively. Positive and negative coefficients are shown in red and blue; horizontal bars indicate 95% confidence intervals, and dashed vertical lines indicate no association (coefficient = 0). AMR, Admixed American; EAS, East Asian; EUR, European; SAS, South Asian.
